# Stress Changes the Material State of a Bacterial Biomolecular Condensate and Shifts its Function from mRNA Decay to Storage

**DOI:** 10.1101/2024.11.12.623272

**Authors:** Luis A. Ortiz-Rodríguez, Hadi Yassine, Ali Hatami, Vidhyadhar Nandana, Christopher A. Azaldegui, Jiayu Cheng, Yingxi Zhu, Jared M. Schrader, Julie S. Biteen

## Abstract

Bacterial ribonucleoprotein bodies (BR-bodies) are dynamic biomolecular condensates that play a pivotal role in RNA metabolism. We investigated how BR-bodies significantly influence mRNA fate by transitioning between liquid- and solid-like states in response to stress. With a combination of single-molecule and bulk fluorescence microscopy, biochemical assays, and quantitative analyses, we determine that BR-bodies promote efficient mRNA decay in a liquid-like condensate during exponential growth. On the other hand, BR-bodies are repurposed from sites of mRNA decay to reservoirs for mRNA storage under stress; a functional change that is enabled by their transition to a more rigid state, marked by reduced internal dynamics, increased molecular density, and prolonged residence time of ribonuclease E. Furthermore, we manipulated ATP levels and translation rates, and we conclude that the accumulation of ribosome-depleted mRNA is a key factor driving BR-body rigification, and that condensate maturation further contributes to this process. Upon nutrient replenishment, stationary-phase BR-bodies disassemble, releasing stored mRNAs for rapid translation, demonstrating that BR-body function is governed by a reversible mechanism for resource management. These findings reveal adaptive strategies by which bacteria regulate RNA metabolism through condensate-mediated control of mRNA decay and storage.

## Introduction

Biomolecular condensates are dynamic macromolecular assemblies that often form via phase separation,^1,2^ a reversible process that enables the cell to respond rapidly to environmental changes.^3^ These assemblies are selectively permeable and can exchange client molecules.^1,4^ Depending on their physical state, which can range from liquid droplets to hydrogels and insoluble aggregates, condensates facilitate various biological functions.^1,5^ Liquid-like condensates promote biochemical reactions by allowing internal diffusion and exchange with the dilute phase, while gel- and solid-like condensates have reduced diffusion and exchange, which slow down biochemical reactions and may potentially sequester biomolecules for storage.^1,2^ Notably, transitions from the liquid to solid state in biomolecular condensates have been linked to neurodegenerative protein aggregation disorders:^6,7^ reducing the fluidity of these condensates can negatively affect their function.^8,9^ However, in other cases, such as germ granules, solid-like states play important roles in mRNA storage and translational control in the developing embryo.^10^

Recently, biomolecular condensates have emerged as a broadly utilized mechanism for organizing diverse biochemical pathways within the bacterial cytoplasm,^11–19^ and this organizational paradigm is particularly important in bacteria because these organisms generally lack membrane-bound organelles.^20,21^ Bacterial ribonucleoprotein bodies (BR-bodies), the organelles that contain the RNA degradation machinery, were the first bacterial condensates demonstrated to assemble via liquid-liquid phase separation (LLPS), driven by interactions between the large intrinsically disordered region (IDR) of ribonuclease E (RNE; Fig. 1a) and RNA.^11^

**Fig. 1.**
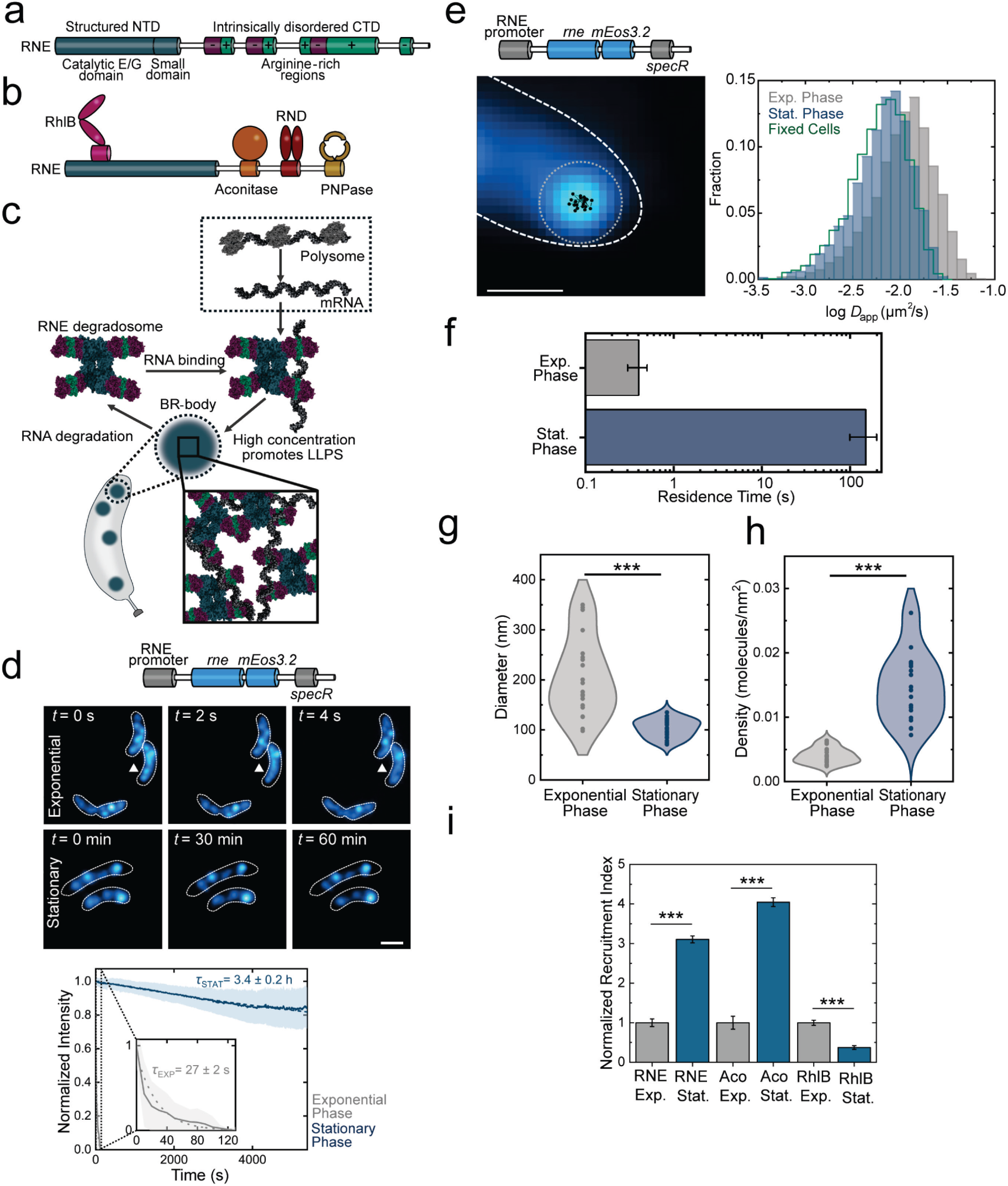
BR-bodies are dynamically arrested and compacted upon entering stationary phase. (a) Domain organization of *C. crescentus* RNase E (RNE). (b) The *C. crescentus* RNA degradosome includes RNE, RhlB, aconitase, RNase D, and PNPase. The RNE NTD has an RhlB binding site. The CTD has interaction sites for aconitase, RNase D, and PNPase. (c) In the working model for BR-body-mediated mRNA decay, untranslated mRNAs and the RNE degradosome can phase separate into a biomolecular condensate, thereby increasing their local concentration and accelerating decay processes within the BR-body. The condensate dissolves as mRNA is degraded, releasing degradosome components and oligonucleotides. (d) Lifespans of condensates based on an RNE-mEos3.2 construct (top). Middle: representative images of *C. crescentus* cells (white outlines) and RNE-mEos3.2 (blue) in each phase as a function of imaging time, *t*; arrowheads indicate representative RNE-mEos3.2 foci. Scale bar: 500 nm. Bottom: decay profiles of photobleaching-corrected intensities with estimates of the average condensate lifespans, *τ*_EXP_ and *τ*_STAT_, in exponential and stationary phases, respectively. Analyses include *N* = 427 foci in 211 cells (exponential) and *N* = 932 foci in 232 cells (stationary). (e-i) Dynamics of degradosome components within BR-bodies across growth phases. (e) Single-molecule tracking of RNE inside BR-bodies. Top-left: genetic construct used to track RNE. Bottom-left: a representative single-molecule trajectory of RNE-mEos3.2 within a BR-body (dotted circle) in a *C. cresentus* cell (dashed lines). The black dots indicate the super-localized positions of the molecule in consecutive 20-ms imaging frames. Scale bar: 500 nm. Right: Changes in the diffusivity of RNE within BR-bodies across growth phases. Number of trajectories analyzed: 5,001 (exponential phase) and 6,347 (stationary phase). The apparent diffusion coefficients for RNE-mEos3.2 trajectories in live cells are compared to the apparent diffusion coefficients of trajectories in fixed cells to distinguish between diffusing and static molecules. (f) Increase in residence time of RNE-mEos3.2 of BR-bodies between exponential and stationary phase. (g) Increase in BR-body diameter between exponential and stationary phases. The BR-body diameter is estimated as the maximum span of the cluster identified using density-based cluster analysis. Number of clusters analyzed: 20 per condition. (h) Increase in the density of RNE-mEos3.2 single-molecule localizations within BR-bodies during the transition from exponential to stationary phase. Each point shows the density in one condensate. Number of clusters analyzed: 20 per condition. (i) Normalized recruitment index of degradosome components RhlB-PAmChy, Aco-PAmChy, and RNE-mEos3.2 inside the BR-bodies in stationary phase relative to the recruitment in exponential phase, indicating the presence of incomplete degradosomes in BR-bodies during stationary phase. Error bars represent propagated standard deviations. All NRI values were normalized to the mean NRI of the exponential phase, setting the exponential phase values to 1. The error for each condition was calculated by propagating the original standard deviation using standard error propagation rules. Error bars in f and i: standard deviation from three biological replicates. *** in g and i denotes *p* < 0.001.

*Caulobacter crescentus* BR-bodies share functional similarities with eukaryotic processing bodies (P-bodies) and stress granules (SGs), as all three contain the mRNA decay machinery. RNE (Fig. 1a), the core scaffold and driver of BR-body function, is the essential and rate-limiting factor in bacterial mRNA decay.^22,23^ Similarly, Xrn1 is a core component of eukaryotic P-bodies and SGs, and it controls cytoplasmic mRNA decay.^24–26^ In addition, similar types of molecules are enriched in BR-bodies (Fig. 1b,c), P-bodies, and SGs, including DEAD-box RNA helicases, Lsm proteins, miRNAs, small-RNAs, and translationally repressed mRNAs.^27^ P-bodies and SGs have been predominantly studied under conditions of cell stress, and some studies indicate that P-bodies promote mRNA decay, while others show an mRNA storage function, whereas SGs have only been found to promote storage.^11,25,28^ These differences in the decay and storage functions are correlated with differences in the mobility of molecules within P-bodies^25,28–31^ and SGs,^32–34^ since P-bodies have been found to have higher mobility and more rapid exchange with the dilute phase than SGs. BR-bodies have been predominantly studied in exponentially growing bacteria cells, during which they appear to have liquid-like properties that promote mRNA decay (Fig. 1c).^11,35^ In contrast, under cell stress, BR-bodies undergo increased condensation that correlates with enhanced survival under these conditions, yet their internal dynamics and rates of mRNA degradation have not been previously explored under stress.^11^ Evidence from multiple bacterial species has shown that mRNA decay rates are reduced under conditions of nutrient limitation, such as when cells enter stationary phase, yet the role of BR-bodies in this slowdown of mRNA decay has not yet been explored.^36–41^

In this study, we investigate changes in the dynamics within BR-bodies between exponential growth and stationary phase and how these differences impact the mRNA decay pathway. We define the material state of a biomolecular condensate as the set of physical properties that determine internal mobility, exchange with the surrounding environment, condensate compaction, and thus biological function. Furthermore, in this work, a condensate is considered liquid-like when it exhibits dynamic internal rearrangement, rapid molecular exchange with the surrounding environment, lower biomolecular density, and short residence times of key components, while rigid condensates show reduced internal molecular diffusion, longer component residence times, increased compaction, and reduced exchange with the surrounding environment. Our live-cell single-molecule microscopy results demonstrate that BR-bodies transition from a dynamic, liquid-like state during exponential growth to a less dynamic, more rigid material state during stress. We demonstrate that this transition is associated with a functional shift from enhancing mRNA decay in exponential phase to promoting mRNA storage in the rigid state. In line with these results, we measure that the *in vitro* maturation of RNE condensates and subsequent arrest of dynamics lead to a strong reduction in RNE activity. Given the similarities between BR-bodies and eukaryotic biomolecular condensates like SGs and P-bodies, our findings further suggest that differences in cell growth conditions may also alter the material properties of P-bodies and SGs to shift the inherent preferences for mRNA decay vs. storage across diverse organisms.

### The transition from exponential to stationary phase is associated with BR-body rigidification

To investigate the dynamics of molecules within BR-bodies and their response to stress, we labeled the endogenous copy of RNE with the green-to-red photoconvertible fluorescent protein mEos3.2 (RNE-mEos3.2; Fig. 1d).^42^ This labeling strategy allows us to study the properties of BR-bodies as a whole using bulk fluorescence in the initial mEos3.2 green state, and to measure their internal dynamics at the single-molecule level in the mEos3.2 red state after photoconversion using 405-nm excitation. We compared BR-body dynamics during exponential and stationary growth phases to understand the effect of non-invasively induced cellular starvation, ensuring physiologically relevant observations of the stress response. We further sought to resolve two previous confounding observations: (1) there is an increased number of BR-bodies and a higher copy number of RNE in each condensate in cells at stationary phase (Extended Data Fig. 1a and Supplementary Figure 1),^11^ a nutrient-deprived state during which bacterial growth and division cease, and yet (2) mRNA decay is slower in stationary phase.

To investigate how stationary phase alters BR-bodies, we performed time-lapse imaging of cells in exponential and stationary phases. In exponential phase, BR-bodies rapidly assemble and disassemble, consistent with their known role in mRNA decay (Fig. 1d, Supplementary Movie 1). However, in stationary phase, BR-bodies are dramatically stabilized (450-fold; Fig. 1d, Supplementary Movie 2). As RNA is required for BR-body phase separation,^11^ the decreased dissolution rate of BR-bodies during stationary phase suggests reduced RNA decay activity, as mRNA degradation is necessary for BR-body disassembly (Fig. 1c).^11^ Single-molecule tracking of RNE-mEos3.2 and the degradosome protein aconitase-PAmChy, previously identified as a direct binding partner to RNE^11,43,44^ and core component of the *C. crescentus* RNA degradosome, revealed measurably reduced diffusivity of RNE and its degradosome client proteins within BR-bodies during stationary phase (Fig. 1e and Extended Data Fig. 1b, c), consistent with a more rigid internal microenvironment. In addition, we compared RNE-mEos3.2 diffusion in both growth phases with the apparent diffusion of immobile RNE-mEos3.2 molecules in fixed cells. While we measured slow diffusion of RNE-mEos3.2 within BR-bodies during exponential phase, this value is higher than our lower bound as determined in fixed cells. In contrast, the diffusion profile of RNE-mEos3.2 in stationary phase cells closely matches that of fixed cells, suggesting that, under our experimental conditions, RNE-mEos3.2 is essentially static within the BR-bodies during stationary phase. Furthermore, the residence time of RNE within BR-bodies is 370-fold longer in stationary phase compared to exponential phase (Fig. 1f). After identifying the RNE-mEos3.2 clusters with a density-based cluster analysis,^45^ we determined the molecular density and estimated the BR-body diameter, *d̄*, as the maximum span of each cluster (Fig. 1g,h). In exponential phase, the BR-bodies are less dense and larger (*d̄* = 206 ± 2 nm) than in stationary phase (*d̄* = 102 ± 1 nm) (mean ± SEM; *p* < 0.001). Importantly, the decreased diffusivity and increased residence time of RNE within the BR-bodies, along with their compaction, are also observed under ethanol stress, indicating that dynamic arrest behavior likely occurs under other inducing stresses (Extended Data Figs. 1d‒f,2). This compaction explains the decreased internal dynamics, where the rigid state constrains RNE within the BR-body.

We also analyzed the diffusion of RNE outside the BR-bodies in both exponential and stationary phase cells to determine whether the observed changes in RNE mobility were specific to the condensate internal microenvironment. The mobility of RNE molecules not associated with the BR-bodies remained unchanged between growth phases (Extended Data Fig. 1c), indicating that the reduced mobility observed within BR-bodies during stationary phases arises from changes in the condensate internal microenvironment rather than reflecting overall cellular changes between growth phases. This lack of a change in diffusion outside the BR-bodies reflects the short timescales probed by our single-molecule trajectories (∼160 ms), which primarily report on local motions and therefore do not capture slower cytoplasmic dynamics.^46^ In contrast, the microenvironment within stationary phase BR-bodies is sufficiently dense that a measurable reduction in RNE-mEos3.2 mobility is evident even on these short timescales, while condensate-level diffusion becomes distinct at longer observation windows (Supplementary Figure 2).

To determine whether BR-bodies retain characteristics of phase-separated condensates during stationary phase, rather than forming irreversible aggregates, we treated cells with 5% 1,6-hexanediol (HD), a commonly used aliphatic alcohol that disrupts the weak hydrophobic interactions essential for phase separation.^47–49^ BR-bodies dissolve within ∼ 25 minutes of HD treatment (Extended Data Fig. 1g), confirming that they remain sensitive to HD despite rigidification during stationary phase. This finding suggests that BR-bodies retain hallmark features of biomolecular condensates, even in the rigid state, and that the observed rigidification reflects a dynamic arrest rather than a transition to an aggregated or misfolded state.

BR-bodies concentrate the RNA degradosome complex, which in *C. crescentus* contains aconitase, RhlB, and PNPase (Fig. 1b).^11,44^ We quantified the recruitment of aconitase and RhlB to BR-bodies during exponential and stationary phases, normalizing by differences in overall expression levels. After normalization, we found that aconitase recruitment increases 4-fold, while RhlB recruitment decreases 3-fold in stationary phase (Fig. 1i). These results reveal a stoichiometric reorganization of BR-bodies under stress. In particular, the concomitant loss of RhlB and enrichment of RNE and aconitase indicate the presence of incomplete degradosomes inside BR-bodies during stationary phase. DEAD-box RNA helicases are central regulators of biomolecular condensates;^50^ their ATPase cycle maintains the condensates in a dynamic, non-equilibrium state, facilitating the enrichment of client molecules and buffering the dispersed ribonucleoprotein pool. The decreased recruitment of the DEAD-box RNA helicase RhlB thus likely impairs the internal remodeling of BR-bodies, potentially contributing to their rigidification during stress.

### Increased levels of poorly translated mRNA and maturation trigger the transition to the rigid material state

During stationary phase, bacterial cells encounter nutrient scarcity, which slows down cellular processes including metabolic activity, leading to lower ATP levels (Fig. 2a).^51^ Reduced ATP levels subsequently impact various processes, including protein synthesis and translation, promote phase separation of proteins with intrinsically disordered regions (IDRs),^18^ contribute to protein aggregation,^12,13^ and cause physical changes in cytoplasm,^46,52^ creating a glassy state that slows the diffusion of mesoscale objects such as BR-bodies (Supplementary Figure 2). Additionally, since DEAD-box helicases like RhlB require ATP hydrolysis to remodel, dissolve, cooperatively bind RNA,^53^ prevent strong interactions, and maintain the fluidity of the condensates,^50^ the low ATP levels at stationary phase may impair these activities.

**Fig. 2.**
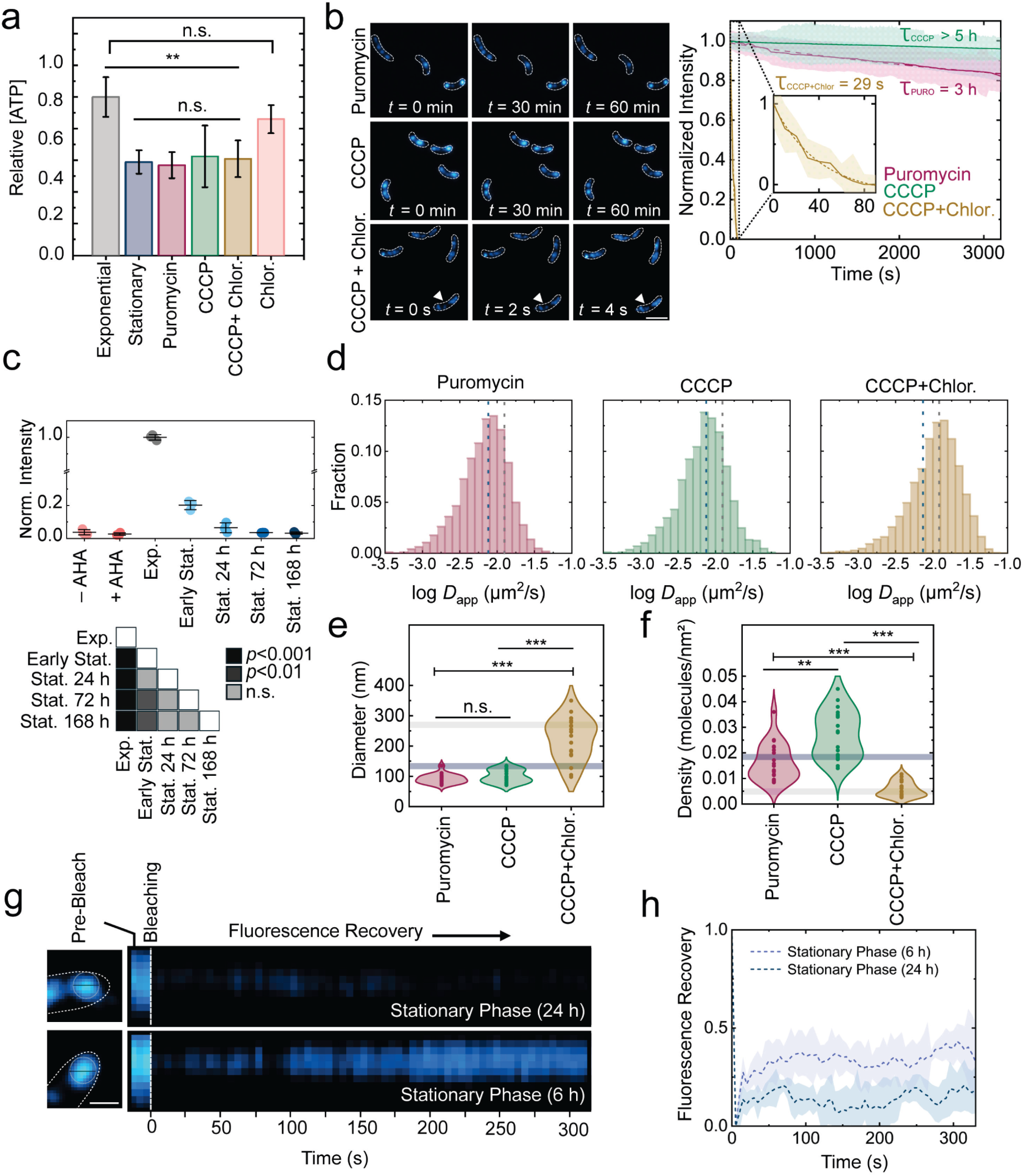
Rigidification of BR-bodies is promoted by increased levels of poorly translated mRNAs. (a) Quantification of intracellular ATP concentrations in *C. crescentus* using a luciferase-based assay following no treatment or treatment as described. ATP levels are normalized to exponential growth levels. (b) Lifespans of BR-bodies in cells under the acute treatment conditions described in (a). Left: Representative images of RNE-mEos3.2 (blue) in *C. crescentus* cells (white outlines) as a function of treatment and imaging time, *t*. Arrowheads indicate representative RNE-mEos3.2 foci. Scale bar: 2 µm. Right: Decay profiles of photobleaching-corrected intensities used to estimate the average condensate lifespan. Analyses include *N* = 308, 347, and 241 foci for puromycin, CCCP, and CCCP + chloramphenicol, respectively, measured across 88, 193, and 301 cells, respectively. (c) Top: relative translational levels in exponential and stationary phases using Biorthogonal Non-Canonical Amino Acid Tagging (BONCAT), and controls with azidohomoalanine (AHA) and chloramphenicol and without AHA. Error bars: standard deviation of three replicates. Bottom: statistical analysis. (d) Impact of acute treatments on the diffusivity of the BR-body RNE scaffold. The distribution of apparent diffusion coefficients for RNE-mEos3.2 trajectories is measured following each treatment. Vertical dashed lines: distribution medians. Number of trajectories analyzed: 2,126 (puromycin), 1,637 (CCCP), and 1,441 (CCCP + chloramphenicol). (e) Effect of acute treatments on the BR-body diameter, which is estimated as the maximum span of the cluster identified using density-based cluster analysis following each treatment. Horizontal lines indicate the mean of the distribution of exponential phase and stationary phase BR-body diameters. Number of clusters analyzed: 20 per condition. (f) Changes in the density of RNE-mEos3.2 single-molecule localizations within BR-bodies following each treatment. Horizontal lines indicate the mean of the distribution of exponential phase and stationary phase BR-body densities. Number of clusters analyzed: 20 per condition. (g) Dynamics inside BR-bodies in cells during prolonged stationary phase. Representative fluorescence recovery after photobleaching (FRAP) experiment kymographs during early (6 h) and late (24 h) stationary phase cells showing the bleaching and recovery of the RNE-mEos3.2 foci (blue) as a function of time, *t*. Scale bar: 500 nm. (h) FRAP of RNE-mEos3.2 in cells at the early and late stationary phase. *N* = 20 foci. Error bars in a and c: standard deviation from three biological replicates. Statistics in a, e, and f: *** denotes *p* < 0.001, ** denotes *p* < 0.005, and n.s. denotes *p* > 0.05.

To separate the effects of low ATP levels and reduced translation on BR-body rigidification, we artificially decreased ATP and translation levels in exponential phase cultures of *C. crescentus*. To lower ATP levels, we treated the cultures with the protonophore carbonyl cyanide *m*-chlorophenyl hydrazone (CCCP) to disrupt the proton motive force,^54^ rapidly inhibiting ATP synthase. A 20-min treatment with 100 μM CCCP was found to reduce cellular ATP concentrations to levels comparable to those observed during stationary phase (Fig. 2a). To reduce translation levels, we used puromycin (150 μg/mL for 30 min), which prematurely cleaves peptide chains^55^ and releases ribosome subunits and mRNA, thereby increasing the availability of ribosome-depleted transcripts in the cytoplasm. Since puromycin treatment also results in ATP levels similar to those observed in stationary phase (Fig. 2a), we treated cells with CCCP and the translation elongation inhibitor chloramphenicol as a control. Chloramphenicol prevents translation by blocking peptide bond formation and accumulating ribosomes on mRNA.^56^ Interestingly, treating exponential phase cultures with CCCP and puromycin leads to long-lived BR-bodies (Fig. 2b), similar to those observed during stationary phase (Fig. 1d). However, when cells are treated with the combination of CCCP and chloramphenicol, BR-bodies remain short-lived, similar to those observed during exponential phase in the absence of stress (Fig. 2b). Similarly, BR-bodies remain short-lived when cells are treated with chloramphenicol only (Supplementary Figure 3). This finding suggests that the increased lifespan of BR-bodies upon treatment with CCCP might be an indirect effect of dropping ATP levels inhibiting translation rather than directly related to ATP availability. Translation is typically the most energy-intensive process during exponential growth,^57,58^ and ATP and GTP are essential for several steps in translation, including the esterification of amino acids to tRNAs, selection of the aa-tRNAs to the ribosome, the assembly of ribosomal subunits, and the translocation of ribosomes along the mRNA. When NTP levels are decreased, these processes slow, resulting in reduced translation rates. Additionally, biorthogonal non-canonical amino acid tagging (BONCAT) revealed a significant decrease in translation levels during the late stationary phase, with reductions of 30-fold, as well as in cells treated with puromycin, CCCP, and the combination of CCCP and chloramphenicol, all relative to exponential phase (Fig. 2c, Extended Data Fig. 3).

Similar to the extended lifespan of BR-bodies observed upon treatment with CCCP and puromycin, these treatments also replicated stationary-phase decreases in RNE-mEos3.2 diffusivity inside the BR-bodies (Fig. 2d, Supplementary Figure 4a), BR-body size (Fig. 2e, Supplementary Figure 4b), RNE molecular density (Fig. 2f, Supplementary Figure 4c), and residence time within BR-bodies (Extended Data Fig. 2). Conversely, cells treated with a combination of CCCP and chloramphenicol do not rigidify. This finding suggests that the rigid material state transition in stationary phase requires increased amounts of ribosome-free mRNA transcripts in the cytoplasm, whereas ATP scarcity alone is insufficient.

We identified low ribosome occupancy as a key factor contributing to the rigid material state of BR-bodies during stationary phase. However, under prolonged stress, biomolecular condensates have been observed to mature from having liquid-like to solid-like properties.^1,59–63^ Initially, their components interact transiently: molecules freely rearrange and exchange with the surrounding phase, resembling liquid behavior. Over time, mechanisms such as entanglement of disordered polypeptides,^64^ vitrification,^65^ percolation,^66^ and nucleation and elongation of amyloid-like fibers^67^ contribute to rigidification and decreased molecular dynamics.^68^ We investigated the role of maturation in BR-body rigidification by *in vivo* fluorescence recovery after photobleaching (FRAP) of BR-bodies (RNE-mEos3.2) in cultures of *C. crescentus* cells at 6 hrs and 24 hrs into stationary phase (Fig. 2g,h). The condensates become less dynamic over time in stationary phase. The BR-body fluorescence recovers only minimally at both time points, consistent with a rigid material state, but BR-bodies are slightly more dynamic at 6 hrs than at 24 hrs. To further investigate the role of maturation, we reconstituted the system *in vitro* and extracted meaningful information based on RNE, the first bacterial protein found to undergo LLPS at physiological concentrations without crowding agents.^11^ FRAP performed on RNE droplets reconstituted *in vitro* and aged with RNA for 30 min, 4 hrs, and 8 hrs (Extended Data Fig. 4) revealed that condensates aged for 30 min were significantly more dynamic, with nearly full fluorescence recovery within 2.5 min, whereas condensates aged for 4 hrs exhibited minimal recovery and 8 hrs were even less dynamic. These results support the role of maturation in changing the material state of BR-bodies.

### The material state transition of BR-bodies during stress shifts their function from promoting mRNA decay to facilitating mRNA storage

Our *in vivo* and *in vitro* results establish that, during stress conditions, the internal dynamics and structure of BR-bodies become more rigid. Previous two-color images of mRNA and RNE using MS2 RNA labeling^69^ and fluorescence *in situ* hybridization (FISH) during exponential growth have shown that transcripts such as *rsaA* exhibit low colocalization (∼28%) with RNE in BR-bodies in conditions allowing active RNA degradation.^35^ However, colocalization increases significantly (∼80%) when RNE endonuclease cleavage is blocked using an RNE active site mutation (ASM) involving the two residues coordinating the catalytic Mg^2+^ ion in the active site^35^, which, taken together, suggests mRNA decay within BR-bodies during exponential growth. These results suggest that the transition to a more rigid material state, along with its associated effects, may enable BR-bodies to store mRNA during stress conditions.

To investigate the role of BR-bodies in mRNA storage, we performed two-color imaging of RNE and mRNA using a live-cell MS2 system and FISH, and we measured mRNA degradation rates using quantitative reverse transcription polymerase chain reaction (qRT-PCR). Notably, this study employs an MS2 system optimized to accurately track short-lived mRNAs.^70^ We selected four key mRNA transcripts needed for *C. crescentus* exponential growth: *rsaA*, *rne*, *spmX*, and *ctrA*. All four mRNAs were previously found to be stabilized in an RNE-IDR truncation mutant incapable of making BR-bodies, suggesting BR-bodies play a role in their degradation.^35,71^ The RsaA S-layer protein is the most abundant protein in *C. crescentus*^72^. RsaA was chosen for comparison with the prior result, where RNE cleavage is blocked using the ASM, providing insight into the role of the catalytic activity of RNE in the condensate and its effect on transcript recruitment into BR-bodies.^35^ RNE is crucial in RNA processing, degradation,^22,23^ BR-body formation, and cell fitness.^11^ The integral membrane protein SpmX enables cell division by localizing the histidine kinase DivJ to the stalked pole.^73,74^ Finally, CtrA is a master regulator of the cell cycle; it controls chromosome replication and transcription, and it modulates the expression of about 25% of cell cycle-regulated genes in *C. crescentus*.^75–79^ These strategically selected transcripts enabled us to thoroughly assess the role of BR-bodies in mRNA storage under stress and to determine the relevance of the stored transcripts.

Images of MS2-labeled *ctrA*, *spmX*, and *rsaA* transcripts (Fig. 3a,b, and Extended Data Fig. 5) highlight their higher recruitment to BR-bodies during stationary phase compared to exponential phase; these data sets were quantified using Pearson correlation coefficients (PCC). All transcripts colocalize more with BR-bodies during stationary phase, similar to the behavior of *rsaA* when RNE cleavage is blocked, suggesting significantly lower degradation activity by RNE during this phase. This mRNA RNE colocalization is further confirmed by FISH (Supplementary Figure 5). To ensure that the observed differences in BR-body colocalization across growth phases reflect transcript-specific recruitment dynamics rather than analytical artifacts or changes in overall transcript abundance, we used transfer-messenger mRNA (tmRNA) as a negative control. tmRNA is abundant in *C. crescentus,* and is strongly depleted from the BR-bodies with a log_2_ fold-change of −2.8.^35^ Upon applying the same colocalization analysis to the target transcripts, we found that tmRNA colocalizes significantly less with BR-bodies (PCC = 0.13 ± 0.04) compared to all other transcripts, even during exponential phase (Supplementary Figure 6). These results confirm that the high levels of colocalization observed for specific transcripts during stationary phase are not simply due to increased transcript abundance; rather, they reflect selective recruitment into BR-bodies under stress.

**Fig. 3.**
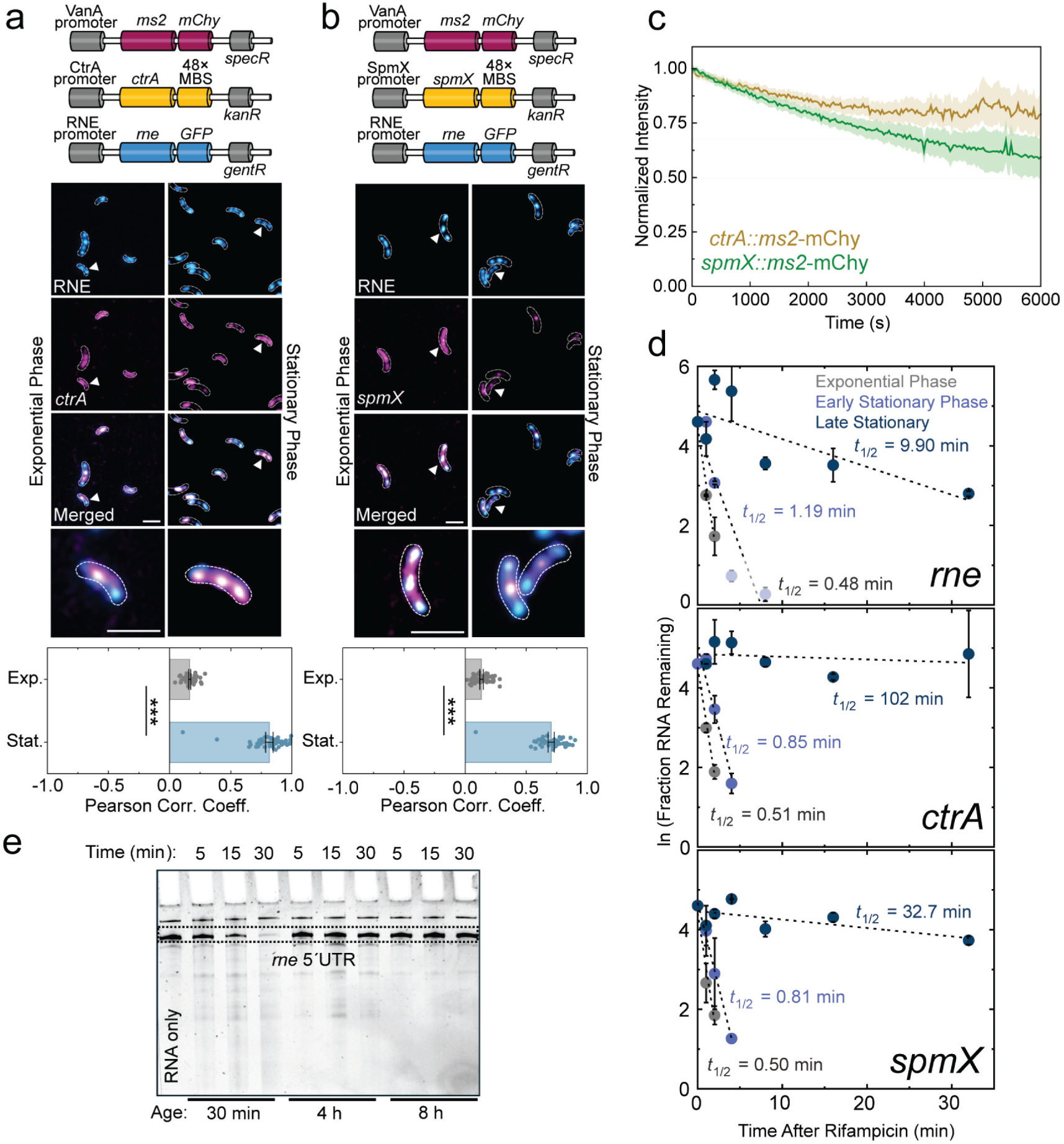
Rigidification of BR-bodies switches their function from mRNA decay to storage of key growth mRNAs. (a,b) Stability of the *ctrA* (a) and *spmX* (b) transcripts in BR-bodies. Top: genetic constructs used to observe the transcript stability within the BR-bodies *in vivo*. Middle: Representative two-color images showing higher recruitment of the *ctrA* and *spmX* transcripts (magenta) to BR-bodies (blue) during stationary phase (left) relative to exponential phase (right). The merged channel is shown on two magnification levels for clarity. Bottom: Differences in recruitment quantified by the Pearson correlation coefficients between channels. Circles: PCC values for each cell. Error bars: SEM from 50 cells analyzed per phase. *** denotes *p* < 0.001. Arrowheads indicate representative foci. Scale bars in (a) and (b): 2 µm. (c) Decay profiles of *ctrA* and *spmX* in the late stationary growth phase. Analyses include *N* = 108 and 174 foci for the *ctrA* and *spmX* transcripts, respectively, measured across 28 and 49 cells, respectively. (d) qRT-PCR measurements of RNA half-lives for *rne*, *ctrA*, and *spmX* transcripts highlight the changes in transcript stability from exponential phase to the late stationary phase. Error bars: standard deviation from three biological replicates. (e) Condensate aging leads to decreased RNE cutting activity *in vitro*, as evidenced by the *rne* mRNA band in a 7% urea TBE/PAGE gel.

We also assessed the stability of these transcripts tagged with MS2-RNA hairpins by analyzing the photobleaching-corrected intensities of MS2-mChy foci colocalized with RNE foci during both growth phases (Fig. 3c and Extended Data Fig. 5). To ensure that our observations accurately reflect transcript stabilization and not artifacts of the labeling system, we employed two versions of the MS2 system which are both utilized with a MS2-mChy dimerization deficient variant:^80^ a 48×-array with weaker MS2 coat protein RNA hairpin affinity^70^ and a 96×-array.^35^ Previous studies have demonstrated that binding of the coat protein to MS2 binding sites artifactually stabilizes mRNA.^81–83^ Therefore, we confirmed that any observed accumulation of the mChy signal within BR-bodies was due to real transcript stabilization and not an effect of the labeling system. As shown in Fig. 3a,b, mChy foci do not colocalize with BR-bodies during exponential phase when using the 48×-array system. However, using the 96×-array version (Extended Data Fig. 5), we observed mChy foci colocalizing with BR-bodies, even during exponential phase. Despite the potential transcript stabilization caused by the more perturbing 96×-array MS2 system, we still observed significant differences in the decay profiles between exponential and stationary phases. During exponential phase, MS2-mChy (96×-array) foci disassemble in under a few min, consistent with rapid internal decay of mRNAs (Extended Data Fig. 5). In contrast, during stationary phase, MS2-mChy (48×-array) foci and stably colocalize with BR-bodies for significantly longer, indicating slowed mRNA decay (Fig. 3c and Extended Data Fig. 5). To investigate the observations of the MS2-tagged RNAs, we measured the RNA decay rates. mRNA half-life measurements by qRT-PCR confirmed this stabilization: *rne*, *spmX*, and *ctrA* exhibit 20-, 65-, and 200-fold stability increases, respectively, from exponential to late stationary phase (Fig. 3d), and the stabilization was confirmed by a second method, mRNA FISH, for *ctrA* mRNA and *spmX* mRNA (Supplementary Figure 7). To test whether stabilization in stationary phase depends on BR-body formation, we measured the stability of *spmX* and *ctrA* transcripts in stationary phase cultures of *C. crescentus* expressing RNE-ΔCTD; these cells cannot assemble BR-bodies.^11^ In this background, transcript stabilization was reduced to only ∼3-fold, confirming that BR-bodies are essential for the pronounced stabilization of mRNA during stress (Supplementary Figure 7). Together, these measurements suggest that the rigid state of RNE is associated with a dramatic slowdown in mRNA decay activity, consistent with a model in which BR-body material state regulates RNA turnover.

Hfq is a well-known regulator of RNA stability, and in *E. coli*, Hfq was observed to form condensates with RNA under conditions of nutrient deprivation, where it was also observed to colocalize with *E. coli* RNE.^16,84,85^ In *C. crescentus*, Hfq was observed to colocalize to a subpopulation of BR-bodies,^43^ suggesting it might impact the stability of transcripts in rigid BR-bodies observed in stationary phase. To test whether the RNA chaperone Hfq, a BR-body constituent,^43^ contributes to transcript stabilization, we measured *ctrA* mRNA decay in a *Δhfq* background. In exponentially growing *Δhfq* cells, *ctrA* mRNA decays slightly more slowly than in wild-type cells, but still degrades rapidly (Supplementary Table 1), indicating that Hfq is not required for rapid turnover in exponential phase. In stationary phase, *ctrA* mRNA remains highly stable in *Δhfq* cells, similarly to wild type, suggesting that Hfq is not required for BR-body-mediated transcript stabilization. Consistently, the diffusion profile of RNE-mEos3.2 within BR-bodies remains unchanged in the *Δhfq* background (Supplementary Figure 8), indicating that the change to the rigid material state does not depend on Hfq. Hence, the stabilization during stationary phase is independent of Hfq and instead reflects protection conferred by the BR-body environment itself. As an additional control, we measured the stability of tmRNA (Supplementary Table 1), a highly abundant transcript that is not enriched in the BR-bodies. tmRNA remained fully stable in both exponential and stationary phases (> 8 min and > 16 min half-lives, respectively), confirming that transcripts not recruited into BR-bodies do not exhibit phase-dependent changes in decay rate. This finding further supports the model that BR-body-mediated stabilization depends on selective mRNA recruitment.

To investigate whether the reduced RNA cleavage activity of RNE is a direct result of rigidification, we assessed the cutting activity of RNE in 30-min, 4-hr, and 8-hr *in vitro*-reconstituted condensates (Fig. 3e). These aged RNE droplets were formed with RNE and RNA, but without Mg^2+^, an essential cofactor for RNA cleavage by RNE. At each droplet age, RNA cleavage was initiated by adding Mg^2+^, and samples were collected at various time points to assay RNA cleavage.^71^ RNE efficiently cleaves the *rne* 5′ UTR in droplets aged for only 30 min,^71^ with the corresponding band in the PAGE gel disappearing within 30 min of Mg^2+^ addition (Fig. 3e). In contrast, virtually no RNA cleavage was observed in the rigidified droplets at the 4-hr and 8-hr timepoints. This finding suggests that maturation into the rigid state can directly shut off RNE cleavage. Furthermore, we hypothesize that BR-body formation during stress serves as a repository for mRNA and a means to sequester RNE and potentially other degradative enzymes in an inactive state. This hypothesis is supported by the higher amount of RNE in condensates during stationary phase, and by our recent work, where we demonstrated that overexpression of RNA during exponential phase leads to growth arrest. Taken together with our *in vitro* results, these results support a model in which BR-body rigidification reduces RNE activity, allowing for selective preservation of mRNAs during stress.

### Fluidized BR-bodies are capable of the functional release of mRNAs

Stored mRNAs may be useful for growth-arrested cells either by acting as a nutrient source to promote regrowth or by being released to ribosomes to restart the translation program upon the addition of new nutrients. To determine if mRNA release and subsequent translation can occur after cells exit stationary phase and enter exponential growth, we assessed whether rigidification is reversible by observing if stationary-state BR-bodies fluidize over time after nutrient replenishment (Extended Data Fig. 6). Indeed, BR-body disassembly was detected after 90 min of nutrient replenishment. We further investigated transcript storage by BR-bodies post-nutrient replenishment by monitoring the colocalization of the *spmX* transcript with BR-bodies in stationary phase cultures (Supplementary Figure 9). After 3 hrs, the colocalization profile of *spmX* with BR-bodies returns to that observed during exponential phase (i.e., PCC = 0.22 ± 0.05 vs. 0.25 ± 0.04 in Fig. 3b), suggesting that stored mRNA exits BR-bodies upon removal of stress.

Additionally, to evaluate whether these transcripts could be translated after exiting BR-bodies, we performed a series of experiments designed to distinguish between translation of pre-existing transcripts and translation of newly transcribed mRNAs. In wild-type *C. crescentus*, nutrient replenishment after stationary phase was examined under conditions where new RNA synthesis was blocked with rifampicin. RNE-mEos3.2 fluorescence recovered substantially over 6 hours following whole-cell bleaching (Fig. 4a,b), indicating that recovery does not require transcription of new mRNAs and must arise from mRNAs that existed prior to rifampicin treatment. To confirm that this recovery reflects active translation rather than delayed protein maturation, we inhibited translation

**Fig. 4.**
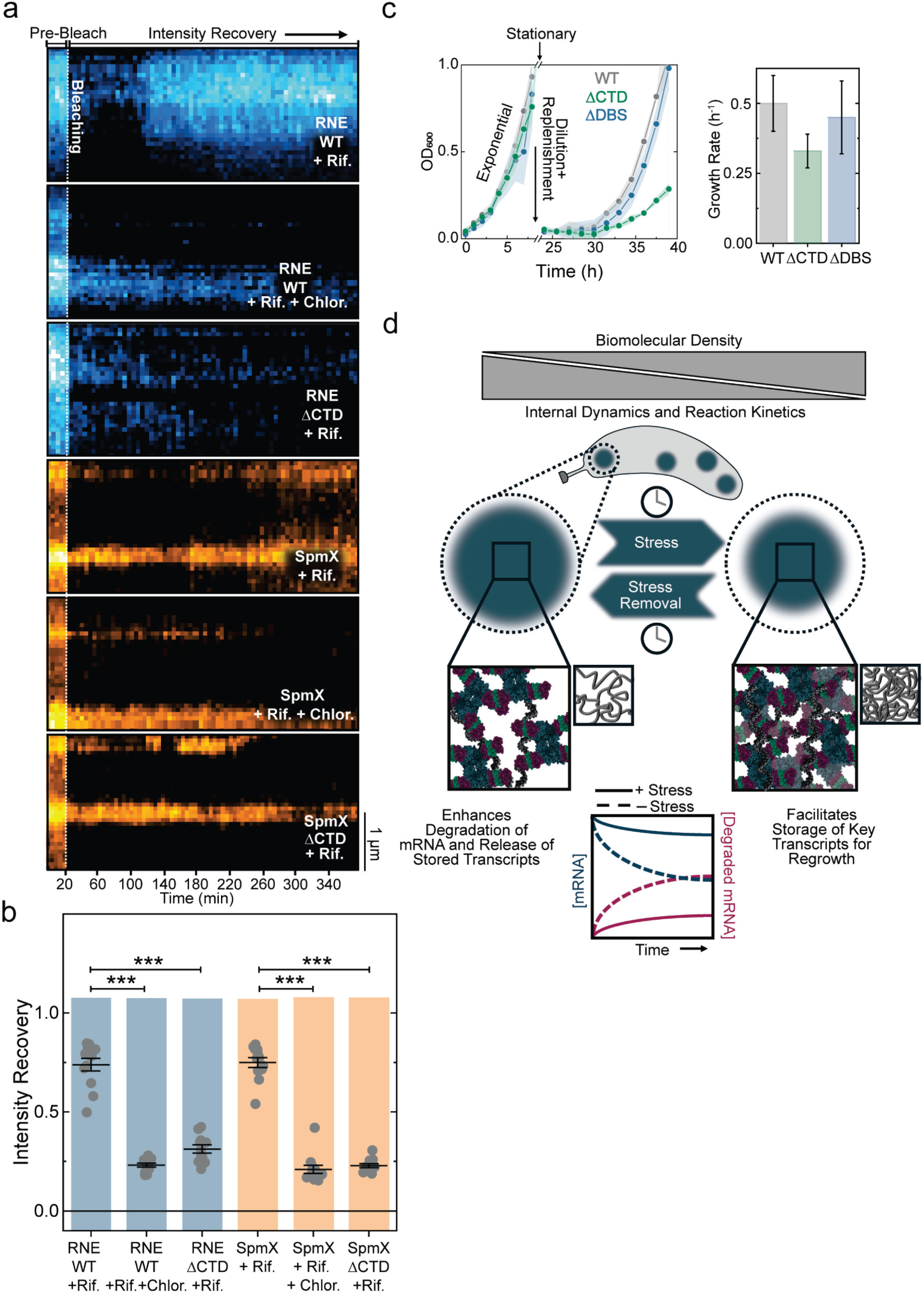
Stored mRNAs can be translated and promote rapid growth. (a) Representative kymographs of SpmX-mEos3.2 in both wild-type RNE and ΔCTD-RNE backgrounds, and of wild-type RNE and RNE-ΔCTD-mEos3.2 alone. Kymographs were generated by measuring cell fluorescence intensity along the long axis of stationary-phase *C. crescentus* cells after nutrient replenishment and inhibition of new transcript synthesis and translation with rifampicin and chloramphenicol, respectively. (b) Quantification of the recovery of the integrated intensity over the whole cell area at *t* = 6 h relative to *t* = 0 h following photobleaching of the whole cell intensity. Each dot below the distribution corresponds to the recovered intensity in one cell (*n* = 12 cells), and the error bars indicate the standard error. *** denotes *p* < 0.001. (c) Functional implications of mRNA storage in BR-bodies. Left: comparison of the growth rate of *C. crescentus* cells with wild-type RNE vs. cells with RNE-ΔCTD (incapable of forming BR-bodies) or RNE-ΔDBS (forms BR-bodies but cannot assemble the degradosome) when cultures are back-diluted into PYE medium after 24 h in stationary phase. Right: growth rates following nutrient replenishment; error bars indicate the standard error. Cells expressing RNE-ΔCTD grow more slowly following nutrient replenishment, whereas cells expressing RNE-ΔDBS grow at rates comparable to wild-type. The initial growth curves, prior to entry into stationary phase, were initiated from back-dilutions of exponentially growing cultures. Three biological replicates were used for each condition. (d) Proposed model for how the material state of BR-bodies tunes their activity, enabling a transition from mRNA decay enhancement during growth to storage during stress. with chloramphenicol, which abolished fluorescence recovery, indicating that the signal arises from newly synthesized protein and not from maturation of pre-existing fluorescent proteins.

We then asked whether BR-bodies are required for this post-stress recovery. In cells expressing RNE-ΔCTD, which are unable to form BR-bodies, fluorescence recovery for SpmX-mEos3.2 and RNE-ΔCTD-mEos3.2 was minimal following nutrient replenishment and was similar to the translation-inhibited wild-type cells (Fig. 4a,b). This result indicates that BR-bodies are required to maintain mRNAs in a translatable state during stationary phase. Importantly, we observed a similar behavior for *spmX* and *rne* transcripts, suggesting that BR-body-mediated transcript storage and translation upon release is a general mechanism not limited to *rne*. To confirm the presence of full-length mRNA that is capable of being translated within stationary phase BR-bodies, we amplified the full-length *ctrA* mRNA by RT-PCR and found that full-length mRNA is indeed present within the BR-body (Extended Data Fig. 5c). The observed translation of released mRNA from BR-bodies suggests that BR-bodies can store functionally intact mRNAs for later translation upon stress removal.

We ultimately assessed the functional implications of stress-induced mRNA storage in BR-bodies by comparing the growth rate of *C. crescentus* cells with wild-type RNE to cells with RNE-ΔCTD after 24 hrs in stationary phase (Fig. 4c). Remarkably, cells with RNE-ΔCTD exhibit a 1.5-fold reduction in growth rate and a lag phase that is 3 hrs longer compared to the wild-type strain following nutrient replenishment. To distinguish the role of BR-body formation from potential pleiotropic effects of disrupting degradosome assembly, we analyzed recovery dynamics in an RNE-ΔDBS strain, which lacks the degradosome binding sites on RNE and therefore cannot assemble the RNA degradosome, but still forms condensates.^11^ Upon nutrient replenishment, RNE-ΔDBS cells recover with kinetics comparable to wild-type, in contrast to the slow recovery observed in the RNE-ΔCTD strain (Fig. 4b,c). These results indicate that degradosome assembly is not required for the efficient exit from stationary phase, and instead support the conclusion that the formation of BR-bodies and their ability to store essential transcripts during stress play a critical role in enabling recovery. Thus, this functional separation-of-function mutant demonstrates that the loss of condensate formation, rather than general defects in RNA metabolism, compromises recovery in the RNE-ΔCTD mutant.

Altogether, these results suggest that during stress, BR-bodies play a critical role in storing essential transcripts, which are then readily available for translation upon return to nutrient-repleted conditions, contributing to a faster transition into exponential growth. This storage mechanism circumvents the need to synthesize new transcripts, thereby facilitating the rapid program of protein production necessary for regrowth. Importantly, we observed in *Agrobacterium tumefaciens* and *Sinorhizobium meliloti* liquid-like BR-bodies also rigidify in stationary phase (Extended Data Fig. 7), suggesting that the material state transition to mRNA storage may be a general feature of BR-bodies.

## Discussion

BR-bodies are multifunctional biomolecular condensates that orchestrate RNA decay and store RNA transcripts during stress (Fig. 4d). Under prolonged stress, BR-bodies rigidify, resulting in decreased internal dynamics, higher biomolecular density, and compaction. This transition extends the residence time of RNE within these condensates, reduces its degradation activity, and leads to an altered degradosome deficient in RhlB. Consequently, RNA transcripts within BR-bodies are stabilized under adverse conditions. Although a significant portion of RNE and mRNAs are distributed outside BR-bodies, our data indicate that BR-body-localized RNE plays the dominant role in transcript stabilization during stationary phase.

The stabilization of mRNA within BR-bodies is a functional bacterial adaptation that allows the preservation of valuable epigenetic information during exposure to a variety of different stressors. Upon stress removal, environmental and cellular conditions shift, triggering the restoration of dynamics and disassembly of BR-bodies. This change facilitates the release of stabilized transcripts back into the cytoplasm, where they can be promptly translated to support recovery and growth. This elegant mechanism, wherein bacteria dynamically modulate the physical and biochemical properties of BR-bodies in response to environmental cues, leverages phase separation and enzyme kinetics to exert temporal and spatial control over RNA degradation and utilization. This mechanism underscores how bacteria use biomolecular condensates not just as active enzymatic reaction sites but also as adaptive tools for managing genetic resources to ensure survival and adaptation in fluctuating environments.

The bacterial aggresome is another subcellular organelle that has recently been reported to store mRNA under arsenic stress in *E. coli*.^12,13,86,87^ Our results indicate that, like aggresomes, BR-bodies store mRNA transcripts during stress conditions. However, the putative mechanisms by which aggresomes and BR-bodies stabilize mRNA differ significantly from one another. Aggresomes have been proposed to protect mRNA by excluding ribonucleases, with protein surface charge playing a crucial role.^86^ Aggresomes are thought to assemble through protein aggregation, and although protein chaperones are enriched within aggresome structures, the molecular trigger of their phase separation has not been uncovered.^12^ However, whether aggresomes in *E. coli* might exclude RNE,^88–90^ the major mRNA decay nuclease, has not yet been explored. In contrast, RNE is the central scaffold necessary for BR-body formation,^11^ and rather than RNE being excluded from mature BR-bodies, the material properties of the BR-bodies shut off the RNase activity. Interestingly, *C. crescentus* BR-bodies have been found to enrich the aggresome marker proteins HslU, DnaK, and ClpB,^43^ suggesting some functional similarity to aggresomes, yet aggresomes have a dark color in phase-contrast microscopy, whereas no such dark aggregates have been observed in *C. crescentus* at stationary phase nor following heat shock, antibiotic treatment, or other stresses.^11,35,91,92^

Despite the mechanistic data in the current study, it is still unclear how mRNA stored within BR-bodies is protected from RNE cleavage when cells transition from stationary to exponential phase. Potential models for future research include the possibility that *C. crescentus* possesses RNE inhibitor proteins stored within BR-bodies, similar to *E. coli* RraA and RraB.^93,94^ Another hypothesis is that the RNE catalytic domain is completely denatured during rigidification and requires refolding by protein chaperones to become active again during the transition back to exponential growth.

Overall, we have determined that the function of the BR-bodies as sites of mRNA storage or decay is dictated by their material state. This model of BR-bodies underscores their role as versatile biomolecular condensates and also demonstrates how biophysical properties influence the biochemical function of condensates. The ability to modulate enzymatic activity through controlled biophysical changes within these condensates reveals a novel paradigm of gene expression regulation, which is pivotal for bacterial survival in fluctuating environments.

## Data Availability

All data generated and analyzed during this study are included in this published article and its Supplementary Information files. Raw source data is available upon request.

## Code Availability

Data processing and analysis scripts for this study were written in MATLAB and Python. The code generated for this study is available on Github. Image analysis scripts: https://github.com/BiteenMatlab/bacterial_condensates; and SMALL-LABS algorithm package: https://github.com/BiteenMatlab/SMALL-LABS.

## Acknowledgments

The authors acknowledge the support of NIH grant R35GM124733 to J.M.S., NIH T32GM142519-03 to HY., and NIH grants R01GM143182 and R01GM144731 to J.S.B. The authors thank Dr. Anthony Vecchiarelli for generously providing access to his wide-field fluorescence microscope. Thanks to Dr. Tamara Hendrickson and the WSU Department of Chemistry for lab space and equipment after the Schrader lab was destroyed in a fire.

The authors declare no competing interests.

## Author contributions

L.A.O.-R. designed, performed, and analyzed experiments (all live-cell single-molecule and ensemble fluorescence imaging, FRAP, FISH, growth curves, and ATP level measurements), interpreted results, and wrote the manuscript. H.Y. designed, performed, and analyzed experiments (strain construction, qRT-PCR, and BONCAT) and interpreted results. V.N., A.H., and YZ designed, performed, and analyzed experiments (all *in vitro* reconstitution and *in vitro* FRAP experiments) and interpreted results. C.A.A. analyzed experiments (developed a script to isolate trajectories inside the condensates) and contributed to writing the revised manuscript. J.C. analyzed experiments (cluster analysis). J.M.S. provided expertise in the design and construction of the strains, interpreted results, and wrote the manuscript. J.S.B. provided expertise in the design of live-cell imaging experiments, designed experiments, interpreted results, and wrote the manuscript.

## References

1. Banani, S. F., Lee, H. O., Hyman, A. A. & Rosen, M. K. Biomolecular condensates: organizers of cellular biochemistry. Nat. Rev. Mol. Cell Biol. 18, 285–298 (2017).

2. Boeynaems, S., Alberti, S., Fawzi, N. L., Mittag, T., Polymenidou, M., Rousseau, F., Schymkowitz, J., Shorter, J., Wolozin, B., Bosch, L. V. D., Tompa, P. & Fuxreiter, M. Protein Phase Separation: A New Phase in Cell Biology. Trends Cell Biol. 28, 420–435 (2018).

3. Yoo, H., Triandafillou, C. & Drummond, D. A. Cellular sensing by phase separation: Using the process, not just the products. J. Biol. Chem. 294, 7151–7159 (2019).

4. Ditlev, J. A., Case, L. B. & Rosen, M. K. Who’s In and Who’s Out—Compositional Control of Biomolecular Condensates. J. Mol. Biol. 430, 4666–4684 (2018).

5. Nakamura, H., DeRose, R. & Inoue, T. Harnessing biomolecular condensates in living cells. J. Biochem. (Tokyo) 166, 13–27 (2019).

6. Aguzzi, A. & Altmeyer, M. Phase Separation: Linking Cellular Compartmentalization to Disease. Trends Cell Biol. 26, 547–558 (2016).

7. Ramaswami, M., Taylor, J. P. & Parker, R. Altered Ribostasis: RNA-Protein Granules in Degenerative Disorders. Cell 154, 727–736 (2013).

8. Zhu, L., Richardson, T. M., Wacheul, L., Wei, M.-T., Feric, M., Whitney, G., Lafontaine, D. L. J. & Brangwynne, C. P. Controlling the material properties and rRNA processing function of the nucleolus using light. Proc. Natl. Acad. Sci. 116, 17330–17335 (2019).

9. Dorone, Y., Boeynaems, S., Flores, E., Jin, B., Hateley, S., Bossi, F., Lazarus, E., Pennington, J. G., Michiels, E., Decker, M. D., Vints, K., Baatsen, P., Bassel, G. W., Otegui, M. S., Holehouse, A. S., Exposito-Alonso, M., Sukenik, S., Gitler, A. D. & Rhee, S. Y. A prion-like protein regulator of seed germination undergoes hydration-dependent phase separation. Cell 184, 4284–4298.e27 (2021).

10. Bose, M., Lampe, M., Mahamid, J. & Ephrussi, A. Liquid-to-solid phase transition of *oskar* ribonucleoprotein granules is essential for their function in *Drosophila* embryonic development. Cell 185, 1308–1324.e23 (2022).

11. Al-Husini, N., Tomares, D. T., Bitar, O., Childers, W. S. & Schrader, J. M. α-Proteobacterial RNA Degradosomes Assemble Liquid-Liquid Phase-Separated RNP Bodies. Mol. Cell 71, 1027–1039.e14 (2018).

12. Pu, Y., Li, Y., Jin, X., Tian, T., Ma, Q., Zhao, Z., Lin, S., Chen, Z., Li, B., Yao, G., Leake, M. C., Lo, C.-J. & Bai, F. ATP-Dependent Dynamic Protein Aggregation Regulates Bacterial Dormancy Depth Critical for Antibiotic Tolerance. Mol. Cell 73, 143–156.e4 (2019).

13. Jin, X., Lee, J.-E., Schaefer, C., Luo, X., Wollman, A. J. M., Payne-Dwyer, A. L., Tian, T., Zhang, X., Chen, X., Li, Y., McLeish, T. C. B., Leake, M. C. & Bai, F. Membraneless organelles formed by liquid-liquid phase separation increase bacterial fitness. Sci. Adv. 7, eabh2929 (2021).

14. Heinkel, F., Abraham, L., Ko, M., Chao, J., Bach, H., Hui, L. T., Li, H., Zhu, M., Ling, Y. M., Rogalski, J. C., Scurll, J., Bui, J. M., Mayor, T., Gold, M. R., Chou, K. C., Av-Gay, Y., McIntosh, L. P. & Gsponer, J. Phase separation and clustering of an ABC transporter in Mycobacterium tuberculosis. Proc. Natl. Acad. Sci. 116, 16326–16331 (2019).

15. MacCready, J. S., Basalla, J. L. & Vecchiarelli, A. G. Origin and Evolution of Carboxysome Positioning Systems in Cyanobacteria. Mol. Biol. Evol. 37, 1434–1451 (2020).

16. McQuail, J., Carpousis, A. J. & Wigneshweraraj, S. The association between Hfq and RNase E in long-term nitrogen-starved Escherichia coli. Mol. Microbiol. 117, 54–66 (2022).

17. Azaldegui, C. A., Vecchiarelli, A. G. & Biteen, J. S. The emergence of phase separation as an organizing principle in bacteria. Biophys. J. 120, 1123–1138 (2021).

18. Saurabh, S., Chong, T. N., Bayas, C., Dahlberg, P. D., Cartwright, H. N., Moerner, W. E. & Shapiro, L. ATP-responsive biomolecular condensates tune bacterial kinase signaling. Sci. Adv. 8, eabm6570 (2022).

19. Lasker, K., von Diezmann, L., Zhou, X., Ahrens, D. G., Mann, T. H., Moerner, W. E. & Shapiro, L. Selective sequestration of signalling proteins in a membraneless organelle reinforces the spatial regulation of asymmetry in Caulobacter crescentus. Nat. Microbiol. 5, 418–429 (2020).

20. Surovtsev, I. V. & Jacobs-Wagner, C. Subcellular Organization: A Critical Feature of Bacterial Cell Replication. Cell 172, 1271–1293 (2018).

21. Treuner-Lange, A. & Søgaard-Andersen, L. Regulation of cell polarity in bacteria. J. Cell Biol. 206, 7–17 (2014).

22. Hammarlöf, D. L., Bergman, J. M., Garmendia, E. & Hughes, D. Turnover of mRNAs is one of the essential functions of RNase E. Mol. Microbiol. 98, 34–45 (2015).

23. Ono, M. & Kuwano, M. A conditional lethal mutation in an *Escherichia coli* strain with a longer chemical lifetime of messenger RNA. J. Mol. Biol. 129, 343–357 (1979).

24. Kedersha, N., Stoecklin, G., Ayodele, M., Yacono, P., Lykke-Andersen, J., Fritzler, M. J., Scheuner, D., Kaufman, R. J., Golan, D. E. & Anderson, P. Stress granules and processing bodies are dynamically linked sites of mRNP remodeling. J. Cell Biol. 169, 871–884 (2005).

25. Hubstenberger, A., Courel, M., Bénard, M., Souquere, S., Ernoult-Lange, M., Chouaib, R., Yi, Z., Morlot, J.-B., Munier, A., Fradet, M., Daunesse, M., Bertrand, E., Pierron, G., Mozziconacci, J., Kress, M. & Weil, D. P-Body Purification Reveals the Condensation of Repressed mRNA Regulons. Mol. Cell 68, 144–157.e5 (2017).

26. Youn, J.-Y., Dyakov, B. J. A., Zhang, J., Knight, J. D. R., Vernon, R. M., Forman-Kay, J. D. & Gingras, A.-C. Properties of Stress Granule and P-Body Proteomes. Mol. Cell 76, 286–294 (2019).

27. Rathnayaka-Mudiyanselage, I., Nandana, V. & Schrader, J. Proteomic composition of eukaryotic and bacterial RNA decay condensates suggests convergent evolution. Curr. Opin. Microbiol. 79, 102467 (2024).

28. Brengues, M., Teixeira, D. & Parker, R. Movement of Eukaryotic mRNAs Between Polysomes and Cytoplasmic Processing Bodies. Science 310, 486–489 (2005).

29. Sheth, U. & Parker, R. Decapping and Decay of Messenger RNA Occur in Cytoplasmic Processing Bodies. Science 300, 805–808 (2003).

30. Xing, W., Muhlrad, D., Parker, R. & Rosen, M. K. A quantitative inventory of yeast P body proteins reveals principles of composition and specificity. eLife 9, e56525 (2020).

31. Blake, L. A., Watkins, L., Liu, Y., Inoue, T. & Wu, B. A rapid inducible RNA decay system reveals fast mRNA decay in P-bodies. Nat. Commun. 15, 2720 (2024).

32. Wheeler, J. R., Matheny, T., Jain, S., Abrisch, R. & Parker, R. Distinct stages in stress granule assembly and disassembly. eLife 5, e18413 (2016).

33. Moon, S. L., Morisaki, T., Khong, A., Lyon, K., Parker, R. & Stasevich, T. J. Multicolour single-molecule tracking of mRNA interactions with RNP granules. Nat. Cell Biol. 21, 162–168 (2019).

34. Mateju, D., Franzmann, T. M., Patel, A., Kopach, A., Boczek, E. E., Maharana, S., Lee, H. O., Carra, S., Hyman, A. A. & Alberti, S. An aberrant phase transition of stress granules triggered by misfolded protein and prevented by chaperone function. EMBO J. 36, 1669–1687 (2017).

35. Al-Husini, N., Tomares, D. T., Pfaffenberger, Z. J., Muthunayake, N. S., Samad, M. A., Zuo, T., Bitar, O., Aretakis, J. R., Bharmal, M.-H. M., Gega, A., Biteen, J. S., Childers, W. S. & Schrader, J. M. BR-Bodies Provide Selectively Permeable Condensates that Stimulate mRNA Decay and Prevent Release of Decay Intermediates. Mol. Cell 78, 670–682.e8 (2020).

36. Chen, H., Shiroguchi, K., Ge, H. & Xie, X. S. Genome-wide study of mRNA degradation and transcript elongation in Escherichia coli. Mol. Syst. Biol. 11, 781 (2015).

37. Barnett, T. C., Bugrysheva, J. V. & Scott, J. R. Role of mRNA Stability in Growth Phase Regulation of Gene Expression in the Group A Streptococcus. J. Bacteriol. 189, 1866–1873 (2007).

38. Hambraeus, G., von Wachenfeldt, C. & Hederstedt, L. Genome-wide survey of mRNA half-lives in Bacillus subtilis identifies extremely stable mRNAs. Mol. Genet. Genomics 269, 706–714 (2003).

39. Dressaire, C., Pobre, V., Laguerre, S., Girbal, L., Arraiano, C. M. & Cocaign-Bousquet, M. PNPase is involved in the coordination of mRNA degradation and expression in stationary phase cells of Escherichia coli. BMC Genomics 19, 848 (2018).

40. Esquerré, T., Laguerre, S., Turlan, C., Carpousis, A. J., Girbal, L. & Cocaign-Bousquet, M. Dual role of transcription and transcript stability in the regulation of gene expression in Escherichia coli cells cultured on glucose at different growth rates. Nucleic Acids Res. 42, 2460–2472 (2014).

41. Morrison, J. M., Anderson, K. L., Beenken, K. E., Smeltzer, M. S. & Dunman, P. M. The staphylococcal accessory regulator, SarA, is an RNA-binding protein that modulates the mRNA turnover properties of late-exponential and stationary phase Staphylococcus aureus cells. Front. Cell. Infect. Microbiol. 2, (2012).

42. Zhang, M., Chang, H., Zhang, Y., Yu, J., Wu, L., Ji, W., Chen, J., Liu, B., Lu, J., Liu, Y., Zhang, J., Xu, P. & Xu, T. Rational design of true monomeric and bright photoactivatable fluorescent proteins. Nat. Methods 9, 727–729 (2012).

43. Nandana, V., Rathnayaka-Mudiyanselage, I. W., Muthunayake, N. S., Hatami, A., Mousseau, C. B., Ortiz-Rodríguez, L. A., Vaishnav, J., Collins, M., Gega, A., Mallikaarachchi, K. S., Yassine, H., Ghosh, A., Biteen, J. S., Zhu, Y., Champion, M. M., Childers, W. S. & Schrader, J. M. The BR-body proteome contains a complex network of protein-protein and protein-RNA interactions. Cell Rep. 42, 113229 (2023).

44. Hardwick, S. W., Chan, V. S. Y., Broadhurst, R. W. & Luisi, B. F. An RNA degradosome assembly in Caulobacter crescentus. Nucleic Acids Res. 39, 1449–1459 (2011).

45. Ester, M., Kriegel, H.-P., Sander, J. & Xu, X. A density-based algorithm for discovering clusters in large spatial databases with noise. in Proc. Second Int. Conf. Knowl. Discov. Data Min. 226–231 (AAAI Press, 1996).

46. Parry, B. R., Surovtsev, I. V., Cabeen, M. T., O’Hern, C. S., Dufresne, E. R. & Jacobs-Wagner, C. The Bacterial Cytoplasm Has Glass-like Properties and Is Fluidized by Metabolic Activity. Cell 156, 183–194 (2014).

47. Kroschwald, S., Maharana, S., Mateju, D., Malinovska, L., Nüske, E., Poser, I., Richter, D. & Alberti, S. Promiscuous interactions and protein disaggregases determine the material state of stress-inducible RNP granules. eLife 4, e06807 (2015).

48. Kroschwald, S., Maharana, S. & Simon, A. Hexanediol: a chemical probe to investigate the material properties of membrane-less compartments. Matters (2017). doi:10.19185/matters.201702000010

49. Lin, Y., Mori, E., Kato, M., Xiang, S., Wu, L., Kwon, I. & McKnight, S. L. Toxic PR Poly-Dipeptides Encoded by the C9orf72 Repeat Expansion Target LC Domain Polymers. Cell 167, 789–802.e12 (2016).

50. Hondele, M., Sachdev, R., Heinrich, S., Wang, J., Vallotton, P., Fontoura, B. M. A. & Weis, K. DEAD-box ATPases are global regulators of phase-separated organelles. Nature 573, 144–148 (2019).

51. Deng, Y., Beahm, D. R., Ionov, S. & Sarpeshkar, R. Measuring and modeling energy and power consumption in living microbial cells with a synthetic ATP reporter. BMC Biol. 19, 101 (2021).

52. Weber, S. C., Spakowitz, A. J. & Theriot, J. A. Nonthermal ATP-dependent fluctuations contribute to the in vivo motion of chromosomal loci. Proc. Natl. Acad. Sci. 109, 7338–7343 (2012).

53. Russell, R., Jarmoskaite, I. & Lambowitz, A. M. Toward a molecular understanding of RNA remodeling by DEAD-box proteins. RNA Biol. 10, 44–55 (2013).

54. Li, X.-Z., Plésiat, P. & Nikaido, H. The Challenge of Efflux-Mediated Antibiotic Resistance in Gram-Negative Bacteria. Clin. Microbiol. Rev. 28, 337–418 (2015).

55. Nathans, D. Puromycin inhibition of protein synthesis: incorporation of puromycin into peptide chains. Proc. Natl. Acad. Sci. 51, 585–592 (1964).

56. Oh, E., Becker, A. H., Sandikci, A., Huber, D., Chaba, R., Gloge, F., Nichols, R. J., Typas, A., Gross, C. A., Kramer, G., Weissman, J. S. & Bukau, B. Selective Ribosome Profiling Reveals the Cotranslational Chaperone Action of Trigger Factor In Vivo. Cell 147, 1295–1308 (2011).

57. Pontes, M. H., Sevostyanova, A. & Groisman, E. A. When Too Much ATP Is Bad for Protein Synthesis. J. Mol. Biol. 427, 2586–2594 (2015).

58. Stouthamer, A. H. A theoretical study on the amount of ATP required for synthesis of microbial cell material. Antonie Van Leeuwenhoek 39, 545–565 (1973).

59. Patel, A., Lee, H. O., Jawerth, L., Maharana, S., Jahnel, M., Hein, M. Y., Stoynov, S., Mahamid, J., Saha, S., Franzmann, T. M., Pozniakovski, A., Poser, I., Maghelli, N., Royer, L. A., Weigert, M., Myers, E. W., Grill, S., Drechsel, D., Hyman, A. A. & Alberti, S. A Liquid-to-Solid Phase Transition of the ALS Protein FUS Accelerated by Disease Mutation. Cell 162, 1066–1077 (2015).

60. Molliex, A., Temirov, J., Lee, J., Coughlin, M., Kanagaraj, A. P., Kim, H. J., Mittag, T. & Taylor, J. P. Phase Separation by Low Complexity Domains Promotes Stress Granule Assembly and Drives Pathological Fibrillization. Cell 163, 123–133 (2015).

61. Zhang, H., Elbaum-Garfinkle, S., Langdon, E. M., Taylor, N., Occhipinti, P., Bridges, A. A., Brangwynne, C. P. & Gladfelter, A. S. RNA Controls PolyQ Protein Phase Transitions. Mol. Cell 60, 220–230 (2015).

62. Lin, Y., Protter, D. S. W., Rosen, M. K. & Parker, R. Formation and Maturation of Phase-Separated Liquid Droplets by RNA-Binding Proteins. Mol. Cell 60, 208–219 (2015).

63. Xiang, S., Kato, M., Wu, L. C., Lin, Y., Ding, M., Zhang, Y., Yu, Y. & McKnight, S. L. The LC Domain of hnRNPA2 Adopts Similar Conformations in Hydrogel Polymers, Liquid-like Droplets, and Nuclei. Cell 163, 829–839 (2015).

64. Watanabe, H. Viscoelasticity and dynamics of entangled polymers. Prog. Polym. Sci. 24, 1253–1403 (1999).

65. Halfmann, R. A glass menagerie of low complexity sequences. Curr. Opin. Struct. Biol. 38, 18– 25 (2016).

66. Mittag, T. & Pappu, R. V. A conceptual framework for understanding phase separation and addressing open questions and challenges. Mol. Cell 82, 2201–2214 (2022).

67. Boke, E., Ruer, M., Wühr, M., Coughlin, M., Lemaitre, R., Gygi, S. P., Alberti, S., Drechsel, D., Hyman, A. A. & Mitchison, T. J. Amyloid-like Self-Assembly of a Cellular Compartment. Cell 166, 637–650 (2016).

68. Linsenmeier, M., Hondele, M., Grigolato, F., Secchi, E., Weis, K. & Arosio, P. Dynamic arrest and aging of biomolecular condensates are modulated by low-complexity domains, RNA and biochemical activity. Nat. Commun. 13, 3030 (2022).

69. Bertrand, E., Chartrand, P., Schaefer, M., Shenoy, S. M., Singer, R. H. & Long, R. M. Localization of ASH1 mRNA Particles in Living Yeast. Mol. Cell 2, 437–445 (1998).

70. Tutucci, E., Vera, M., Biswas, J., Garcia, J., Parker, R. & Singer, R. H. An improved MS2 system for accurate reporting of the mRNA life cycle. Nat. Methods 15, 81–89 (2018).

71. Nandana, V., Al-Husini, N., Vaishnav, A., Dilrangi, K. H. & Schrader, J. M. Caulobacter crescentus RNase E condensation contributes to autoregulation and fitness. Mol. Biol. Cell 35, ar104 (2024).

72. Awram, P. & Smit, J. The Caulobacter crescentus Paracrystalline S-Layer Protein Is Secreted by an ABC Transporter (Type I) Secretion Apparatus. J. Bacteriol. 180, 3062–3069 (1998).

73. Wheeler, R. T. & Shapiro, L. Differential Localization of Two Histidine Kinases Controlling Bacterial Cell Differentiation. Mol. Cell 4, 683–694 (1999).

74. Radhakrishnan, S. K., Thanbichler, M. & Viollier, P. H. The dynamic interplay between a cell fate determinant and a lysozyme homolog drives the asymmetric division cycle of Caulobacter crescentus. Genes Dev. 22, 212–225

75. Ryan, K. R., Judd, E. M. & Shapiro, L. The CtrA Response Regulator Essential for *Caulobacter crescentus* Cell-cycle Progression Requires a Bipartite Degradation Signal for Temporally Controlled Proteolysis. J. Mol. Biol. 324, 443–455 (2002).

76. Wortinger, M., Sackett, M. J. & Brun, Y. V. CtrA mediates a DNA replication checkpoint that prevents cell division in Caulobacter crescentus. EMBO J. 19, 4503–4512 (2000).

77. Laub, M. T., Chen, S. L., Shapiro, L. & McAdams, H. H. Genes directly controlled by CtrA, a master regulator of the Caulobacter cell cycle. Proc. Natl. Acad. Sci. 99, 4632–4637 (2002).

78. Laub, M. T., McAdams, H. H., Feldblyum, T., Fraser, C. M. & Shapiro, L. Global Analysis of the Genetic Network Controlling a Bacterial Cell Cycle. Science 290, 2144–2148 (2000).

79. Quon, K. C., Marczynski, G. T. & Shapiro, L. Cell Cycle Control by an Essential Bacterial Two-Component Signal Transduction Protein. Cell 84, 83–93 (1996).

80. LeCuyer, K. A., Behlen, L. S. & Uhlenbeck, O. C. Mutants of the Bacteriophage MS2 Coat Protein That Alter Its Cooperative Binding to RNA. Biochemistry 34, 10600–10606 (1995).

81. Garcia, J. F. & Parker, R. MS2 coat proteins bound to yeast mRNAs block 5′ to 3′ degradation and trap mRNA decay products: implications for the localization of mRNAs by MS2-MCP system. RNA 21, 1393–1395 (2015).

82. Heinrich, S., Sidler, C. L., Azzalin, C. M. & Weis, K. Stem–loop RNA labeling can affect nuclear and cytoplasmic mRNA processing. RNA 23, 134–141 (2017).

83. Garcia, J. F. & Parker, R. Ubiquitous accumulation of 3′ mRNA decay fragments in Saccharomyces cerevisiae mRNAs with chromosomally integrated MS2 arrays. RNA 22, 657 (2016).

84. Guan, J., Hurto, R. L., Rai, A., Azaldegui, C. A., Ortiz-Rodríguez, L. A., Biteen, J. S., Freddolino, L. & Jakob, U. HP-Bodies – Ancestral Condensates that Regulate RNA Turnover and Protein Translation in Bacteria. 2025.02.06.636932 Preprint at 10.1101/2025.02.06.636932 (2025)

85. Goldberger, O., Szoke, T., Nussbaum-Shochat, A. & Amster-Choder, O. Heterotypic phase separation of Hfq is linked to its roles as an RNA chaperone. Cell Rep. 41, 111881 (2022).

86. Pei, L., Xian, Y., Yan, X., Schaefer, C., Syeda, A. H., Howard, J., Liao, H., Bai, F., Leake, M. C. & Pu, Y. Bacterial stress granule protects mRNA through ribonucleases exclusion. 2024.04.27.591437 Preprint at 10.1101/2024.04.27.591437 (2024)

87. Bollen, C., Louwagie, S., Deroover, F., Duverger, W., Khodaparast, L., Khodaparast, L., Hofkens, D., Schymkowitz, J., Rousseau, F., Dewachter, L. & Michiels, J. Composition and liquid-to-solid maturation of protein aggregates contribute to bacterial dormancy development and recovery. Nat. Commun. 16, 1046 (2025).

88. Mudd, E. A., Krisch, H. M. & Higgins, C. F. RNase E, an endoribonuclease, has a general role in the chemical decay of Escherichia coli mRNA: evidence that rne and ams are the same genetic locus. Mol. Microbiol. 4, 2127–2135 (1990).

89. Górna, M. W., Carpousis, A. J. & Luisi, B. F. From conformational chaos to robust regulation: the structure and function of the multi-enzyme RNA degradosome. Q. Rev. Biophys. 45, 105–145 (2012).

90. Bandyra, K. J., Bouvier, M., Carpousis, A. J. & Luisi, B. F. The social fabric of the RNA degradosome. Biochim. Biophys. Acta BBA - Gene Regul. Mech. 1829, 514–522 (2013).

91. Schramm, F. D., Schroeder, K., Alvelid, J., Testa, I. & Jonas, K. Growth-driven displacement of protein aggregates along the cell length ensures partitioning to both daughter cells in Caulobacter crescentus. Mol. Microbiol. 111, 1430–1448 (2019).

92. Wortinger, M. A., Quardokus, E. M. & Brun, Y. V. Morphological adaptation and inhibition of cell division during stationary phase in Caulobacter crescentus. Mol. Microbiol. 29, 963–973 (1998).

93. Lee, K., Zhan, X., Gao, J., Qiu, J., Feng, Y., Meganathan, R., Cohen, S. N. & Georgiou, G. RraA: a Protein Inhibitor of RNase E Activity that Globally Modulates RNA Abundance in E. coli. Cell 114, 623–634 (2003).

94. Górna, M. W., Pietras, Z., Tsai, Y.-C., Callaghan, A. J., Hernández, H., Robinson, C. V. & Luisi, B. F. The regulatory protein RraA modulates RNA-binding and helicase activities of the E. coli RNA degradosome. RNA 16, 553–562 (2010).

