## Supplemental Information for "Stress Changes the Material State of a Bacterial Biomolecular Condensate and Shifts its Function from mRNA Decay to Storage"


**Contents:**


- Extended Data Figures 1 – 7
- Methods
- Supplementary Figures 1– 10
- Supplementary Tables 1– 5
- Supplementary Movie Captions
- Supplementary References

Extended Data Figures

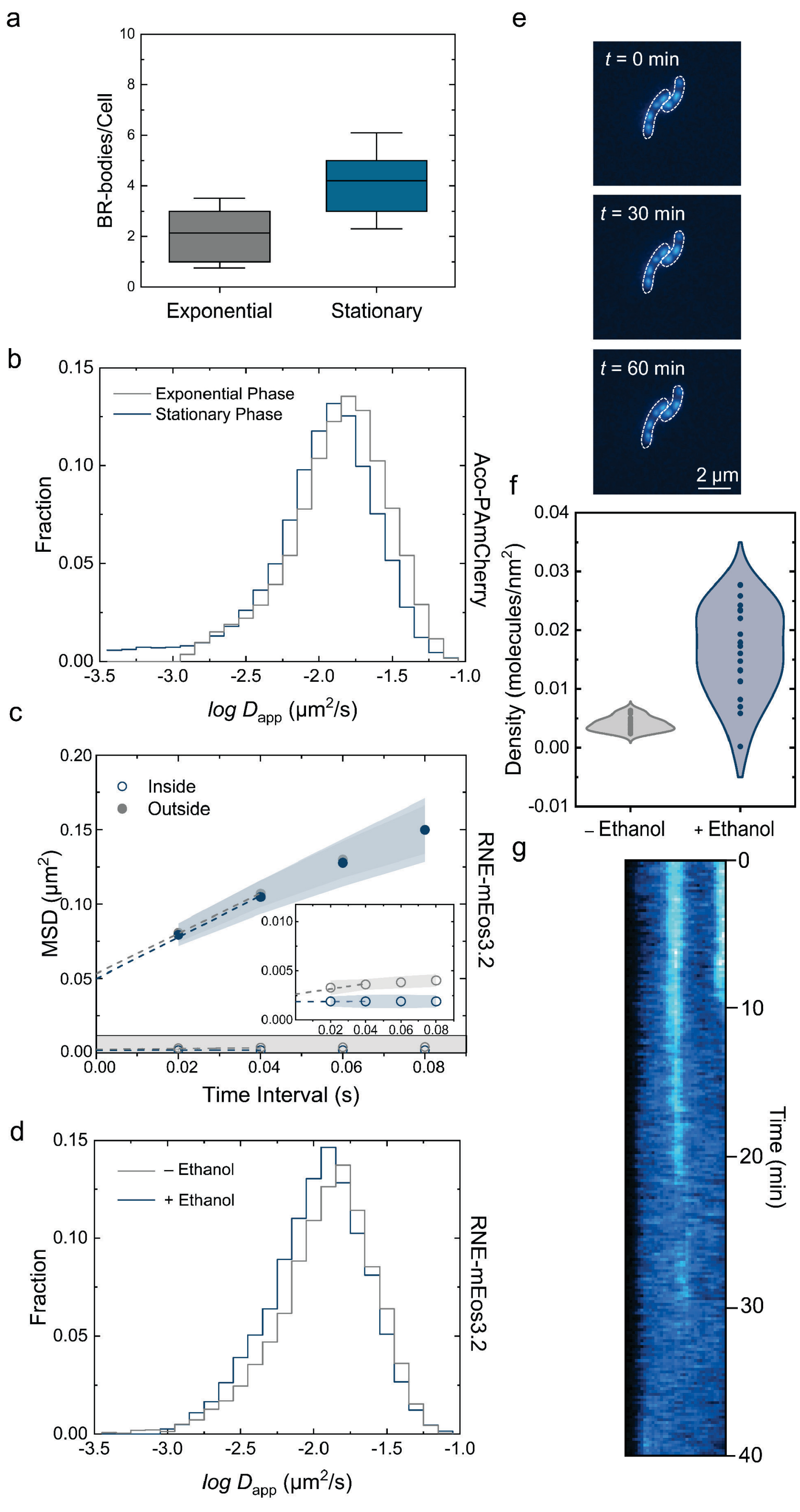


**Extended Data Fig. 1│ Phase-dependent and chemically induced changes in BR-body dynamics, composition, and material state.**

**Extended Data Fig. 1 │ Phase-dependent and chemically induced changes in BR-body dynamics, composition, and material state.** (a) Number of BR-bodies per cell in exponential and stationary phase. Error bars are the standard deviation. *n* = 131 and 109 cells for exponential and stationary phase, respectively. (b) Distributions of diffusion coefficients of aconitase-PAmCherry trajectories inside BR-bodies during exponential and stationary phase. (c) Mean squared displacement (MSD) as a function of time lag for RNE-mEos3.2 trajectories inside and outside BR-bodies in exponential and stationary phases. These curves were used to extract ensemble-averaged diffusion coefficients. Inset: Enlarged view of the highlighted gray region, showing the MSD as a function of time interval for RNE-mEos3.2 trajectories within BR-bodies. Shaded regions indicate the SEM. (d) Diffusion coefficient distributions of RNE-mEos3.2 trajectories inside BR-bodies during exponential phase, with or without treatment with 10% ethanol. (e) Representative images of *C. crescentus* cells (white outlines) expressing RNE-mEos3.2 (blue) over time, *t*, during exponential phase with treatment with 10% ethanol. Scale bar: 2 μm. (f) Molecular density of RNE-mEos3.2 inside BR-bodies in exponential phase with and without 10% ethanol treatment. *N* = 20 clusters. (g) Representative kymograph of a stationary phase cell (x-axis) expressing RNE-mEos3.2 following treatment with 5% 1,6-hexanediol, showing dissolution of BR-bodies over the course of 40 min.

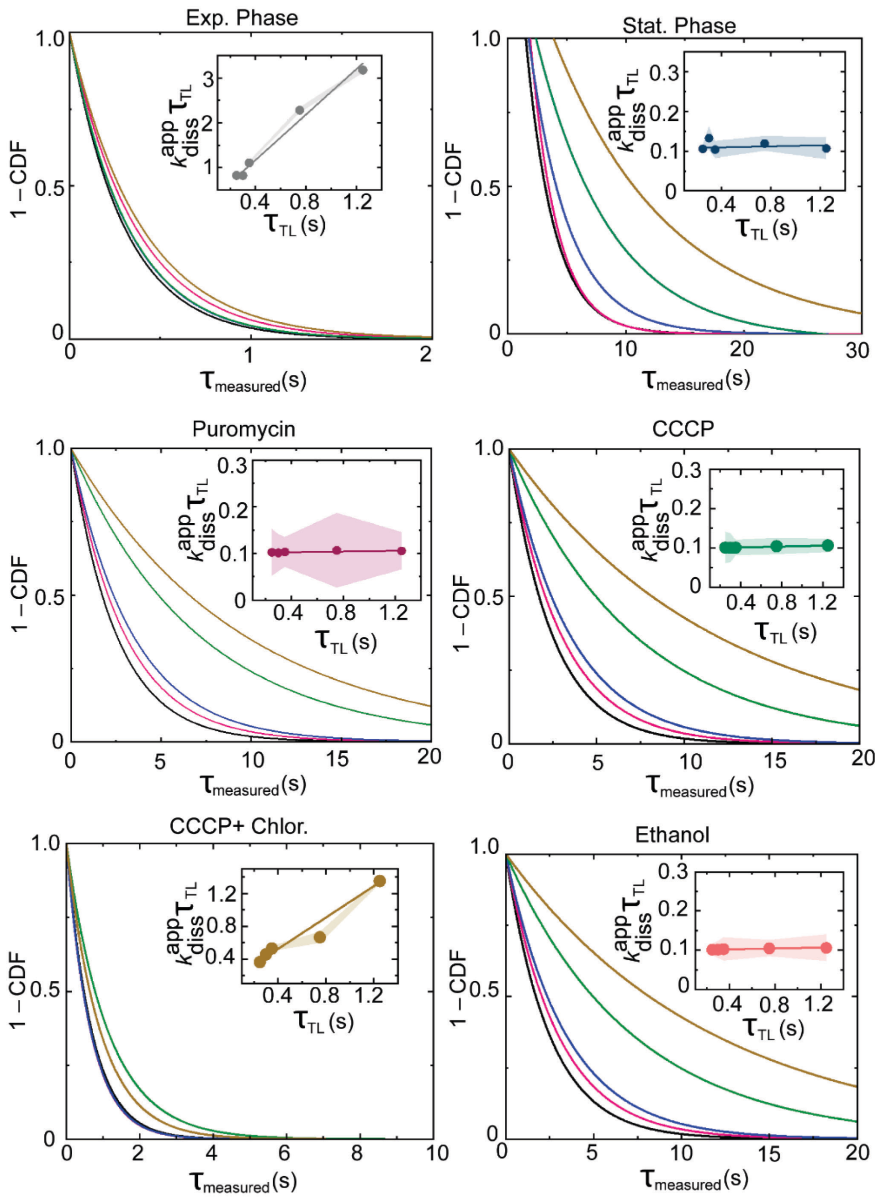


**Extended Data Fig. 2 | Change in RNE-mEos3.2 residence time inside BR-bodies in *C. crescentus* cells across growth phases and under different acute stress conditions.** Fitted exponential decay curves from plots of 1 minus the cumulative probability function (CDF) of observed residence times imaged by slow tracking ($t_{integration}$ = 250 ms) using continuous imaging (black) and time delays (pink, blue, green, and gold for 300, 350, 750, and 1250 ms, respectively). The inset within each (1 – CDF) vs. $\tau_{measured}$ plot shows the linear fit of $k_{diss}^{app}\tau_{TL}$ *vs*. $\tau_{TL}$, from which the dissociation rate constant, $k_{diss}^{app}\tau_{TL}$, which is the reciprocal of the residence time, is extracted (Methods Equation 1). For each $\tau_{measured}$, at least three biological replicates were analyzed.

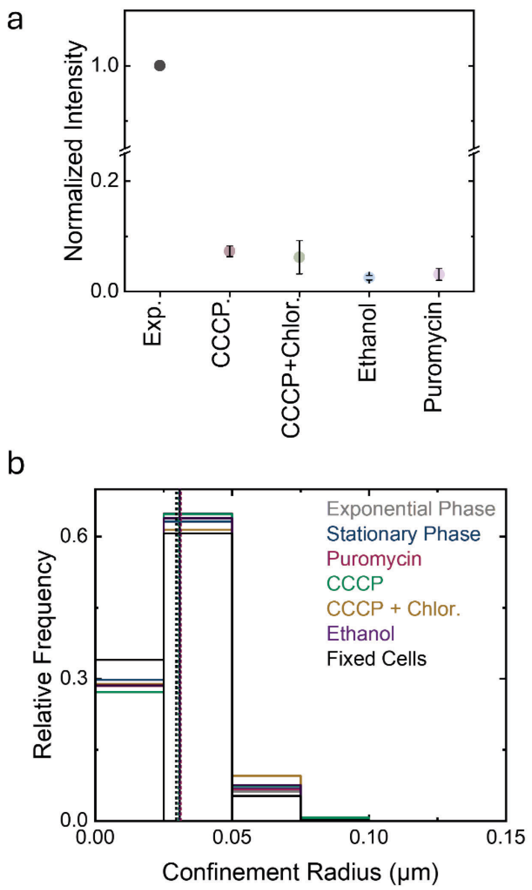


**Extended Data Fig. 3│Comparison of translation levels and diffusivity of BR-bodies in *C. crescentus* cells across growth phases and following acute treatments.** (a) Relative translational levels in exponential phase in the presence and absence of CCCP, both CCCP and chloramphenicol, ethanol, and puromycin using Biorthogonal Non-Canonical Amino Acid Tagging (BONCAT). Error bars: standard deviation from three replicates. (b) The distributions of confinement radii for RNE-mEos3.2 focus trajectories in living cells (colors) were compared to the confinement radii of trajectories in fixed cells (black) to differentiate between diffusing and static foci. Only the first eight frames of each focus trajectory were used for this analysis. Vertical dashed lines: medians of the distributions.

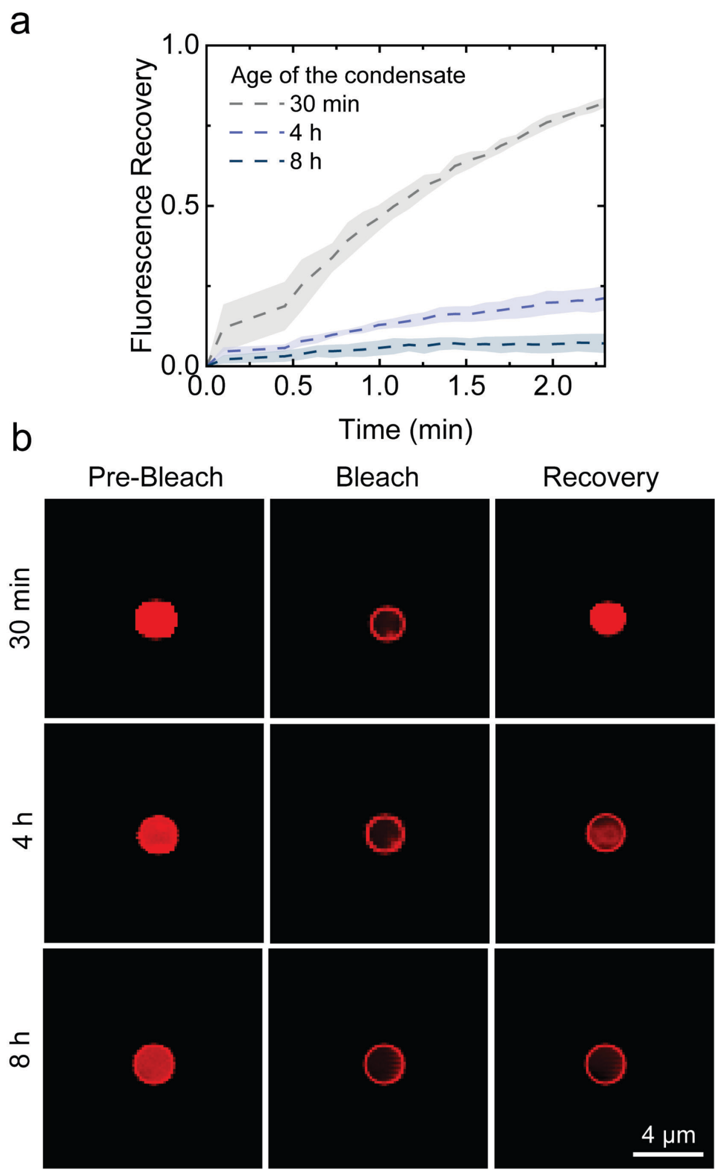


**Extended Data Fig. 4 | Aging of *in vitro*-reconstituted RNE condensates leads to decreased internal dynamics as measured by FRAP.** (a) Fluorescence recovery after photobleaching (FRAP) and (b) representative images of reconstituted RNE-Cy5 *in vitro* condensates at the indicated time points after formation. Error bars represent the standard deviation. Three replicates were obtained for each condition.

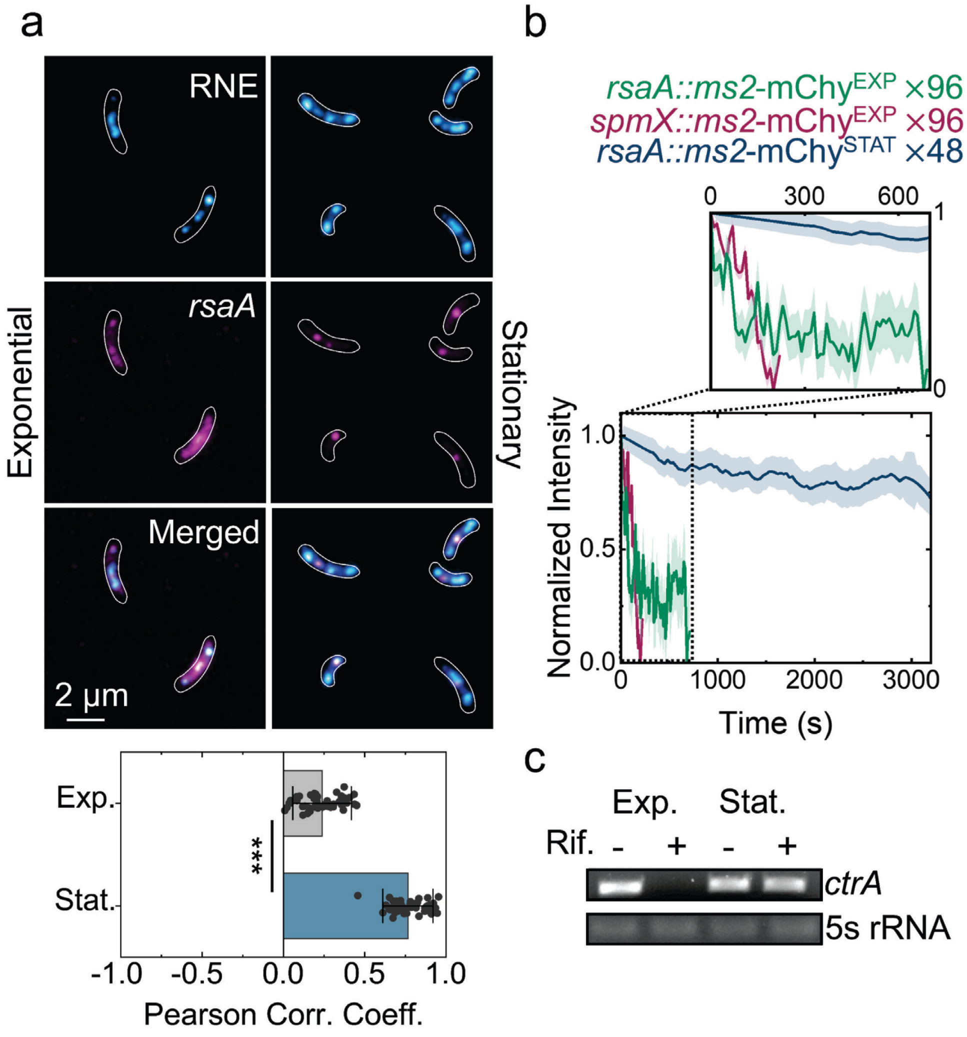


**Extended Data Fig. 5 | Comparison of mRNA transcript stability between exponential and stationary phases.** (a) Representative two-color images showing higher recruitment of the *rsaA* (48× array) transcript (magenta) to BR-bodies (blue) in *C. crescentus* cells (white outlines) during stationary phase (left) relative to exponential phase (right). Bottom: Differences in recruitment quantified by the Pearson correlation coefficients between channels. Error bars: standard deviation from 50 cells analyzed per phase. *** denotes $p < 0.001$. Circles represent the PCC of each cell. (b) Fluorescence decay profiles of *rsaA* and *spmX* foci. (c) Full-length *ctrA* mRNA remains stable in stationary phase. Total RNA was collected from cells in exponential phase and stationary phase with or without a 16-minute rifampicin treatment. The mRNAs were reverse transcribed into full-length *ctrA* or 5S rRNA cDNA, which was then used as a template to amplify the PCR product. 5S rRNA was used as a loading control. The 5S rRNA was PCR amplified for 10 cycles, while *ctrA* was amplified for 25 – 30 cycles.

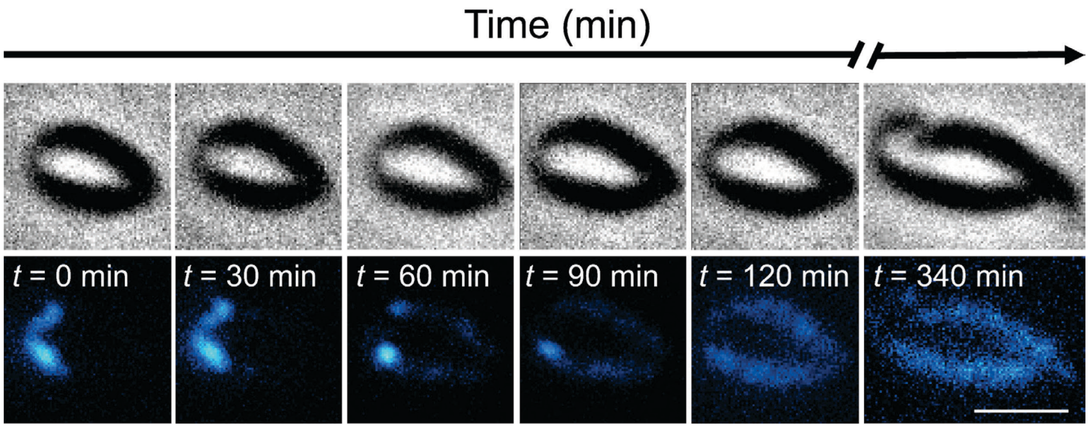


**Extended Data Fig. 6│BR-bodies disassemble upon nutrient replenishment**. Representative images of *C. crescentus* cells (top: phase contrast) and RNE-YFP (bottom: blue) as a function of imaging time, *t*. Fresh M2G medium was added at *t* = 0 min. Scale bar: 2 μm.

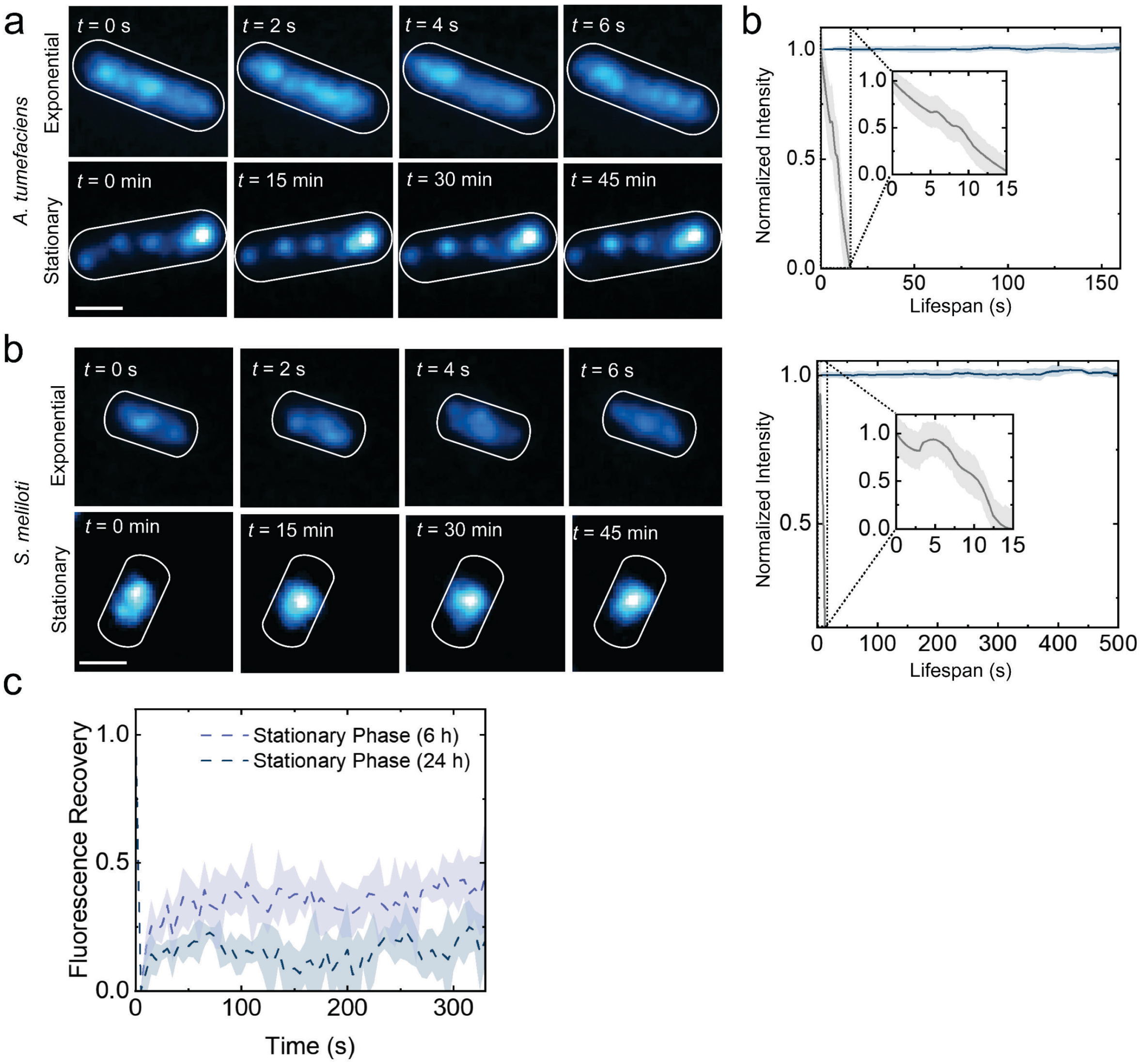


**Extended Data Fig. 7│RNE dynamics in *Agrobacterium tumefaciens* and *Sinorhizobium meliloti* during exponential and stationary growth phases.** (a) Lifespans of condensates in exponential and stationary growth phases of *A. tumefaciens*. Left: Representative images of RNE-YFP (blue) inside *A. tumefaciens* cells (white outlines) in each growth phase as a function of imaging time, *t*. Right: decay profiles of photobleaching-corrected focus intensities. Analyses include *N* = 53 foci (exponential) and *N* = 65 foci (stationary). (b) Lifespans of condensates in exponential and stationary growth phases of *S. meliloti*. Left: Representative images of RNE-YFP (blue) inside *S. meliloti* cells (white outlines) in each growth phase as a function of imaging time, *t*. Right: decay profiles of photobleaching-corrected focus intensities. Analyses include *N* = 26 foci (exponential) and *N* = 31 foci (stationary). (c) Fluorescence recovery after photobleaching (FRAP) of RNE-YFP in *A. tumefaciens* cells at early (6 h) and late (24 h) stationary phase. Scale bars in (a) and (b): 1 μm.

Methods

### Methods

#### Plasmid and bacterial strains and construction

*JS769 rne::rneΔCTD specR*

1 kB regions upstream and downstream of the RNaseE NTD were PCR-amplified from *Caulobacter crescentus* NA1000 using primers Ups F and Ups R, and primers Down F and Down R. The pNTPS138 vector[1] was PCR-amplified using primers pNTPS F and pNTPS R. The DNA fragments were run on a 1% agarose gel and extracted using a GeneJet Gel Extraction kit. The vector sample was then subjected to Dpn1 treatment followed by column purification using the GeneJET PCR Purification Kit. The 1 kB insert regions were then assembled into the pNTPS138 vector via Gibson Assembly (NEB) and transformed into chemically competent *Escherichia coli* DH10B cells and plated on LB agar supplemented with kanamycin (Kan) (30 µg/mL) for selection. Colonies exhibiting resistance to Kan were screened for the presence of the insert by PCR and verified by Sanger sequencing (Genewiz).

The spectinomycin (Spec) resistance cassette was amplified from pGFP-1[1] using primers Spec F and Spec R, while the pNPTS138 RNE-ΔCTD plasmid vector was PCR-amplified using primers ΔCTD F and ΔCTD R. The Spec resistance cassette was assembled into the pNTPS138 vector via Gibson Assembly mastermix. The assembly mixture was transformed into chemically competent *E. coli* DH10B cells and plated on LB agar supplemented with Spec (50 µg/mL) and Kan for selection. Colonies exhibiting resistance to both Spec and Kan were screened for the presence of the insert by PCR and verified by Sanger sequencing (Genewiz).

For the RNE-ΔCTD strain, the purified pNTPS138 RNE-ΔCTD SpecR plasmid was introduced into NA1000 cells via tri-parental mating and plated on PYE agar plates supplemented with Nalidixic acid (Nal) + Spec + Kan (20 µg/mL Nal; 100 µg/mL Spec; 25 µg/mL Kan). The resulting *kanR*/*specR* colonies were grown in PYE without antibiotics and then counter-selected on PYE + Spec + 3% sucrose plates. SpecR and Kan-sensitive colonies were identified and screened by PCR to confirm the deletion of the CTD on the chromosome.

*JS283 spmX::ms2 96× array vanA::Ms2-DM-mChy rne::rne-msfGFP specR/kanR/gentR*

The *spmX::ms2* 96× arrayC-1 plasmid was generated by amplifying the last 500 bp of the *spmX* gene using the primers pspmX F and pspmX R. The resulting PCR product was column-purified, digested with KpnI and EcoRI, and ligated into pFlgC-1. Resulting *specR* colonies were screened for the spmX insert which was validated by Sanger sequencing (Genewiz). The resulting plasmid was transformed into NA1000 and selected on PYE Spec plates. A phage lysate of strain *JS25*[2] (containing *vanA*::Ms2-DM-mChyC-2) was transduced and selected on PYE Kan/Spec plates. Finally, a phage lysate of *JS87*[2] (containing *rne*::*rne-msfGFPC*-4) was transduced, and colonies were selected on PYE Kan/Spec/Gent.

*JS287 rsaA::ms2 96× array vanA::Ms2-DM-mChy rne::rne-msfGFP specR/kanR/gentR*

This strain was generated in reference [2].

*JS775 rne::rne-mEos3.2C-1 SpecR*

mEos3.2 was purchased as an IDT G-block and digested with NcoI and NheI, then PCR-purified. The mEos3.2 insert was ligated with pYFPC-1, which was also digested with NcoI and NheI and gel-purified following restriction digestion. The resulting plasmid, pmEos3.2C-1, was transformed into DH10B cells and selected on LB Spec plates, followed by Sanger sequencing for verification. Plasmid prne-msfGFPC-1 was digested with EcoRI and NheI and gel-purified. mEos3.2 was digested from pmEos3.2C-1 with EcoRI and NheI, and the mEos3.2 insert and prneC-1 vector were ligated and selected on LB Spec plates. The resulting colonies were screened for the mEos3.2 insert, and the DNA sequence was confirmed by Sanger sequencing (Genewiz). The final prne-mEos3.2C-1 plasmid was transformed into NA1000 cells and selected on PYE Spec plates.

*JS776 rne::rne-ΔCTD-mEos3.2 SpecR*

The NTD of RNase E was PCR-amplified using primers HY118F and HY118R. The insert was PCR purified and digested with NdeI and EcoRI. The pmEos3.2C-1 plasmid was used as a template to generate the mEos3.2 vector using primers HY117F and HY115R. The DNA fragments were run on a 1% agarose gel and were gel extracted using a GeneJet Gel Extraction kit. The vector sample was then subjected to Dpn1 treatment followed by column purification using the GeneJET PCR Purification Kit. The RNE NTD insert fragment was then assembled into the mEos3.2C-1 vector via Gibson Assembly (NEB) and transformed into chemically competent *E. coli* DH10B cells and plated on LB agar supplemented with Spec (50 µg/mL) . The resulting SpecR colonies were grown in liquid LB + Spec (50 µg/mL) and miniprepped using the GeneJET Plasmid Miniprep Kit. The plasmid clones were screened via restriction digestion (Nde1). pRNE*ΔCTD*-mEos3.2C-1 plasmid sequencing verification was done by whole plasmid sequencing (Genewiz).

*JS770 spmX::spmX-mEos3.2 specR*

SpmX was PCR-amplified from NA1000 cells using primers HY116F and HY116R. The pmEos3.2C-1 plasmid was used as a template to generate the mEos3.2 vector using primers HY115F and HY115R. The DNA fragments were run on a 1% agarose gel and were gel extracted using a GeneJet Gel Extraction kit. The vector sample was then subjected to Dpn1 treatment followed by column purification using the GeneJET PCR Purification Kit. The SpmX insert fragment was then assembled into the mEos3.2C-1 vector via Gibson Assembly (NEB) and transformed into chemically competent *E. coli* DH10B cells and plated on LB agar supplemented with Spec (50 µg/mL). The resulting SpecR colonies were grown in liquid LB + Spec (50 µg/mL) and miniprepped using the GeneJET Plasmid Miniprep Kit. The plasmid clones were screened via restriction digestion (Nde1). pSpmx-mEos3.2C-1 plasmid sequencing verification was done by whole plasmid sequencing (Genewiz).

For the *spmX::spmX-mEos3.2 specR* strain, the purified pSpmx-mEos3.2C-1 plasmid was introduced into NA1000 cells via tri-parental mating and plated on PYE+ Nal + Spec (20 µg/mL Nal; 100 µg/mL Spec) plates. The resulting SpecR colonies were grown in PYE + Spec (25 µg/mL Spec) liquid media.

*JS804 rne::rne-ΔDBS-mEos3.2 SpecR*

The CTD (*ΔDBS)* of RNase E was PCR-amplified from pRNE(*ΔDBS)C-1*[3] using primers HY120F and HY120R. The pmEos3.2C-1 plasmid was used as a template to generate the mEos3.2 vector using primers HY119F and HY119R. The DNA fragments were run on a 1% agarose gel and were gel extracted using a GeneJet Gel Extraction kit. The vector sample was then subjected to Dpn1 treatment followed by column purification using the GeneJET PCR Purification Kit. The RNE CTD (*ΔDBS)* insert fragment was then assembled into the mEos3.2C-1 vector via Gibson Assembly (NEB) and transformed into chemically competent *E. coli* DH10B cells and plated on LB agar supplemented with Spec (50 µg/mL). The resulting SpecR colonies were grown in liquid LB + Spec (50 µg/mL) and miniprepped using the GeneJET Plasmid Miniprep Kit. The plasmid clones were screened via restriction digestion (Nde1). pRNE*CTD*(*ΔDBS)*-mEos3.2C-1 plasmid sequencing verification was done by whole plasmid sequencing (Genewiz).

For the *rne::rne-ΔDBS-mEos3.2 SpecR* strain, the purified pRNE*ΔDBS*-mEos3.2C-1 plasmid was introduced into NA1000 cells via tri-parental mating and plated on PYE+ Nal + Spec (20 µg/mL Nal; 100 µg/mL Spec) plates. The resulting SpecR colonies were grown in PYE + Spec (25 µg/mL) liquid media.

*JS805 hfq::tet rne::rne-mEos3.2C-1 specR/tetR*

Phage lysate from strain *JS775* harboring pRNE-mEos3.2C-1 was transduced into CJW5477[4] cells and selected on PYE+ Spec + Tet plates (100 µg/mL Spec; 2 µg/mL Tet). The resulting SpecR/TetR colonies were grown in PYE + Spec + Tet (25 µg/mL Spec; 1 µg/mL Tet) liquid media.

JS806 *NA1000 rne::rneΔCTD Kan$^R$*

The primer pair HY1'F and HY1'R were initially phosphorylated at their 5' ends via T4 PNK. The T4 PNK reaction was incubated at 37°C for 1 hour. The T4 PNK enzyme was then heat-inactivated by incubating the sample at 65°C for 20 minutes. The pRNE(NTD)-Apex2-FlgC-2[5] plasmid was used as a template for an inverse PCR reaction using the phosphorylated primers HY1'F and HY1'R. The DNA fragment was run on a 1% agarose gel and gel extracted using a GeneJet Gel Extraction kit. The sample was then subjected to Dpn1 treatment followed by column purification using the GeneJET PCR Purification Kit. The DNA fragment was self-ligated using T4 DNA ligase and transformed into chemically competent *E. coli* DH10B cells and plated on LB agar supplemented with Kan (50 µg/mL). The resulting KanR colonies were grown in liquid LB + Kan (30 µg/mL) and miniprepped using the GeneJET Plasmid Miniprep Kit. The plasmid clones were screened via restriction digestion (Nde1). pRNE*ΔCTD*-C-2 plasmid sequencing verification was done by whole plasmid sequencing (Genewiz).

For the *rne::rneΔCTD KanR* strain, the purified pRNE*ΔCTD*-C-2 plasmid was introduced into NA1000 cells via tri-parental mating and plated on PYE+ Nal + Kan (20 µg/mL Nal; 25 µg/mL Kan) plates. The resulting KanR colonies were grown in PYE + Kan (5 µg/mL) liquid media.

JS807 *NA1000 rne::rneΔCTD spmX::spmX-mEos3.2 specR Kan$^R$*

Phage lysate from strain *JS806* harboring pRNE*ΔCTD*-C-2 was transduced into JS770 cells and selected on PYE+ Spec + Kan (100 μg/mL Spec; 25 μg/mL Kan) plates. The resulting SpecR/KanR colonies were grown in PYE + Spec + Kan (25 μg/mL Spec; 5 μg/mL Kan) liquid media.

*JS401 rne:rne-eYFP gentR*

peYFPC-4[1] and pRNE-eYFPC-1[6] were digested with Nde1 and Kpn1. The DNA fragment was run on a 1% agarose gel and was gel-extracted using a GeneJet Gel Extraction kit. The RNE-eYFP insert was ligated into the peYFPC-4 vector using T4 DNA ligase. The ligation reaction was then used to transform pRNE-eYFPC-4 into DH10B *E. coli* cells. The cells were selected on LB + Gent (30 μg/mL) plates, and the resulting GentR colonies were grown in liquid LB + Gent (15 μg/mL) and miniprepped using the GeneJET Plasmid Miniprep Kit. The plasmid clones were screened for the presence of the insert by PCR and verified by Sanger sequencing (Genewiz).

For the *rne:rne-eYFP gentR* strain, the purified pRNE-eYFPC-4 plasmid was transformed into NA1000 cells by electroporation and selected on PYE + Gent (5 μg/mL) plates. The resulting *gentR* colonies were grown in PYE + Gent (0.5 μg/mL) liquid media.

*JS771 acnA::acnA-PAmChy specR*

PAmCherry was PCR-amplified from our RNase E-PAmCherry[6] construct using primers HY24F and HY24R. The CFPC-1 vector[1] was PCR-amplified using primers HY23F and HY23R, removing the CFP gene. The DNA fragments were run on a 1% agarose gel and were gel extracted using a GeneJet Gel Extraction kit. The vector sample was then subjected to Dpn1 treatment followed by column purification using the GeneJET PCR Purification Kit. PAmCherry was then assembled into the vector via Gibson Assembly and transformed into DH10B *E. coli* cells and selected on LB + Spec plates. Using the GeneJET Plasmid Miniprep Kit, the SpecR colonies obtained were grown in liquid LB + Spec and miniprepped and screened via restriction digestion (Nde1). PAmCherryC-1 plasmid sequencing verification was done by Sanger sequencing (Genewiz).

Aconitase was PCR-amplified from NA1000 cells using primers HY47F and HY47R. The PAmCherryC-1 vector was PCR amplified using primers HY48F and HY48R. The DNA fragments were run on a 1% agarose gel and were gel extracted using a GeneJet Gel Extraction kit. The vector sample was then subjected to Dpn1 treatment followed by column purification using the GeneJET PCR Purification Kit. Aconitase was then assembled into the PAmCherry vector via Gibson Assembly and transformed into DH10B *E. coli* cells and selected on LB + Spec plates. Using the GeneJET Plasmid Miniprep Kit, the *specR* colonies obtained were grown in liquid LB + Spec and mini-prepped and screened via restriction digestion (Nde1). PAmCherryC-1 plasmid sequencing verification was done by Sanger sequencing (Genewiz).

To generate the NA1000 *acnA::acnA-PAmChy specR* strain, the purified aconitase-PAmCherryC-1 plasmid was mated into NA1000 cells via tri-parental mating and plated on PYE + Nal + Spec plates. The resulting *specR* colonies were grown in PYE + Spec liquid media.

*JS772 acnA::acnA-PAmChy rne::rne-eYFP specR gentR*

Phage lysate from strain *JS401* harboring pRNE-eYFPC-4 was transduced into *JS771* cells and selected on PYE+ Spec + Gent plates (100 µg/mL Spec concentration; 5 µg/mL Gent concentration). The resulting SpecR/GentR colonies were grown in PYE + Spec + Gent (25 µg/mL Spec concentration; 0.5 µg/mL Gent concentration) liquid media.

*JS773 rhlB::rhlB-PAmChy SpecR*

RhlB was PCR-amplified from NA1000 cells using primers HY49F and HY49R. The PAmCherryC-1 vector was PCR amplified using primers HY50F and HY50R. The DNA fragments were run on a 1% agarose gel and gel extracted using a GeneJet Gel Extraction kit. The vector sample was then subjected to Dpn1 treatment followed by column purification using the GeneJET PCR Purification Kit. RhlB was then assembled into the PAmCherry vector via Gibson Assembly and transformed into DH10B *E. coli* cells and selected on LB + Spec plates. Using the GeneJET Plasmid Miniprep Kit, the SpecR colonies obtained were grown in liquid LB + Spec and miniprepped and screened via restriction digestion (Nde1). PAmCherryC-1 plasmid sequencing verification was done by Sanger sequencing (Genewiz).

To generate NA1000 *rhlB::rhlB-PAmChy SpecR*, the purified rhlB-PAmCherryC-1 plasmid was mated into NA1000 cells via tri-parental mating and plated on PYE + Nal + Spec plates. The resulting SpecR colonies were grown in PYE + Spec media.

*JS774 rhlB::rhlB-PAmChy rne::rne-eYFP specR gentR*

Phage lysate from strain *JS401* harboring pRNE-eYFPC-4 was transduced into *JS773* cells and selected on PYE + Spec + Gent plates. The resulting SpecR/GentR colonies were grown in PYE + Spec + Gent liquid media.

*JS777 ctrA::ms2 48× array kanR*

The ms2 48× array repeat sequence was PCR-amplified from an IDT G block into two distinct fragments with overlapping overhangs using primers HY55F and HY55R to generate fragment 1 and primers HY56F and HY54R to generate fragment 2. The YFPC-1[1] vector was PCR amplified using primers HY53F and HY53R, removing the YFP gene. The DNA fragments were run on a 1% agarose gel and gel extracted using a GeneJet Gel Extraction kit. The vector sample was then subjected to Dpn1 treatment followed by column purification using the GeneJET PCR Purification Kit. The ms2 48× array sequence was then cloned into the vector via Gibson Assembly (NEB) and transformed into DH10B *E. coli* cells and selected on LB + Kan plates. Using the GeneJET Plasmid Miniprep Kit, the kanR colonies obtained were grown in liquid LB + Kan and miniprepped and screened via restriction digestion (EcoR1). The ms2 48× array plasmid sequence verification was done by Sanger sequencing (Genewiz).

CtrA was PCR-amplified from NA1000 cells using primers HY63F and HY63R. The ms2 48× array vector was PCR amplified from the ms2 48× array plasmid using primers HY64F and HY64R. The DNA fragments were run on a 1% agarose gel and were gel extracted using a GeneJet Gel Extraction kit. The vector sample was then subjected to Dpn1 treatment followed by column purification using the GeneJET PCR Purification Kit. CtrA was then assembled into the ms2 48× array vector via Gibson Assembly and transformed into DH10B *E. coli* cells and selected on LB + Kan plates. Using the GeneJET Plasmid Miniprep Kit, the KanR colonies obtained were grown in liquid LB + Kan and miniprepped and screened via restriction digestion (EcoR1). The ctrA ms2 48× array plasmid sequence verification was done by Sanger sequencing (Genewiz).

To generate NA1000 *ctrA::ms2 48× array kanR* strain, the purified ctrA ms2 48× array plasmid was mated into NA1000 cells via tri-parental mating and plated on PYE + Nal + Kan plates. The resulting KanR colonies were grown in PYE + Kan media.

*JS778 spmX::ms2 48× array kanR*

SpmX was PCR-amplified from NA1000 cells using primers HY57F and HY57R. The ms2 48× array vector was PCR amplified from the ms2 48× array plasmid using primers HY58F and HY58R. The DNA fragments were run on a 1% agarose gel and were gel extracted using a GeneJet Gel Extraction kit. The vector sample was then subjected to Dpn1 treatment followed by column purification using the GeneJET PCR Purification Kit. SpmX was then assembled into the ms2 48× array vector via Gibson Assembly and transformed into DH10B *E. coli* cells and selected on LB + Kan plates. Using the GeneJET Plasmid Miniprep Kit, the KanR colonies obtained were grown in liquid LB + Kan and miniprepped and screened via restriction digestion (EcoR1). The spmX ms2 48× array plasmid sequence verification was done by Sanger sequencing (Genewiz).

To generate NA1000 *spmX::ms2 48× array kanR* strain*,* the purified spmX ms2 48× array plasmid was mated into NA1000 cells via tri-parental mating and plated on PYE + Nal + Kan plates. The resulting KanR colonies were grown in PYE + Kan media.

*JS779 rsaA::ms2 48× array kanR*

RsaA was PCR-amplified from NA1000 cells using primers HY61F and HY61R. The ms2 48× array vector was PCR amplified from the ms2 48× array plasmid using primers HY62F and HY62R. The DNA fragments were run on a 1% agarose gel and were gel extracted using a GeneJet Gel Extraction kit. The vector sample was then subjected to Dpn1 treatment followed by column purification using the GeneJET PCR Purification Kit. RsaA was then assembled into the ms2 48× array vector via Gibson Assembly and transformed into DH10B *E. coli* cells and selected on LB + Kan plates. Using the GeneJET Plasmid Miniprep Kit, the KanR colonies obtained were grown in liquid LB + Kan and miniprepped and screened via restriction digestion (EcoR1). The rsaA ms2 48× array plasmid sequence verification was done by Sanger sequencing (Genewiz).

To generate NA1000 *rsaA::ms2 48× array kanR* strain*,* the purified rsaA ms2 48× array plasmid was mated into NA1000 cells via tri-parental mating and plated on PYE + Nal + Kan plates. The resulting KanR colonies were grown in PYE + Kan media.

*JS780 ctrA::ms2 48× array vanA::Ms2-DM-mChy rne::rne-msfGFP specR/kanR/gentR*

Phage lysate from strain *JS777* harboring ctrA ms2 48× array KanR was transduced into *JS285* cells and selected on PYE + Spec + Gent + Kan plates. The resulting SpecR/GentR/KanR colonies were grown in PYE + Spec + Gent + Kan liquid media.

*JS789 spmX::ms2 48× array vanA::Ms2-DM-mChy rne::rne-msfGFP specR/kanR/gentR*

Phage lysate from strain *JS778* harboring spmX ms2 48× array KanR was transduced into *JS285* cells and selected on PYE + Spec + Gent + Kan plates. The resulting SpecR/GentR/KanR colonies were grown in PYE + Spec + Gent + Kan liquid media.

*JS790 rsaA::ms2 48× array vanA::Ms2-DM-mChy rne::rne-msfGFP specR/kanR/gentR*

Phage lysate from strain *JS779* harboring rsaA ms2 48× array KanR was transduced into *JS285* cells and selected on PYE + Spec + Gent + Kan plates. The resulting SpecR/GentR/KanR colonies were grown in PYE + Spec + Gent + Kan liquid media.

**Cell growth and treatment**

All *C. crescentus* strains used in this study were derived from the wild-type strain NA1000. Cultures were grown at 28 °C with shaking at 250 rpm in peptone-yeast extract (PYE) medium or M2 minimal medium supplemented with 0.2% glucose (M2G).[7] Where necessary, antibiotics were added to the media at the following concentrations: gentamycin (0.5 μg/mL), kanamycin (5 μg/mL), spectinomycin (25 μg/mL), and streptomycin (25 μg/mL). Experiments were performed during either the exponential phase ($OD_{600}$ ~ 0.7) or stationary phase (24 hours after reaching an $OD_{600}$ of 1.2). For acute treatments, exponential phase cells were treated with puromycin (150 μg/mL) for 30 minutes, CCCP (100 μM) in DMSO for 20 minutes, or a combination of chloramphenicol (200 μg/mL) and CCCP, or 10% ethanol. For the combination of chloramphenicol and CCCP, the chloramphenicol was added 1 minute before adding the CCCP. Stress treatments were conducted by exposing the cells to specific stress conditions for the indicated durations, followed by spotting 2 μL of culture onto a 2% (w/v) agarose pad for imaging. All imaging was conducted at 28 °C using an objective heater (Bioptechs).

*Agrobacterium tumefaciens* and *Sinorhizobium meliloti* were cultured in PYE medium supplemented with gentamycin (50 μg/mL) at 28 °C with shaking at 250 rpm. Experiments with these strains were conducted during the exponential phase ($OD_{600}$ 0.5-0.7) or stationary phase (36 hours post-inoculation).

**Liquid growth curves**

The NA1000 and RNE-ΔCTD strains were cultured in PYE medium at 28 °C to either exponential ($OD_{600}$ ~ 0.6) or stationary phase (24 hours after $OD_{600}$ reached 1.2). Cells were subsequently diluted in fresh PYE medium to an $OD_{600}$ of 0.05. $OD_{600}$ was measured approximately every hour. $OD_{600}$ was measured in a cuvette using a Biochrom WPA CO8000 cell density meter at 600 nm. For the RNE-ΔCTD strain, Spec (25 mg/mL) was added to the initial cultures of the RNE-ΔCTD strain. Growth rates were extracted by fitting the growth curves with a single-exponential function.

**Biorthogonal Non-Canonical Amino Acid Tagging (BONCAT)**

BONCAT was performed on NA1000 cells across various growth phases, treated with 10% ethanol after 30 minutes and 2 hours, and CCCP (100 µM for 20 min). Control samples included exponential phase cultures without azidohomoalanine (AHA) or with 100 µg/mL chloramphenicol. For the assay, 1.5 mL of cells were incubated with 200 µM of AHA for 30 minutes at 28 °C. Post-incubation, cells were washed once with 500 µL PBS, then fixed in 100 µL of 4% formaldehyde for 30 minutes at 28 °C. Cells were subsequently washed once with 500 µL PBS, then resuspended in 100 µL ice-cold 70% ethanol and incubated on ice for 30 minutes. After another PBS wash, cells were resuspended in 300 µL 0.1 mM iodoacetamide and incubated at 28 °C for 30 minutes. Following a final wash, cells were resuspended in a freshly prepared Alkyne-Cy5 cocktail (50 µM $CuSO_4$, 250 µM tris(3-hydroxypropyltriazolylmethyl)amine (THPTA), 25 µM Alkyne-Cy5, 1 mM Aminoguanidine HCl, 2.5 mM sodium ascorbate in PBS) and incubated at room temperature for 30 minutes. The cocktail was chilled on ice for 10 minutes prior to application. After two washes in PBS, cells were resuspended in 20-200 µL PBS, depending on the pellet size, and imaged using the fluorescence channel with a 100 ms exposure time. Fluorescence intensity was calculated using the MicrobeJ[8] software tool.

**Relative intracellular ATP level measurement**

Intracellular ATP levels were quantified using a luciferase-based assay (BacTiter-Glo), following the manufacturer's protocol (Promega). Luminescence was measured using a microplate reader (Molecular Devices SpectraMax iD3) with white-walled luminescence 96-well plates to minimize crosstalk between wells. All cultures were adjusted to the same optical density.

**mRNA half-life measurement**

*C. crescentus* NA1000 cells were grown to exponential phase ($OD_{600}$ ~ 0.3-0.6), early stationary phase (OD = 1.2), or late stationary phase (72 hours post-OD 1.2). Additionally, JS263 cells were grown to exponential and late stationary phase. At time zero (before rifampicin addition), 1 mL of cells were collected and immediately vortexed with 2 mL of RNAprotect Bacterial reagent. Additional 1-mL aliquots were taken at specific time intervals (1, 2, 4, 8, 16, and 32 minutes) after the addition of rifampicin (200 µg/mL) and treated similarly. TRIzol was pre-warmed to 65 °C. Cells in RNAprotect were pelleted by centrifugation at 8000 rpm for 2 minutes, resuspended in 1 mL of pre-warmed TRIzol, and incubated at 65 °C for 10 minutes. Subsequently, 200 µL of chloroform was added, and the mixture was incubated at room temperature for 5 minutes before centrifuging at maximum speed for 10 minutes. The aqueous phase was transferred to a new tube, followed by an additional chloroform extraction with 500 µL. The upper aqueous phase was again collected, to which 2 µL of glycogen and 700 µL of ice-cold isopropanol were added, and samples were incubated overnight at −80 °C. The next day, samples were centrifuged at maximum speed at 4 °C for 1 hour. The supernatant was discarded, and the pellet was washed with 800 µL of ice-cold 80% ethanol, followed by another centrifugation step. The supernatant was discarded, then the pellet was air-dried and resuspended in 50 µL of resuspension buffer (10 mM Tris-HCl, pH 7.0, 0.1 mM EDTA).

For RT-qPCR, a master mix was prepared containing 1× Luna Universal One-Step Reaction Mix, 1× Luna WarmStart RT Enzyme Mix, 0.4 µM each of forward and reverse primers, and milliQ water. The following primer pairs were used to test the half-lives of each these RNAs: tmRNA/*ssRA* (5' ssRA F tccggcttcggccgaacta and 5' ssRA R tggagccgccgggaatcg), *ctrA* (CtrA F actgatgctgaagtctgaagg and CtrA R gattgaggtcgagcaggataag), *spmX* (SpmX F ctgctgctctatgatctcatctc and SpmX R gaagttgtcgaggccgatatt), *rne* (RNE F cgcgaaatccgtgatcgt and RNE R cgcgtctcgaacactatctaatc), and 5S (5S F ccattccgaactcggtcgttaag and 5S R tggcggcgacctactct). RNA samples (100 ng/µL) were aliquoted into a 96-well plate, mixed with the master mix, and briefly centrifuged. qRT-PCR reactions were performed using a QuantStudio Real-Time PCR system. RNA decay rates were calculated by fitting a linear curve to the natural logarithm of the fraction of RNA remaining at each time point. The quantity of RNA at each time point was normalized to the amount detected in the time zero sample as 100%. The mRNA half-life was determined using linear regression of the ln (percentage of RNA remaining) at each time point of RNA extraction and converted the slope to a half-life using ($t_{1/2}$ = -ln(2)/slope).

To assess full-length mRNA levels at the stationary phase, reverse transcription reactions were performed using extracted total RNA as the template, by applying the Invitrogen first-strand cDNA synthesis technique with Superscript III. The RNA templates used were extracted from NA1000 cells from time points 0 and 16 minutes in both log and stationary phase. 2 pmol of *ctrA* RT FL R primer (tcaggcggcgttaacctgctc) and 1 µL of dNTP mix (10 mM each of dATP, dGTP, dCTP, and dTTP) were combined with 1 ug of total RNA. The sample was heated to 65 °C for 5 mins and then incubated on ice for 1-2 mins. Afterward,1 µL of DTT (0.1 M), 4 µL of 5× First-strand buffer, 1 µL of SUPERase-In RNase inhibitor (20 units/ µL), and 1 µL Superscript III (200 Units/ µL) were added to the mixture, and the samples were incubated at 55 °C for 60 min. The samples were heated at 70 °C for 15 min to inactivate the reaction. PCR was then performed using the resulting cDNA as a template for amplification. A 50 µL PCR reaction was conducted by adding 25 µL of Betaine (5 M), 12 µL of 5× HF-buffer, 7.5 µL of milliQ water, 3 µL of ctrA RT FL R primer (10 µM), 3 µL of ctrA RT FL F primer (10 µM) (atgcgcgtactgttgatcgagg), 1 µL dNTP mix (10 mM), 2 µL of cDNA from the reverse transcriptase reaction, and 0.5 µL of Phusion enzyme (2 Units/ µL). 25 PCR cycles were performed on the log phase samples, while 30 PCR cycles were performed on the stationary phase samples. 25 µL from each PCR sample was then run on a 1% agarose gel and imaged post ethidium bromide staining.

The same experiment was repeated utilizing 5S RT FL primers (5S RT FL F gacctggtggctatgccgg; 5S RT FL R gacctggcggcgacctac) to measure the full-length mRNA levels of 5S ribosomal RNA in log and stationary phase. 10 PCR cycles were performed on both log and stationary phase samples.

**Time-lapse BR-body disassembly fluorescent imaging**

*JS87 cells* were cultured overnight in PYE + Gent medium and then inoculated into M2G medium. The cells were incubated in M2G at 28 °C for 24 hours post cell reaching an OD of 1.2. 1 µL of cells was immobilized on a 1.5 % (w/v) agarose pad made from fresh M2G medium and sealed with melted wax. To take the images, a Nikon Eclipse NI-E fitted with a CoolSNAP MYO-CCD camera and a 100× Oil CFI Plan Fluor (Nikon) objective was controlled using Nikon Elements software. The same field of view was imaged under the phase-contrast and GFP channels every 30 minutes over the span of 6 hours.

**Wide-field fluorescence and phase-contrast microscopy**

Wide-field fluorescence and phase-contrast images were acquired with a Nikon Ti2-E motorized inverted microscope equipped with a SOLA 365 LED light source, a 100× oil immersion objective, and a Hamamatsu Orca-Flash 4.0 LTS camera. The system was controlled via NIS Elements software. Fluorescent protein fusions were imaged using specific filter sets from Chroma: mEos3.2 and YFP fusions were visualized with a YFP filter set (C-FL YFP, hard coat, high signal-to-noise ratio, zero shift, excitation: 500 ± 10 nm, emission: 535 ± 15 nm, dichroic mirror: 515 nm); GFP fusions with a GFP filter set (C-FL GFP, hard coat, high S/N, zero shift, excitation: 436 ± 10 nm, emission: 480 ± 20 nm, dichroic mirror: 455 nm); mCherry fusions with a Texas Red filter set (C-FL Texas Red, hard coat, high S/N, zero shift, excitation: 560 ± 20 nm, emission: 630 ± 37.5 nm, dichroic mirror: 585 nm); and CFP fusions with a CFP filter set (C-FL CFP, hard coat, high S/N, zero shift, excitation: 436 ± 20 nm, emission: 480 ± 40 nm, dichroic mirror: 455 nm).

**Live-cell fluorescence recovery after photobleaching (FRAP)**

FRAP was performed using a Nikon Ti2-E motorized inverted microscope, equipped with a SOLA 365 LED source, a 100× objective (oil immersion), and a Hamamatsu Orca-Flash 4.0 LTS camera. Control images were taken prior to bleaching, and subsequently, the regions of interest (ROIs) were bleached with a laser at 405 nm and 30% power (~15 mW). Image acquisition was performed with 300-ms integration times. More than ten different ROIs were chosen per sample. Images were analyzed with Fiji[9]: the off-cell background signal was subtracted from the bleached ROI and an unbleached ROI of the same size within the cell. The resulting intensity in the ROI was normalized such that the intensity before bleaching equals one and the first frame after bleaching equals zero.

**Live-cell single-molecule fluorescence microscopy**

*Caulobacter crescentus* cells expressing RNE-mEos3.2, RNE-YFP/Aconitase-PAmChy, and RNE-YFP/RhlB-PAmChy fusions were cultured overnight in 75 mL of PYE medium until they reached the exponential phase ($OD_{600}$ ~ 0.7). The following day, the cultures were diluted 1:100 into 75 mL of M2G medium and grown to an $OD_{600}$ ~ 0.7 for imaging during the exponential phase and stress experiments, or for 24 hours post-$OD_{600}$ ~ 1.2 for imaging during the stationary phase. The cells were immobilized on 2% (w/v) agarose pads made from spent M2G medium. To prepare the spent medium, an aliquot of the culture was centrifuged at 6000 rpm for 8 minutes, and the supernatant was filtered twice through a 0.22-µm syringe filter. Imaging was performed using a wide-field

Olympus IX71 inverted microscope equipped with a 100× oil immersion objective (1.40 NA). Photoconversion of mEos3.2 and PAmChy was achieved with 75- to 150-ms pulses of a 405 nm laser (Coherent Cube 405-100; 2 W/cm$^2$), followed by imaging with a 561 nm laser (Coherent Sapphire, 561-50; 0.34 kW/cm$^2$). For RNE-YFP/Aconitase-PAmChy and RNE-YFP/RhlB-PAmChy strains, RNE-YFP was imaged for 10 seconds using a 488-nm laser (Coherent Cube 488-50; 0.4 W/cm$^2$) immediately after the 561-nm imaging to locate BR-bodies. Frames were captured at 50 Hz using a 512 × 512 Photometrics Evolve EMCCD camera.

**Live-cell single-molecule data analysis**

The recorded movies were analyzed using the SMALL-LABS[10] algorithm for single-molecule localization and tracking. To classify RNE-mEos3.2 trajectories between inside or outside of BR-bodies, we accounted for the dynamic and transient nature of BR-body formation, especially in exponential phase cells. Each movie was divided into consecutive 30-second submovies, reflecting the approximate lifetime of BR-bodies in exponential phase cells under our imaging conditions. For each submovie, a composite image was generated and used to segment BR-body foci using intensity-based thresholding. Only foci containing at least three single-molecule trajectories were retained for analysis. Overlapping or ambiguous foci were excluded to ensure reliable assignment. Single-molecule trajectories were then categorized based on their spatial overlap with BR-body regions: 'In' if entirely within a focus for the entire duration, 'In/Out' if overlapping with a focus for 25% to 99% of their duration, and 'Out' if overlapping for less than 25% of their duration. This strict classification ensures that 'In' trajectories reflect on motion within the BR-body interior and are not influenced by entry or exit events. To assess whether condensate motion could affect our measurements, we analyzed BR-body motion using trajectories matched in length to those of individual RNE molecules. This analysis showed no significant difference in BR-body mobility between exponential and stationary phases. For aconitase-PAmChy, the same trajectory classification strategy was used. In exponential phase, aconitase-PAmChy itself served as the BR-body marker for segmentation. In stationary phase, BR-body masks were generated by summing 10-second 488 nm channel movies collected immediately after the 561 nm imaging, allowing consistent identification and segmentation of BR-bodies under both conditions.

To characterize the diffusion of single RNE molecules within BR-bodies, we calculated the 2D apparent diffusion coefficient ($D_{app}$) for each trajectory by fitting the mean-squared displacement (MSD) as a function of time lag ($\tau$) over the interval 20–100 ms. Ensemble-averaged diffusion coefficients were also obtained by fitting the MSD curve averaged across all trajectories. Given that RNE-mEos3.2 diffusion within BR-bodies during stationary phase is nearly indistinguishable from that observed in fixed cells, indicating motion is limited by localization precision, we did not calculate the anomalous diffusion exponent $\alpha$, as it is no longer informative of confinement under these conditions.

**Cluster analysis**

*C. crescentus* expressing RNase E-mEos3.2 was grown to $OD_{600}$ ~ 0.7 for imaging during exponential-phase and stress experiments, and for 24 hours after cells reached $OD_{600}$ ~ 1.2 for imaging during stationary phase. Prior to imaging, the cells were fixed using formaldehyde cross-linking with 1% fixation buffer followed by a 10-minute incubation at room temperature and 30 minutes incubation on ice, and finally washed three times with M2G. Before imaging, cells were resuspended in M2G and were subsequently spotted on 2% (w/v) agarose in M2G medium pads with Fluoresbrite carboxylate YG beads at a concentration of 5.0 × $10^7$ beads/mL as fiducial markers. Before tracking, movies were corrected for drift. Tracking was performed using SMALL-LABS.

Oversampling, caused by including multiple localizations of the same protein that emit for more than one frame, can create false clustering. Before analyzing clustering in fixed cells, over-sampling was removed with a radial and temporal filter. For each single molecule, a spatial filter with size of $\sqrt{{\sigma_x}^2 + {\sigma_y}^2}$ was used, where $\sigma$ is the localization precision (Supplementary Figure 10).[11] The temporal threshold was acquired by fitting the probability of finding another localization within the spatial threshold to an exponential decay and using the time constant as the threshold. To determine which class of single molecules was a cluster, Density-Based Spatial Clustering of Applications with Noise (DBSCAN)[12] was used; the constant neighborhood radius, $\varepsilon$, was set to 50 nm, and the minimum points was set to 20. The diameter of the clusters was calculated as the maximum span of each cluster. The number of localizations per cluster used for the determination of BR-body density and the fractions of localizations of RNE, aconitase, and RhlB correspond to the number of localizations after temporal and radial filtering identified with DBSCAN with the parameters reported above.

To assess the recruitment of RNA degradosome components (RNase E, aconitase, and RhlB) to BR-bodies in *C. crescentus*, we combined single-molecule tracking (SMT) with whole-cell fluorescence intensity analysis. For each protein and growth phase (exponential and stationary), we calculated the fraction of single molecules localized within BR-bodies. Three biological replicates were analyzed per condition. To account for differences in total protein expression across growth phases, we normalized the fraction of BR-body-localized trajectories by the total cellular fluorescence intensity of the respective protein. These values were obtained by summing the fluorescence signal across all frames of the SMT acquisition movies for each replicate. This normalized recruitment index reflects the proportion of total cellular protein localized to BR-bodies, enabling direct comparison of BR-body enrichment between conditions.

**RNE residence times in the BR body**

Due to the fast photobleaching of fluorescent proteins, continuous single-molecule tracking cannot accurately determine residence time, as both dissociation and photobleaching contribute to signal loss. To overcome these factors, we performed time-lapse imaging to capture the residence behavior of RNE-mEos3.2 molecules at multiple timescales. In our time-lapse imaging, every frame was captured with a 250-ms integration time, $\tau_{\text{int}}$, and a time delay, $\tau_{\text{delay}}$ = 300, 350, 750, or 1250 ms, was introduced between each pair of consecutive frames. An integration time of 250 ms was chosen to intentionally blur fast-diffusing RNE-mEos3.2 molecules not associated with BR-bodies, thereby enhancing the detection of slow-moving molecules likely localized within BR-bodies. To further ensure accurate assignment, we applied the same trajectory classification strategy used in the RNE-mEos3.2 mobility analysis to filter the resulting trajectories.

To measure the timescale of RNE residence inside the BR-bodies, we modeled the interaction between RNE and the BR-bodies as a simple association/dissociation reaction that has a forward reaction constant of $k_{diss}^{app}$. This apparent dissociation rate of RNE includes contributions from the true dissociation rate, $k_{diss}$, and the photobleaching rate, $k_{pb}$. $k_{res}^{app}$ was obtained by plotting the $(1 - CDF)$ distribution of RNE dwell times, $\tau_{measured}$, for each of the time-lapse experiments and fitting to a single-exponential function, where the extracted rate corresponds to $k_{res}^{app}$. Photobleaching of mEos3.2 only occurs when laser illumination is on (i.e., only during $\tau_{int}$) while the actual dissociation of RNE from the BR-bodies can occur at any time during the time-lapse period, $\tau_{TL} = \tau_{int} + \tau_{delay}$. These processes contribute independently to $k_{diss}^{app}$, and respectively produce the two terms on the right-hand side of:

$$k_{diss}^{app}\tau_{TL} = \tau_{TL}k_{diss} + \tau_{int}k_{pb} \quad (1)$$

As shown in Equation (1), the relationship between $k_{diss}^{app}\tau_{TL}$ and $\tau_{TL}$ is linear. Hence, the residence time, $\tau_{res}$, was extracted from the reciprocal of the slope, $k_{diss}$, of the plot of $k_{diss}^{app}\tau_{TL}$ versus $\tau_{TL}$. The linear regression also took the uncertainty of each data point from the exponential fitting into consideration and gave the final slope, $k_{diss}$, and its associated error.

**Determination of mRNA storage with the MS2 system**

Strains *JS777*, *JS778*, *JS779*, *JS780*, *JS283*, and *JS287* containing RNase E-GFP and a non-dimerizable mutant of the Ms2 coat protein fused to mCherry, along with an array of MS2 RNA hairpins fused to the 3' end of the *rsaA*, *spmX*, *ctrA*, or *rne* genes, respectively, were grown to $OD_{600}$ ~ 0.7 for imaging during exponential phase and stress experiments. These strains were grown for 24 hours after reaching $OD_{600}$ ~1.2 for imaging during stationary phase. The cells were immobilized on 2% (w/v) agarose: spent M2G medium pad and imaged using the system described above ("Wide-field fluorescence and phase contrast imaging" section), and Fluoresbrite carboxylate YG beads were added at a concentration of $5.0 \times 10^7$ beads/mL as fiducial markers. Time-lapse movies were collected using 250-ms integration times and a delay between frames of 10 and 30 seconds for exponential and stationary phases, respectively. After data collection, the movies were drift-corrected by using the drift path of the fiducial markers.

Colocalization between the RNE-GFP and the mRNA::*ms2*-mCherry foci was performed through thresholding. After correcting for photobleaching in both channels by implementing the exponential fitting method in ImageJ,[13] Gaussian fitting was used to acquire the position of the foci in the GFP channel, and an ROI of 3 × 3 pixels (210 nm × 210 nm in our imaging setup) was used to get the intensity of the foci. The focus intensity was defined as the average intensity of the ROI. The same ROI in the same position was then applied to the mCherry channel, and the intensity was measured as in the GFP channel. An intensity threshold of 2000 counts was set to determine whether or not there was a focus at the same position in the mCherry channel. Only ROIs in the mCherry channel that contained foci colocalized with the foci in the GFP channel were considered for the analysis.

Tracking was performed in foci that were colocalized in both channels. The minimum trajectory length was set to 4 frames. After tracking, the intensities of each focus in both channels were divided by the intensity of the focus at time zero (when the focus forms) to determine the intensity enhancement of each focus at each time. Recruitment of transcripts across growth phases and acute stresses was quantified using Pearson correlation coefficients (PCC). For each cell, the PCC was computed from the pixel intensities of the RNA channel and the pixel intensities of the BR-body marker channel within the segmented cell mask after background subtraction.

**Translation of *rne* upon nutrient replenishment**

Nutrients were replenished by adding fresh M2G medium to stationary phase cultures of *C. crescentus* RNE-mEos3.2 and RNE-ΔCTD-mEos3.2, while simultaneously adding rifampicin at a concentration of 75 µg/mL to inhibit transcription. Following this inhibition, the mEos3.2 fluorescence intensity was bleached using 488-nm excitation, and intensity recovery was monitored. The cells were incubated on the microscope stage with an objective heater set to 28 °C.

**Fluorescence *in-situ* hybridization (FISH)**

Cells were grown then fixed with 1% formaldehyde for 15 minutes at room temperature, followed by an additional 30 minutes on ice. The cells were harvested by centrifugation at 7500 rpm for 4 minutes. The resulting cell pellets were washed twice with ice-cold 1× PBS. The cells were then pelleted again at 7500 rpm for 4 minutes and washed three times with GTE buffer. Cell lysis was performed using 4000U of RNase-free lysozyme at 37 °C for 2 hours. Following lysis, the cells were centrifuged, and the pellet was resuspended in 100 µL of hybridization buffer (Stellaris RNA FISH) containing 1 µL of the FISH probe (Biosearch). The control involved treating the samples with RNaseA (ThermoFisher) before adding the FISH probe. The samples were incubated overnight at 42 °C. Post-hybridization, the cells were centrifuged and washed with 1 mL of wash buffer A at 42 °C for 30 minutes, followed by a wash with 1 mL of wash buffer B (Stellaris) at 42 °C for 5 minutes. Finally, the cells were resuspended in GTE buffer supplemented with 0.1% TritonX-100, mounted on slides using 2% agarose pads, and imaged in the wide-field fluorescence microscopy system described above ("Wide-field fluorescence and phase-contrast microscopy" section). The FISH tmRNA/*ssrA*-Cy3 probe used was 5'-CGGCGAACTCTTCAGCGAAGTTATCGTTCGC-3' (Integrated DNA Technologies). The probes for *ctrA*, *spmX,* and *rsaA* (Biosearch) are reported in Supplementary Table 5. The fluorophore used for *ctrA* and *spmX* was Quasar 570, and Quasar 670 was used for *rsaA*.

**Protein purification**

An overnight culture of *E. coli* BL21(DE3) transformed with the pVN030 plasmid encoding MBP-RNE CTD (461-898 aa) was reinoculated into 1.5 L of LB medium containing 30 µg/mL kanamycin and grown at 37 °C with shaking at 200 rpm until the $OD_{600}$ reached approximately 0.6. Expression of MBP-RNE CTD was induced by adding 0.5 mM IPTG, followed by incubation at 37 °C with shaking at 140 rpm for 3 hours. Cells were harvested by centrifugation at 5000 rpm for 15 minutes and resuspended in 20 mL of lysis buffer (20 mM Tris, pH 7.4, 500 mM NaCl, 10% glycerol, 10 mM imidazole, 1 mM PMSF, 2 tablets protease inhibitor cocktail, and 10 µg/mL DNase I). The cells were lysed using sonication at 4 °C, over 18 cycles of 5 seconds on and 20 seconds off. The lysate was centrifuged at 14,000 rpm for 45 minutes at 4 °C, and the supernatant was loaded onto pre-equilibrated Ni-NTA resin to bind the 6× His-MBP-RNE (461-868). The resin-bound protein was washed with 5 column volumes each of low salt buffer (20 mM Tris, pH 7.4, 150 mM NaCl, 5% glycerol, 10 mM imidazole) and high salt buffer (20 mM Tris, pH 7.4, 1000 mM NaCl, 5% glycerol, 10 mM imidazole), before elution with elution buffer (20 mM Tris, pH 7.4, 150 mM NaCl, 5% glycerol, 250 mM imidazole). The protein was concentrated using an Amicon concentrator with a 30 kDa cutoff and dialyzed against 20 mM Tris, pH 7.4, 150 mM NaCl buffer. The concentrated protein was stored at −80 °C.

Full-length GFP-RNE, encoded by the pVN059 plasmid, was purified using the same protocol as previously described.[14] The Ni-NTA affinity-purified GFP-RNE full-length was concentrated using an Amicon concentrator with a 50 kDa cutoff and further purified by size exclusion chromatography on an S200 Sephadex column to remove contaminant proteins. The final protein was eluted in 20 mM Tris, pH 7.4, 150 mM NaCl, 2% glycerol, concentrated to ~10 mg/mL, and stored at −80 °C.

TEV protease was purified using a similar protocol, except the plasmid conferred ampicillin resistance. The Ni-NTA affinity-purified TEV protease was concentrated using an Amicon concentrator with a 10 kDa cutoff and further purified by size exclusion chromatography on an S200 Sephadex column in 20 mM Tris, pH 7.4, 150 mM NaCl, and 5% glycerol. The protein was stored at −80 °C in 1 mg/mL aliquots.

***In vitro* transcription of RNase E 5' UTR**

RNase E 5' UTR was *in vitro* transcribed and purified as described previously.[15] The plasmid containing RNase E 5' UTR (pVN053) was linearized using Nhe1 restriction enzyme, which served as the template for *in vitro* transcription (IVT). The IVT reaction mixture contained 2.7 mg of linearized plasmid, 21 µg homemade T7 RNA polymerase, 2.5 mM NTPs, 1× reaction buffer (50 mM Tris-Cl pH 7.4, 15 mM MgCl2, 5mM DTT, 2 mM spermidine), 0.01 U/µL PPase in a total volume of 1000 µL and the reaction was carried out at 37 °C for 4 hours. The transcribed RNA loaded onto 7% Urea gel was extracted using phenol-chloroform and ethanol precipitation methods. The precipitated RNA was resuspended in nuclease-free water.

**pCp Cy5 labeling of RNase E 5' UTR**

*In vitro* transcribed RNase E 5' UTR (4.5 µM) was incubated with 33 µM of pCp Cy5 in a total reaction volume of 100 µL consisting of 50 units of T4 RNA ligase 1, 1mM ATP, 10% DMSO, 1× T4 RNA ligase reaction buffer for 16 hours at 16 °C. Following the reaction, T4 RNA ligase 1 was heat-inactivated at 65 °C for 15 minutes, and the reaction mixture was subjected to a phenol-choloroform extraction to remove the enzyme. The unincorporated pCp Cy5 was removed by passing the labeled RNA through a Sephadex G-50 column and elution in nuclease-free water.

***In vitro* condensate assay and RNA degradation in the condensates**

20 µM of MBP-RNase E CTD (461-868) was treated with 1 µM of TEV protease at room temperature in 20 mM Tris pH 7.4 and 100 mM NaCl in a total reaction volume of 10 µL for 45 minutes to cleave off

MBP tag from RNase E CTD. The formation of RNase E CTD condensates was verified by microscopy, followed by the addition of 1 μM GFP RNase E full-length and 0.5 μM RNase E 5' UTR Cy5. After 10 minutes, the recruitment of GFP RNase E full-length and RNase E 5' UTR into RNase E CTD condensates was verified using a Nikon Eclipse NI-E with CoolSNAP MYO-CCD camera and 100× Oil CFI Plan Fluor (Nikon) objective under GFP and Cy5 channels with 50 ms exposures. The condensates were incubated for 30 minutes, 4 hours, or 8 hours in separate tubes, followed by the addition of 200 μM $MgCl_2$ to each tube to stimulate GFP RNase E full-length endonucleolytic activity. At the end of each time point, 2 μL from the reaction was quenched with 8 μL quench buffer (20mM Tris pH 7.4, 50mM EDTA, 0.1% SDS) and 3 μL of quenched reaction was mixed with 7 μL loading buffer (95% formamide, 10mM EDTA, 0.1% SDS, 0.025% bromophenol blue, 0.025% xylene cyanol), boiled at 90 °C for 2 minutes before loaded onto 7% urea PAGE. The gel was run in a Biorad Mini-PROTEAN tetra cell system at 190 V for 45 minutes. The gel was stained with 1× SYBR gold for 20 minutes and scanned using a Thermofisher iBright imaging system.

**RNE CTD (461-868) generation and Cy5 labeling**

6×-His-MBP-RNase E CTD and TEV protease were incubated at a molar ratio of 20:1 in 20 mM Tris (pH 7.4) and 150 mM NaCl for 16 hours at 4 °C with gentle rocking. The cleavage of MBP was confirmed using SDS-PAGE. The reaction mixture was then loaded onto pre-equilibrated Ni-NTA resin and incubated for 5 minutes to bind the 6× His-MBP to the resin. The unbound RNase E CTD was collected, and its purity was assessed via SDS-PAGE. RNase E CTD was concentrated using an Amicon concentrator with a 10-kDa cutoff. Cy5 labeling was performed following previously described protocols.[15] Briefly, Cy5 NHS ester dye dissolved in DMSO was incubated with RNase E CTD in an equal molar ratio in 0.1 M sodium bicarbonate buffer (pH 8.5) at room temperature for 4 hours. Excess dye was removed by dialysis against 20 mM Tris (pH 7.4) and 150 mM NaCl using a dialysis cassette with a 3-kDa cutoff.

***In vitro* FRAP of RNE CTD condensates**

To form RNE CTD Cy5 condensates, 20 μM RNE CTD Cy5 was incubated in 20 mM Tris (pH 7.4) and 100 mM NaCl at room temperature for 20 minutes, in a total reaction volume of 10 μL. Following this incubation, 1 μM GFP-RNE full-length and 0.5 μM RNE 5' UTR were added to the condensates. The mixtures were incubated for 30 minutes, 4 hours, and 8 hours, respectively. At the end of each incubation period, 6 μL of the reaction mixture was placed onto a homemade welled slide, covered with a coverslip, and FRAP was performed using a Zeiss LSM 800 microscope. The diameters of the bleached and reference areas within the condensates were set to 2 μm. The resulting intensity in the ROI was normalized such that the intensity before bleaching equals one and the first frame after bleaching equals zero.

**Statistical analysis of distributions**

$p > 0.05$ was considered not significant. $*p \leq 0.05$, $**p \leq 0.01$, $***p \leq 0.001$ were considered significant using a two-tailed Student's t-test.

Supplementary Figures

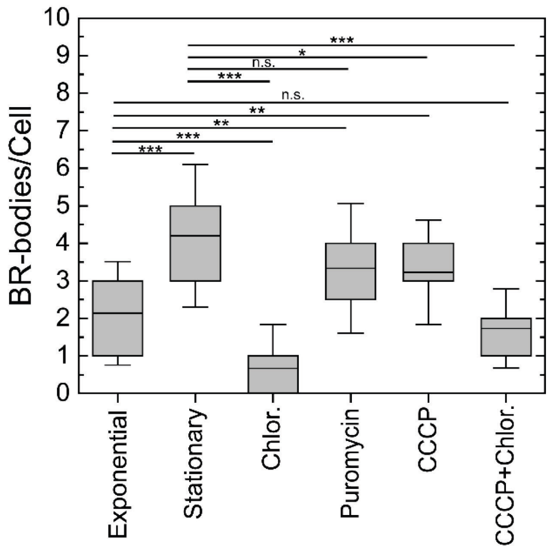


**Supplementary Figure 1.** Quantification of the number of BR-bodies per cell as a function of growth phase or acute stress. Error bars indicate the standard deviation. n.s. > 0 .05, *$p \leq 0.05$, **$p \leq 0.01$, ***$p \leq 0.001$. $n$ = 131, 109, 72, 78, 116, and 92 cells for exponential phase, stationary phase, chloramphenicol, puromycin, CCCP, and CCCP+chloramphenicol, respectively.

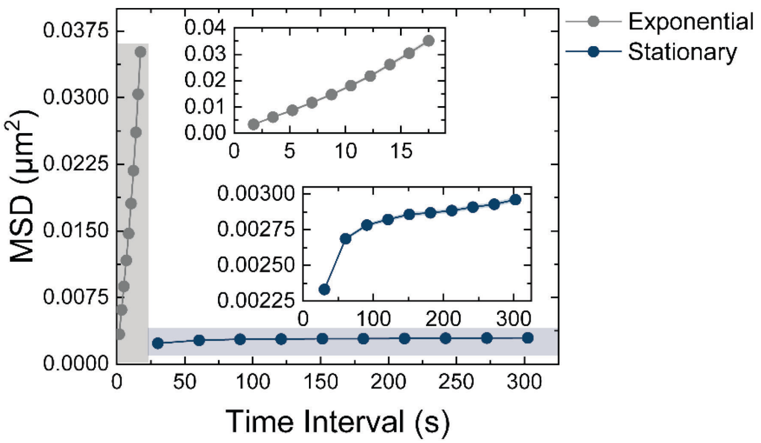


**Supplementary Figure 2.** Mean squared displacement (MSD) as a function of time interval for BR-bodies during exponential and stationary phase. BR-bodies in stationary phase cells are nearly immobile compared to those in exponential phase cells, reflecting a substantial decrease in mobility. Due to the large difference in diffusion, both curves are also shown in insets (highlighted by colored rectangles) to aid visualization on appropriate scales. Number of trajectories analyzed: 3,761 (exponential phase) and 12,265 (stationary phase).

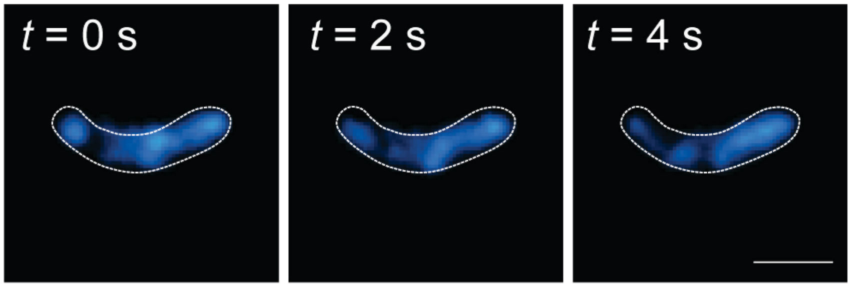


**Supplementary Figure 3.** Representative images of *C. crescentus* cells (white outlines) and RNE-mEos3.2 (blue) following chloramphenicol (200 μg/mL) treatment as a function of imaging time. Scale bar: 500 nm.

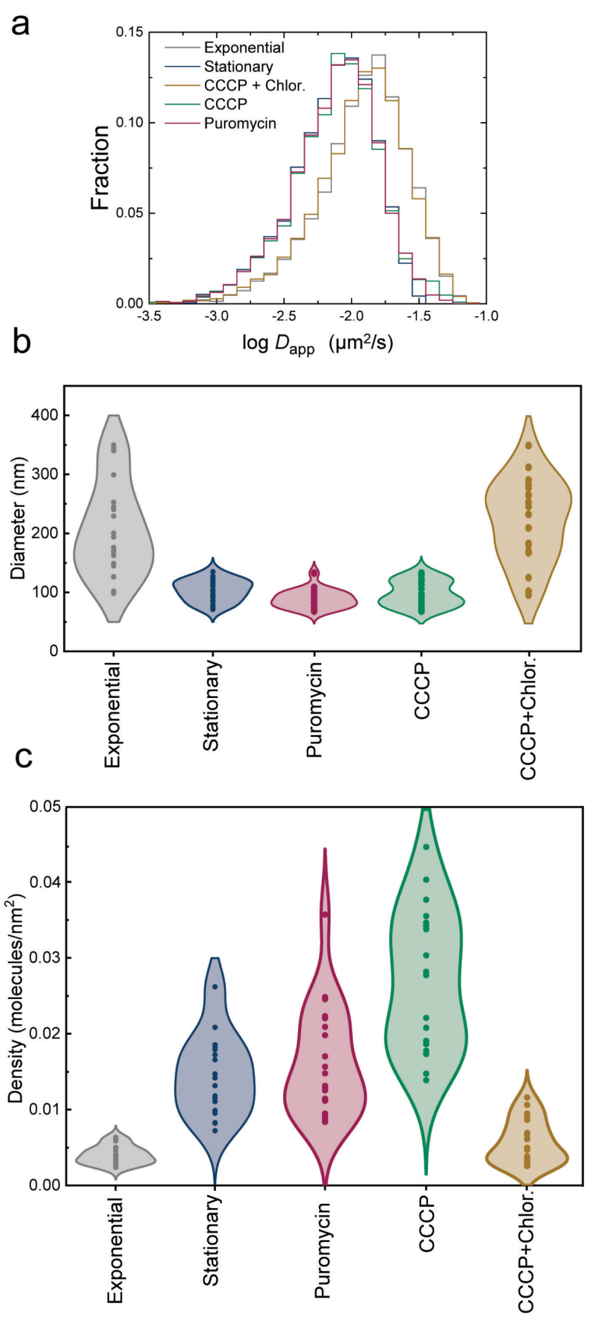


**Supplementary Figure 4.** Comparison of (a) apparent diffusion coefficient distributions, (b) BR-body diameters, and (c) BR-body molecular density across growth phases and acute stress conditions. Number of trajectories analyzed in (a): 5,001 (exponential phase), 6,347 (stationary phase), 2,126 (puromycin), 1,637 (CCCP), and 1,441 (CCCP + chloramphenicol). Number of clusters analyzed in (b) and (c): 20.

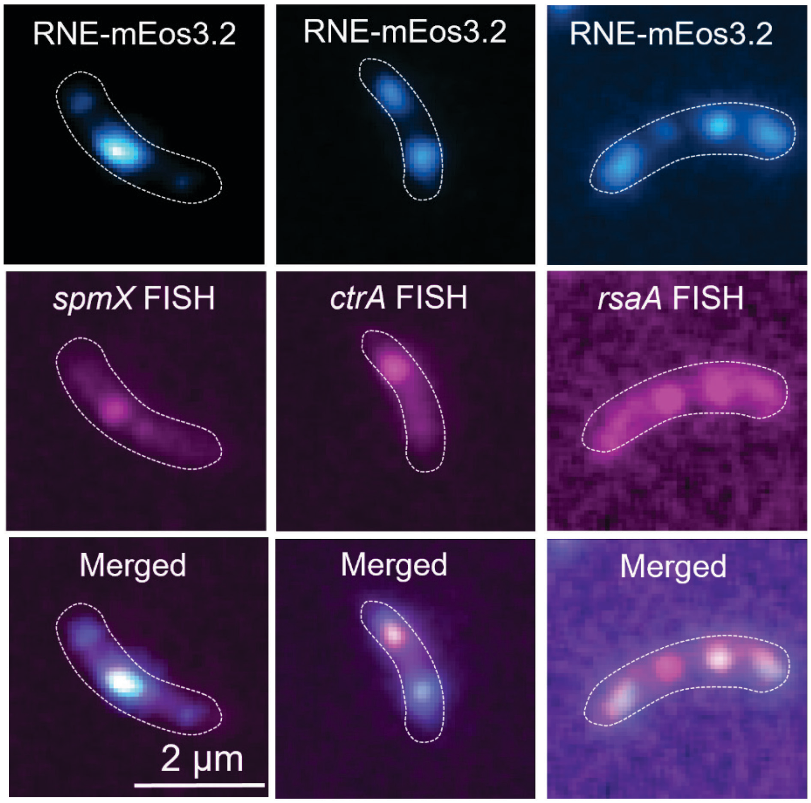


**Supplementary Figure 5.** mRNA fluorescence in situ hybridization (FISH) of selected transcripts during stationary phase. Cyan: Fluorescence images of RNE-mEos3.2 in representative *C. crescentus* cells (white outlines). Magenta: mRNA FISH images of *spmX*, *ctrA*, and *rsaA* transcripts in the same *C. crescentus* cells, respectively.

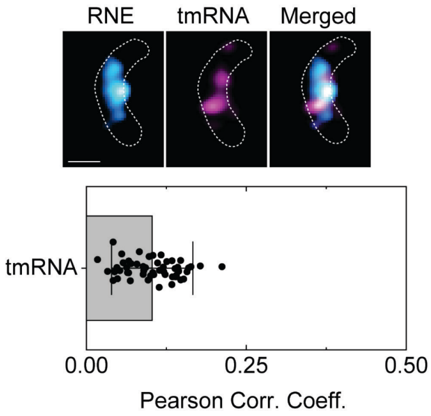


**Supplementary Figure 6.** *ssrA*/tmRNA FISH (top) and quantification of colocalization profile using the Pearson correlation coefficient (bottom). Cyan: Fluorescence images of RNE-mEos3.2 in a representative *C. crescentus* cell (white outlines). Magenta: tmRNA FISH images. Scale bar: 500 nm. Error bars indicate the standard deviation of the Pearson correlation coefficients obtained for each of 50 cells.

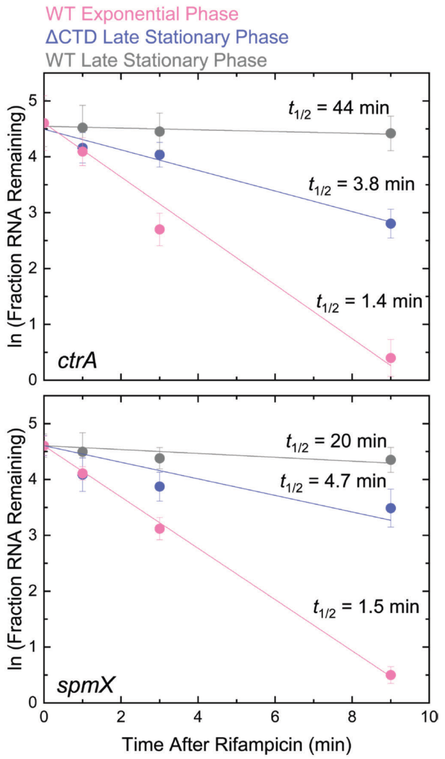


**Supplementary Figure 7.** BR-body formation is required for robust mRNA stabilization during stationary phase. Fraction of RNA remaining over time as measured by mRNA FISH for *ctrA* and *spmX* transcripts in wild-type exponential phase (pink), wild-type late stationary phase (gray), and RNE-ΔCTD late stationary phase (blue). Transcripts are rapidly degraded in exponential phase and in the absence of BR-bodies (RNE-ΔCTD) but are significantly stabilized in wild-type stationary phase, indicating that BR-body formation is necessary for transcript protection during stress. Error bars: standard deviation from three biological replicates.

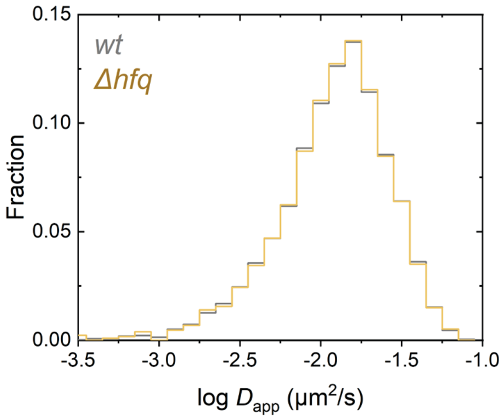


**Supplementary Figure 8.** Apparent diffusion coefficient distributions of RNE-mEos3.2 within BR-bodies in wild-type and Δ*hfq* *C. crescentus* cells during exponential phase. The similarity in diffusion profiles indicates that the absence of Hfq does not affect RNE-mEos3.2 mobility within BR-bodies under these conditions. 2,441 and 5,001 trajectories were analyzed for Δ*hfq* and *wt*, respectively.

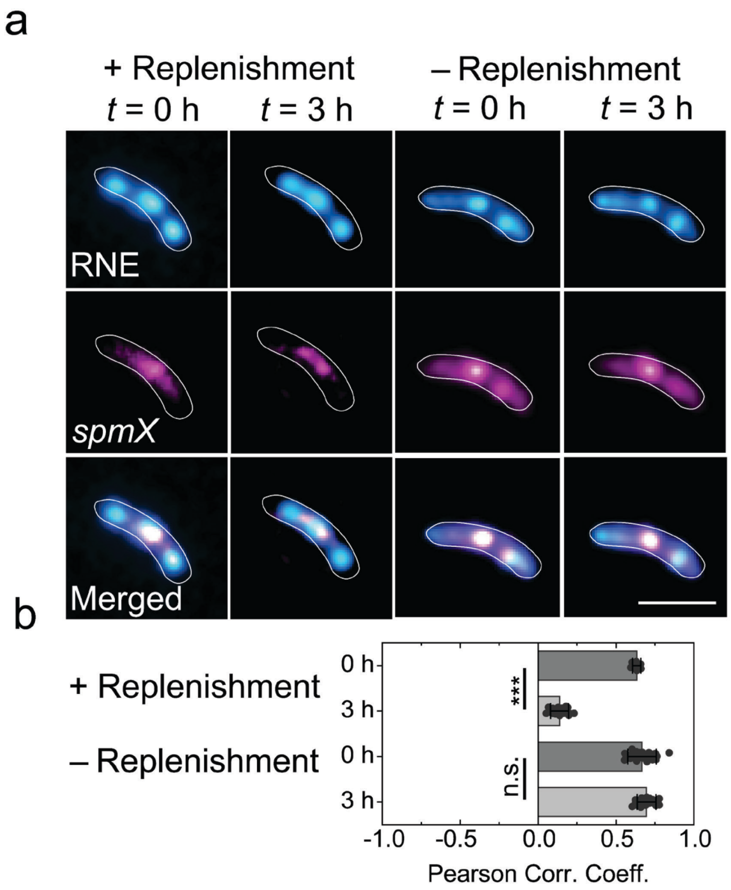


**Supplementary Figure 9.** Recruitment of mRNA into the BR-bodies after nutrient replenishment mimics mRNA recruitment in exponential phase. (a) Representative two-color images showing the recruitment profile of the *spmX* transcript (magenta) to BR-bodies (blue) in *C. crescentus* cells (white outlines) at time, *t*, following nutrient replenishment (left) and without nutrient replenishment (right). Right: Control. Scale bar: 2 μm. (b) Differences in recruitment were quantified by the Pearson correlation coefficients between channels. Error bars: standard deviation from 50 cells analyzed per phase. *** denotes $p < 0.001$. n.s. denotes $p > 0.05$.

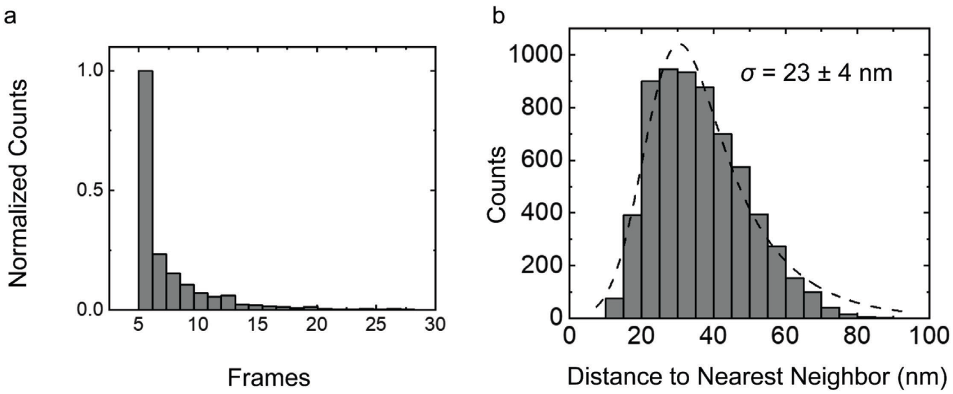


**Supplementary Figure 10**. Calculation of single-molecule RNE-mEos3.2 trajectory lengths and localization precisions in *C. crescentus cells*. *C. crescentus* cells expressing RNE-mEos3.2 were fixed, and single RNE-mEos3.2 molecules were photoconverted and then localized to obtain trajectories. (a) Distribution of trajectory lengths of RNE-mEos3.2 in fixed cells (imaging frame time: 20 ms). Trajectories lasting fewer than 4 frames were not used in our analysis. (b) Distribution of the nearest neighbor distances, $d$, in fixed cells based on repeated localizations of the same RNE-mEos3.2 molecule. The nearest neighbor distribution histogram, $p(d)$, was fit (dashed line) to Equation (2), as described in Ref. [91]. $\sigma$ is the localization precision, and $\omega$ is the Gaussian standard deviation of a short-range correction term centered at $d_c$. $a$, $A_1$, and $A_2$ are adjustable parameters.

$$p(d) = A_1\left(\frac{d}{\sqrt{2\sigma^2}}e^{\frac{-a^2}{4\sigma^2}}\right) + A_2\left(\frac{1}{\sqrt{2\pi\omega^2}}e^{\frac{-(d-d_C)^2}{2\omega^2}}\right) \qquad (2)$$

Supplementary Tables

**Supplementary Table 1. RNA half-lives measured by RT-qPCR.** Half-lives were determined for *ctrA* and 5S RNA in a Δ*hfq* background, for *tmRNA/ssrA*, *rne*, *ctrA*, and *spmX* during exponential, early stationary, and late stationary phases in a wild-type background.

| | | |
|---|---|---|
| *ctrA* (Δ*hfq*) | Exponential Phase | 0.8 ± 0.1 min |
| | Stationary Phase | 3466 ± 10 min |
| 5S (Δ*hfq*) | Exponential Phase | 24 ± 10 min |
| | Stationary Phase | 39 ± 10 min |
| tmRNA/*ssRA* | Exponential Phase | 13 ± 7 min |
| | Late Stationary Phase | 45 ± 12 min |
| *rne* (WT) | Exponential Phase | 0.48 ± 0.08 min |
| | Early Stationary Phase | 1.2 ± 0.4 min |
| | Late Stationary Phase | 9 ± 4 min |
| *ctrA* (wt) | Exponential Phase | 0.51 ± 0.06 min |
| | Early Stationary Phase | 0.9 ± 0.2 min |
| | Late Stationary Phase | 102 ± 20 min |
| *spmX* (wt) | Exponential Phase | 0.5 ± 0.1 min |
| | Early Stationary Phase | 0.81 ± 0.04 min |
| | Late Stationary Phase | 33 ± 17 min |

**Supplementary Table 2. Strains used in this work**

| STRAIN | SOURCE | IDENTIFIER |
|---|---|---|
| ***Caulobacter crescentus* NA1000** | Lucy Shapiro, Stanford University School of Medicine | N/A |
| **NA1000 *rne::rneΔCTD specR*** | This paper | *JS769* |
| **NA1000 *spmX::spmX-mEos3.2 specR*** | This paper | *JS770* |
| **NA1000 *acnA::acnA-PAmChy specR*** | This paper | *JS771* |
| **NA1000 *acnA::acnA-PAmChy rne::rne-eYFP specR gentR*** | This paper | *JS772* |
| **NA1000 *rhlB::rhlB-PAmChy SpecR*** | This paper | *JS773* |
| **NA1000 *rhlB::rhlB-PAmChy rne::rne-eYFP specR gentR*** | This paper | *JS774* |
| **NA1000 *rne::rne-eYFP gentR*** | This paper | *JS401* |
| **NA1000 *rne::rne-msfGFP gentR*** | Ref. [6] | *JS87* |
| **NA1000 *vanA::Ms2-DM-mChy rne::rne-msfGFP specR/gentR*** | Ref. [2] | *JS285* |
| **NA1000 *spmX::ms2 96 array vanA::Ms2-DM-mChy rne::rne-msfGFP specR/kanR/gentR*** | Ref. [2] | *JS283* |
| **NA1000 *rsaA::ms2 96 array vanA::Ms2-DM-mChy rne::rne-msfGFP specR/kanR/gentR*** | Ref. [2] | *JS287* |
| **NA1000 *rne::rneΔDBS SpecR*** | Ref. [3] | *JS801* |

| | | |
|---|---|---|
| **NA1000 *rne::rne-ΔDBS-mEos3.2 SpecR*** | This paper | *JS804* |
| **NA1000 *hfq::tet rne::rne-mEos3.2C-1 specR/tetR*** | This paper | *JS805* |
| **NA1000 *rne::rneΔCTD Kan$^R$*** | This paper | *JS806* |
| **NA1000 *rne::rneΔCTD spmX::spmX-mEos3.2 specR Kan$^R$*** | This paper | *JS807* |
| **NA1000 *hfq::tet tetR*** | Ref. [4] | *CJW5477* |
| **NA1000 *hfq::spec SpecR*** | Gift of Lucy Shapiro | *JS263* |
| **NA1000 *rne::rne-mEos3.2C-1 SpecR*** | This paper | *JS775* |
| **NA1000 *rne::rne-ΔCTD-mEos3.2 SpecR*** | This paper | *JS776* |
| **NA1000 *ctrA::ms2 48 array kanR*** | This paper | *JS777* |
| **NA1000 *spmX::ms2 48 array kanR*** | This paper | *JS778* |
| **NA1000 *rsaA::ms2 48 array kanR*** | This paper | *JS779* |
| **NA1000 *ctrA::ms2 48 array vanA::Ms2-DM-mChy rne::rne-msfGFP specR/kanR/gentR*** | This paper | *JS780* |
| **NA1000 *spmX::ms2 48 array vanA::Ms2-DM-mChy rne::rne-msfGFP specR/kanR/gentR*** | This paper | *JS789* |
| **NA1000 rsaA::ms2 48 array vanA::Ms2-DM-mChy rne::rne-msfGFP specR/kanR/gentR** | This paper | *JS790* |
| **Agrobacterium tumefaciens C56 *rne::rne-YFP gentR*** | Ref. [6] | *JS5* |
| **Sinorhizobium meliloti *rne::rne-YFP gentR*** | Ref. [6] | *JS9* |
| ***E. coli* DH10B** | Thermofisher scientific | EC0113 |
| ***E. coli* TOP10** | Thermofisher scientific | *C404010* |

**Supplementary Table 3. Chemicals, peptides, and recombinant proteins used in this work**

| ***ITEM*** | ***SOURCE*** | ***IDENTIFIER*** |
|---|---|---|
| Gibson Master Mix | New England BioLabs Inc. | E2611S |
| L-Azidohomoalanine (AHA) | ClickChemistryTools | 1066-25 |
| Paraformaldehyde, 96% | ThermoFisher Scientific | A11313.22 |
| Cy5 Alkyne | ClickChemistryTools | TA116-1 |
| Iodoacetamide | Sigma-Aldrich | 144-48-9 |
| THPTA | Click Chemistry Tools | 1010-100 |
| Copper (II) sulfate pentahydrate | Sigma-Aldrich | 7758-99-8 |
| TRizol Reagent | Ambion | 15596018 |
| Spectinomycin | Sigma-Aldrich | S6501-25G |
| Rifampicin | Sigma-Aldrich | R7382-1G |
| Kanamycin | Sigma-Aldrich | K1377-5G |

| | | |
|---|---|---|
| Chloramphenicol | Sigma-Aldrich | C0378-25G |
| Nalidixic Acid | Sigma-Aldrich | N8878-5G |
| Sucrose | Sigma-Aldrich | S5016-25G |
| Phusion DNA polymerase | Thermo Scientific | F-530L |
| RNAprotect Bacterial Reagent | QIAGEN | 76506 |
| Luna® Universal One-Step RT-qPCR Kit | NEB | E3005L |
| Qubit RNA HS Assay Kit | Thermo Fischer Scientific | Q32851 |
| Chloroform | Thermofisher scientific | AC423550010 |
| Glycogen, RNA grade | Thermofisher scientific | RO551 |
| Sodium Ascorbate | Thermofisher scientific | 134-03-2 |
| 2-PROPANOL, ANHYDROUS | Sigma-Aldrich | 67-63-0 |
| ETHANOL-D6 (D, 99%), ANHYD. | Sigma-Aldrich | 1516-08-1 |
| EDTA (0.5 M), pH 8.0, RNase-free | Sigma-Aldrich | AM9261 |
| Tris (1 M), pH 7.0, RNase-free | Thermofisher scientific | AM9851 |
| Agar | Thermofisher scientific | DF0001-17-0 |
| Bactopeptone | Fisherchemicals | 211677 |
| Yeast extract | Thermofisher scientific | 92144-500G-F |
| UltraPure™ Ethidium Bromide, 10 mg/mL | Sigma-Aldrich | 15585011 |
| Thermo Scientific™ TriTrack DNA Loading Dye (6X) | Thermofisher scientific | FERR1161 |
| Magnesium sulfate | Thermofisher scientific | M7506-1KG |
| Calcium chloride (97%) | Sigma-Aldrich | 746495-500G |
| Potassium phosphate dibasic anhydrous | Sigma-Aldrich | P288-500 |
| Ammonium Chloride (99.5%) | Fisherchemicals | A9434-1KG |
| Sodium phosphate dibasic | Sigma-Aldrich | S7907-1KG |
| Luria Broth Base | Sigma-Aldrich | 12795084 |
| Iron (II) sulfate heptahydrate | Thermofisher scientific | F8633-250G |
| FD Dpn1 | Sigma-Aldrich | ER1701 |
| T4 DNA Ligase | Thermofisher scientific | EL0011 |
| Agarose | Thermofisher scientific | A7705 |
| RNAprotect Bacteria Reagent | Qiagen | 76506 |
| Gentamycin Sulfate | Sigma-Aldrich | 345814-1GM |
| Carbonyl Cyanide *m*-Chlorophenylhydrazone | Sigma-Aldrich | 555-60-2 |
| Glucose (99.5%) | Sigma-Aldrich | G8270-1KG |
| FD EcoR1 | Thermofisher scientific | FD0274 |
| FD Nde1 | Thermofisher scientific | FD0585 |
| FD Kpn1 | Thermofisher scientific | FD0524 |
| Paraffin wax | Sigma-Aldrich | 327204-1KG |
| Oxytetracycline, 96% | Thermofisher scientific | J62427-09 |
| T4 Polynucleotide Kinase (T4 PNK) | New England Biolabs | M0201S |

**Supplementary Table 4. Primers used for strain construction**

| ITEM | SEQUENCE |
|---|---|
| ***Primer Ups F for JS769 rne::rneΔCTD specR*** | atctggatccagcgcctgaagagcgtcg |
| ***Primer Ups R for JS769 rne::rneΔCTD specR*** | gttgtcattcttagtcatcgaccacgaccg |
| ***Primer Down F for JS769 rne::rneΔCTD specR*** | cgatgactaagaatgacaacggcccgcg |
| ***Primer Down R for JS769 rne::rneΔCTD specR*** | tagcgaattcgaggaagccctcaagcttg |
| ***Primer pNTPS F for JS769 rne::rneΔCTD specR*** | gggcttcctcgaattcgctagcttcggc |
| ***Primer Spec F for JS769 rne::rneΔCTD specR*** | cgatgactaagatcttttctacggggtc |
| ***Primer Spec R for JS769 rne::rneΔCTD specR*** | gttgtcattcctagatgccgactaccttg |
| ***Primer ΔCTD F for JS769 rne::rneΔCTD specR*** | cggcatctaggaatgacaacggcccgcg |
| ***Primer ΔCTD R for JS769 rne::rneΔCTD specR*** | agaaaagatcttagtcatcgaccacgaccg |
| ***Primer pspmX F for JS283 spmX::ms2 96 array vanA::Ms2-DM-mChy rne::rne-msfGFP specR/kanR/gentR*** | ataaggtacccggcgccggtgctgttcg |
| ***Primer pspmX R for JS283 spmX::ms2 96 array vanA::Ms2-DM-mChy rne::rne-msfGFP specR/kanR/gentR*** | atattagaattcctactcttcgtcgctcacatcggggc |
| ***IDT G-block for JS775 rne::rne-mEos3.2C-1 SpecR*** | ccatggagggcgacaccttctataacaaggtgcgcttctat<br>ggcaccaacttcccggccaatggccccgtgatgcagaag<br>aagaccctgaagtgggaacccagcaccgaaaagatgtac<br>gtgcgcgacggcgtcctgacgggcgacatcgagatggcgc<br>tgctcctggagggcaacgcgcactatcgttgcgatttccgc<br>accacctacaaggcgaaggagaagggcgtgaagctcccg<br>ggcgcccatttcgtcgaccattgcatcgaaatcctgtccca<br>cgataaggattataataaggtgaagctgtacgagcacgcc<br>gtggcgcattccgggctgccggataacgcccgccgttaag<br>ctagcaaagaattcgaacgttacgcgtcaccggtcggcca<br>ccatgtcggcgatcaagccggacatgaagatcaagctgcg<br>catggagggcaatgtgaacgggcaccacttcgtgatcgac<br>ggcgacggcaccgggaagccgttcgagggcaagcagtcg<br>atggacctggaggtcaaggagggcgggccgctccccttcg<br>cgttcgacatcctcacgaccgccttccactacggcaaccg<br>ggtgtttgccaagtatccggacaacatccaggactatttca<br>agcagtcgttcccgaagggctatagctgggagcgctcgctg<br>accttcgaggacggcggtatctgcaacgcccgcaacgac<br>atcaccatggagggcgacaccttctataacaaggtgcgctt<br>ctatggcaccaacttcccggccaatggccccgtgatgcag<br>aagaagaccctgaagtgggaacccagcaccgaaaagat<br>gtacgtgcgcgacggcgtcctgacgggcgacatcgagatg<br>gcgctgctcctggagggcaacgcgcactatcgttgcgattt<br>ccgcaccacctacaaggcgaaggagaagggcgtgaagct |

| | |
|---|---|
| | cccgggcgcccatttcgtcgaccattgcatcgaaatcctgt cccacgataaggattataataaggtgaagctgtacgagca cgccgtggcgcattccgggctgccggataacgcccgccgt taagctagcaaa |
| ***Primer HY118F for JS776 rne::rne-ΔCTD-mEos3.2 SpecR*** | attgcatatgccacgcgcgagcgcaaca |
| ***Primer HY118R for JS776 rne::rne-ΔCTD-mEos3.2 SpecR*** | cgttcgaattgtcatcgaccacgaccgacacg |
| ***Primer HY117F for JS776 rne::rne-ΔCTD-mEos3.2*** | ggtcgatgacaattcgaacgttacgcgtc |
| ***Primer HY115R for JS776 rne::rne-ΔCTD-mEos3.2*** | tcgcgcgtggcatatgcaattggacgtctc |
| ***Primer HY116F for JS770 spmX::spmX-mEos3.2 specR*** | attgcatatgacaagcttctgaacaggc |
| ***Primer HY116R for JS770 spmX::spmX-mEos3.2 specR*** | cgttcgaattctcttcgtcgctcacatc |
| ***Primer HY115F for JS770 spmX::spmX-mEos3.2 specR*** | cgacgaagagaattcgaacgttacgcgtc |
| ***Primer HY115R for JS770 spmX::spmX-mEos3.2 specR*** | agaagcttgtcatatgcaattggacgtctc |
| ***Primer HY120F for JS804 rne::rne-ΔDBS-mEos3.2 SpecR*** | attgcatatgagcctgcacgcgggcgac |
| ***Primer HY120R for JS804 rne::rne-ΔDBS-mEos3.2 SpecR*** | cgttcgaattcggcgcggtgatctcgttcg |
| ***Primer HY119F for JS804 rne::rne-ΔDBS-mEos3.2 SpecR*** | caccgcgccgaattcgaacgttacgcgtc |
| ***Primer HY119R for JS804 rne::rne-ΔDBS-mEos3.2 SpecR*** | cgtgcaggctcatatgcaattggacgtctc |
| ***Primer HY1'F for JS806 NA1000 rne::rneΔCTD KanR*** | ggtcgatgactaagctagctgcagcccg |
| ***Primer HY1'R for JS806 NA1000 rne::rneΔCTD KanR*** | agctagcttagtcatcgaccacgaccgac |
| ***Primer HY24F for JS771 acnA::acnA-PAmChy specR*** | ggtcggccacatggtgagcaagggcgag |
| ***Primer HY24R for JS771 acnA::acnA-PAmChy specR*** | ctgcagctagttacttgtacagctcgtccatg |
| ***Primer HY23F for JS771 acnA::acnA-PAmChy specR*** | gtacaagtaactagctgcagcccggggg |
| ***Primer HY23R for JS771 acnA::acnA-PAmChy specR*** | tgctcaccatgtggccgaccggtgacgc |
| ***Primer HY47F for JS771 acnA::acnA-PAmChy*** | ccttaattaacaaccgcatcaccccgga |
| ***Primer HY47R for JS771 acnA::acnA-PAmChy*** | cgaattctccgtcggccttggccaggtt |
| ***Primer HY48F for JS771 acnA::acnA-PAmChy*** | caaggccgacggagaattcgaacgttacg |
| ***Primer HY48R for JS771 acnA::acnA-PAmChy*** | gatgcggttgttaattaaggcgcctgcag |

| | |
|---|---|
| ***Primer HY49F for JS773 rhlB::rhlB-PAmChy SpecR*** | ccttaattaacgcaccatgccgacgactatgtccaccg |
| ***Primer HY49R for JS773 rhlB::rhlB-PAmChy SpecR*** | cgaattctcccttcgcgccgcgcggcgg |
| ***Primer HY50F for JS773 rhlB::rhlB-PAmChy SpecR*** | cggcgcgaagggagaattcgaacgttacg |
| ***Primer HY50R for JS773 rhlB::rhlB-PAmChy SpecR*** | gcatggtgcgttaattaaggcgcctgcag |
| ***IDT G-block for JS777 ctrA::ms2 48 array kanR*** | caattggatccaatcttgacgtccgtttgattacgatcaaga<br>ttggatccaataggtcgccgaaccacgatgcgaggaaacg<br>catatgtcgaagaagcctgcaggcgccttaattaatatgca<br>tggtaccttaagatctcgagctccggagaattcgaacgtta<br>cgcgtcaccggtcggccaccatggtgagcaagggcgagg<br>agctgttcaccggggtggtgcccatcctggtcgagctggac<br>ggcgacgtaaacggccacaagttcagcgtgtccggcgagg<br>gcgagggcgatgccacctacggcaagctgaccctgaagtt<br>catctgcaccaccggcaagctgcccgtgccctggcccac<br>cctcgtgaccaccctgacctggggcgtgcagtgcttcagc<br>cgctaccccgaccacatgaagcagcacgacttcttcaagt<br>ccgccatgcccgaaggctacgtccaggagcgcaccatctt<br>cttcaaggacgacggcaactacaagacccgcgccgaggt<br>gaagttcgagggcgacaccctggtgaaccgcatcgagctg<br>aagggcatcgacttcaaggaggacggcaacatcctgggg<br>cacaagctggagtacaactacatcagccacaacgtctata<br>tcaccgccgacaagcagaagaacggcatcaaggccaac<br>ttcaagatccgccacaacatcgaggacggcagcgtgcag<br>ctcgccgaccactaccagcagaacacccccatcggcga<br>cggccccgtgctgctgcccgacaaccactacctgagcac<br>ccagtccgccctgagcaaagaccccaacgagaagcgcg<br>atcacatggtcctgctggagttcgtgaccgccgccgggatc<br>actctcggcatggacgagctgtacaagtaatctagaagctc<br>ggtattcctgagcttgttaacaccgatccgcatggcaccgtt<br>tcggtgtgcgtaggagggtgattgcgcggtagtcctgcgcat<br>agtggaaataccttagccagccctttaaaatgcggtatgtc<br>acaaggtacagggcgtattcgtccctgtacgaagagcacc<br>tttcggtagcgtctaactacacctcctgtcctcgaatcgcag<br>ggtattcctctgcgtaggtgtcggagtgtgacgcatagttcca<br>accgtcacagggacccttgataatggggtattcctccattg<br>acacacgatacgcttcacgtaaatatacggacagtccgtt<br>cttcataacagaagcgtaatcgtcttctctcttctttgccagc<br>cgcggtcatcgccatgcaagaaatatcgcagcacttggtc<br>ggtagtcctgaccacaaggctgacgcctagtctttacagct<br>gcctactgcccttatcggtgcaacatgcggtaatcctgcat<br>gctaacttgttcatactatataggatgtgtttgctgccgcaga<br>aataaacccacgcggtaatcctgcgtgcgttctagtgtcttg<br>cgtgagcggtagcctggcatcagccatgccccggatgtgg<br>ggtagtcctccacaaagggagaacatccatatatagtggg<br>gagtctctcgtataagtgcttatgcacagggtattcctctgtg |

tattgagtatgaatcgactccctggaagaaagagtatatttc
tccttgccgacgcggtagtcctgcgtcattgccgccaggtg
gcgacgctcggttgatgcctgaaagcatttcaccgaagctc
ggtattcctgagctaagcttgaactgttccccaagggtcggt
ctacaggctgtggcgtcacacatgcgcggtagtcctgcgca
ccagcttggatgtcctcgcgactctttagatgtcttggagca
ggagcaggcagggcgtattcgtccctgcgagaatagtgctt
gaggttcgaaacaggacatacatcttgagagaaagccgca
gggtattcctctgcgcggcgagacggcctacgcagactaa
cgtataatacatcaaacatgtcgtgaatggggtattcctcca
ttcctaataggattgtgtaagaaggcgaggtgcagtggctgc
taaactctatagaagcgtaatcgtcttctcaatcgagtataa
ctatttgaatgcgcgctactcgcaggcatagccaatgtggtc
ggtagtcctgaccatggttaaccaggcaccgccctgtcga
atatccgggtcacaataggctttgcatgcggtaatcctgcat
gtaatagctcgtatgtggtgtccggtcccggttaacaagctt
atgacgtaccacgcggtaatcctgcgtgatctgaatgtgaa
tcgcgtagcatgtgcggggtgcaacgttcgagttcggtgtgg
ggtagtcctccacaacttcaccgctagatcaccattggtct
cagagccataatagtcctgctaacacagggtattcctctgt
ggccatagggtcatgtgcgtgatccgcgtcgtccgggtatag
agttaatgagacgcggtagtcctgcgtccccgcatgcatgt
gccggaatgctccgctacccactcggtttataaattgagct
cggtattcctgagctgatctaagagaaacagcgcgatcatc
tgacaccaacttgatactgatcgctgcgcggtagtcctgcg
cacggctccaatcacaccagccatgtagtgagtatactcc
tcatgcgtatcccagggcgtattcgtccctgactctgcgtac
atcggatccccgctccgaccacactccagaccacggagc
cgcagggtattcctctgcgaatggggtattcctccattgtgc
gaaaaactgcgtggtagacatcctgacgaaattcagggca
atagcgagaagcgtaatcgtcttcttcgttgggcgcgcccg
cgaccagatgctatgcaacttgcagctgagaatctggtcgg
tagtcctgaccagggtcgtcagctgatcgacaagctggttg
gcagtgttatcgatgtgcatgcatgcggtaatcctgcatgtg
cctgcctttaatggtgcgattgtgttgttccgccatgagatgg
gagtacacgcggtaatcctgcgtgcacttgccgacattcta
gtctctataaggcaactgtgtccagaggcgttatgtggggta
gtcctccacacgttagatagttctaggctcagtttgagggat
ctcacatgtctagctccccacagggtattcctctgtggtgga
taattcgctactctcactgcttctgtaaacgaaaagagatag
aatgacgcggtagtcctgcgtcatggaggaggttatcgtag
cgtattcatcttatctacaatcaagaacgcgagctcggtatt
cctgagctcagcgggctctgatagaattgcatgtctttgggc
actgttaagatatcgttgcgcggtagtcctgcgcacacatga
tagtgcctctggatccttaggtctcgaccaccatgtgactaa
ccagggcgtattcgtccctgctcattgttctcggtggacgttg
gttgaaacagggctttgtagagtcatccgcagggtattcctc
tgcgttccctccagggacattatcgggaccattagtatggcg

| | |
|---|---|
| | caaattctgataaatggggtattcctccatttagtcctacga cacttccgatgtacagaagggccaccacattcatagacga gaagcgtaatcgtcttctcacgccaggttcccccaagttag gggactgcgcgggtggcccccccccgctggtcggtagtcc tgaccaagtacaacccactaaccagaacaccgctccctg gggatctttgcaatgatcatgcggtaatcctgcatgcgcata agggaaacacgcaaaaccacgaggtcgacgggagccgc aacacgcacgcggtaatcctgcgtgtcacaccaaccata gggaaacgacgcgactggttgcttcgacagcctgcgtgtgg ggtagtcctccacagagatccggcgcgagacatggtgcac catgcagaggcgtagagagctccgcacagggtattcctctg tgacaaactaacaggttatctaggaagattcggccaaccgt cggtaatactagacgcggtagtcctgcgtcgctagcataaa acgaaaggctcagtcgaaagactgggcctttcgttttatctg ttgtttgtcggtgaacgctctcctgagtaggacaaatccgcc gggagcggatttgaacgttgcgaagcaacggcccggaggg tggcgggcaggacgcccgccataaactgccaggcatcaa attaagcagaaggccatcctgacggatggcctttcgatcg |
| ***Primer HY55F for JS777 ctrA::ms2 48 array kanR*** | gccaccatggtctagaagctcggtattcc |
| ***Primer HY55R for JS777 ctrA::ms2 48 array kanR*** | gaacctcaagcactattctcgcagggac |
| ***Primer HY53F for JS777 ctrA::ms2 48 array kanR*** | ctttcgatcggctagctgcagcccgggg |
| ***Primer HY53R for JS777 ctrA::ms2 48 array kanR*** | agcttctagaccatggtggccgaccggtg |
| ***Primer HY63F for JS777 ctrA::ms2 48 array kanR*** | gcatggtacctcgtcggaaatcgacacc |
| ***Primer HY63R for JS777 ctrA::ms2 48 array kanR*** | ttcgaattcttcaggcggcgttaacctg |
| ***Primer HY64F for JS777 ctrA::ms2 48 array kanR*** | cgccgcctgaagaattcgaacgttacgc |
| ***Primer HY64R for JS777 ctrA::ms2 48 array kanR*** | tttccgacgaggtaccatgcatattaattaag |
| ***Primer HY57F for JS778 spmX::ms2 48 array kanR*** | gcatggtaccacaagcttctgaacaggc |
| ***Primer HY57R for JS778 spmX::ms2 48 array kanR*** | ttcgaattctctactcttcgtcgctcac |
| ***Primer HY58F for JS778 spmX::ms2 48 array kanR*** | cgaagagtagagaattcgaacgttacgc |
| ***Primer HY58R for JS778 spmX::ms2 48 array kanR*** | agaagcttgtggtaccatgcatattaattaag |
| ***Primer HY61F for JS779 spmX::ms2 48 array kanR*** | gcatggtacctcgacgccagcgccgtca |
| ***Primer HY61F for JS779 spmX::ms2 48 array kanR*** | ttcgaattctttaggcgagcgtcaggacttcg |

| | |
|---|---|
| ***Primer HY62F for JS779 spmX::ms2 48 array kanR*** | gctcgcctaaagaattcgaacgttacgc |
| ***Primer HY62F for JS779 spmX::ms2 48 array kanR*** | ctggcgtcgaggtaccatgcatattaattaag |

**Supplementary Table 5. mRNA FISH probes used in this work.**

| *ctrA* probes | *spmX* probes | *rsaA* probes |
|---|---|---|
| taatggtgaatgtttcccgt | agtttgagacagacgcggat | gtgtacgcagtcaccaactg |
| cactttgccggagagttaat | gtttcatcgcctagattgac | tctgggtttgagtcgcgtac |
| tggcgtattcgaacgtcttt | tcttgatgagatcgacggcg | taggtctggatggcaacagc |
| catcctcgatcaacagtacg | ttctgacggtacccttcgaa | ggcaacgccggtgaagaact |
| ttcagacttcagcatcagtt | cgtccttttccgagacagag | tggtcgagtcgaccaggaag |
| tccgtcgtatagacgttgaa | tgagatcatagagcagcagc | aacttcgagtagtacgcgtc |
| tagatcttgcccagatcgac | gtgtgctcgttcaccgaatg | gatgaagcggttttcctgag |
| gagcaggataagatcgtagt | tcgaactggttctggttcag | tggccaggttgatcgagaag |
| tcatgtccggaagattgagg | tattgaaggcgaagcacacc | cgtacgaaacgcccgtgtag |
| agggtgcgcagaacatcgat | gagcggaggaagttgtcgag | cgatgatcttgtcataggcg |
| atcatgatgggcgtgttgat | aaccagtgcatcgatgacga | cggtcaggtagtcgatgttg |
| ttgaccttggtgtcgatttc | cgtgaggaacagagtcttct | ttgacggccagatcgatgtc |
| cttggtcatgtagtcgtcgg | tcatagtcgaccttgggacg | ttcaggatggtgccgatcag |
| atcatttcgtccttgtggaa | cggaacggcgtgaactcgac | cgacaggtcgttgatcatcg |
| tgacccttcgaacgacggac | ctgttcagaagcttgtggcg | gacggataggcggtgaacag |
| cggtcttgatgaccgactgg | gattaaagaccgacgagccg | tggtcagcgagagggtcgaa |
| acaggtggttcaggaacatt | tgatcagcaggctggtgttg | aacgaacgtgtcgttgttgg |
| gtcgatgatcttcagttccg | acggtcaggatccagaggac | tcacgttcatcgtttcgatg |
| tgcgcagcttgcagatgaag | tagccgaagagcttgacctg | aagacgtgttcagggtgatc |
| cagaccgtctcaatgtggtg | cgagaccacgaagcagatcg | ttgcgtcaacgtggtgttca |
| aacgactcaggcggcgttaa | cgaggcaagaaacctactct | agagtcggtgatcgtcacag |
| tgggttctacagatacgctg | | caggttgacggtcgtcagag |
| agaaaacgcccaccttttgg | | gtcagaccattgacgttcag |
| | | ggtgaaaccatcgtcagcag |
| | | tcgaaccagcgatgttgatg |
| | | cgagatgttcagggtcgtcg |
| | | tgtgcgaggtgatcgtgacg |
| | | ttgacgttggccaccagaac |
| | | gtcagcgctgaacgacgaac |
| | | cgacgcggagggtttcgaag |
| | | caacgttggtgaaggtcgtc |
| | | agaacggtcaggccgacatt |

| | | |
|---|---|---|
| | | gacagggtcaggttgaacac |
| | | cgtcaccacgatcgacttgg |
| | | ggtgttggtcaggttcagac |
| | | acacgaaggtcacagccgag |
| | | gcggatcgtgacgacttcac |
| | | caccgatgatggtgtcattg |
| | | gatatcgaagatatccgcgc |
| | | gtcggtgatcgtcacgaaag |
| | | ttcgtcgagatgccgacgag |
| | | aactggaaccacttggcaac |
| | | gtcaacgacgacataggtgt |
| | | agaccggtcagcttgatcac |
| | | tcaggacttcggtggcgaag |

### Supplementary Movie Captions

#### Supplementary Movie 1.

Representative fluorescence movie of BR-bodies in an exponential-phase culture of *C. crescentus* expressing RNE-mEos3.2 (shown in blue). BR-bodies are short-lived in exponential phase. Field of view: 19 × 17 μm.

#### Supplementary Movie 2.

Representative fluorescence movie of BR-bodies in a stationary-phase culture of *C. crescentus* expressing RNE-mEos3.2 (shown in blue). BR-bodies are long-lived in stationary phase. Field of view: 22 × 20 μm.

**Supplementary References**


1. Thanbichler, M., Iniesta, A. A. & Shapiro, L. A comprehensive set of plasmids for vanillate- and xylose-inducible gene expression in Caulobacter crescentus. *Nucleic Acids Res.* **35,** e137 (2007).
2. Al-Husini, N., Tomares, D. T., Pfaffenberger, Z. J., Muthunayake, N. S., Samad, M. A., Zuo, T., Bitar, O., Aretakis, J. R., Bharmal, M.-H. M., Gega, A., Biteen, J. S., Childers, W. S. & Schrader, J. M. BR-Bodies Provide Selectively Permeable Condensates that Stimulate mRNA Decay and Prevent Release of Decay Intermediates. *Mol. Cell* **78,** 670-682.e8 (2020).
3. Passos, C., Tomares, D., Yassine, H., Schnorr, W., Hunter, H., Wolfe-Feichter, H. K., Velier, J., Dzurik, K. G., Grillo, J., Gega, A., Saxena, S., Schrader, J. & Childers, W. S. BR-Bodies Facilitate Adaptive Responses and Survival During Copper Stress in Caulobacter crescentus. 2025.03.11.642215 Preprint at https://doi.org/10.1101/2025.03.11.642215 (2025)
4. Irnov, I., Wang, Z., Jannetty, N. D., Bustamante, J. A., Rhee, K. Y. & Jacobs-Wagner, C. Crosstalk between the tricarboxylic acid cycle and peptidoglycan synthesis in Caulobacter crescentus through the homeostatic control of α-ketoglutarate. *PLOS Genet.* **13,** e1006978 (2017).
5. Yassine, H. & Schrader, J. M. APEX2 proximity labeling of RNA in bacteria. 2024.09.18.612050 Preprint at https://doi.org/10.1101/2024.09.18.612050 (2024)
6. Al-Husini, N., Tomares, D. T., Bitar, O., Childers, W. S. & Schrader, J. M. α-Proteobacterial RNA Degradosomes Assemble Liquid-Liquid Phase-Separated RNP Bodies. *Mol. Cell* **71,** 1027-1039.e14 (2018).
7. Schrader, J. M. & Shapiro, L. Synchronization of Caulobacter Crescentus for Investigation of the Bacterial Cell Cycle. *J. Vis. Exp. JoVE* e52633 (2015). doi:10.3791/52633
8. Ducret, A., Quardokus, E. M. & Brun, Y. V. MicrobeJ, a tool for high throughput bacterial cell detection and quantitative analysis. *Nat. Microbiol.* **1,** 1–7 (2016).

9. Schindelin, J., Arganda-Carreras, I., Frise, E., Kaynig, V., Longair, M., Pietzsch, T., Preibisch, S., Rueden, C., Saalfeld, S., Schmid, B., Tinevez, J.-Y., White, D. J., Hartenstein, V., Eliceiri, K., Tomancak, P. & Cardona, A. Fiji: an open-source platform for biological-image analysis. *Nat. Methods* **9,** 676–682 (2012).

10. Isaacoff, B. P., Li, Y., Lee, S. A. & Biteen, J. S. SMALL-LABS: Measuring Single-Molecule Intensity and Position in Obscuring Backgrounds. *Biophys. J.* **116,** 975–982 (2019).

11. Endesfelder, U., Malkusch, S., Fricke, F. & Heilemann, M. A simple method to estimate the average localization precision of a single-molecule localization microscopy experiment. *Histochem. Cell Biol.* **141,** 629–638 (2014).

12. Ester, M., Kriegel, H.-P., Sander, J. & Xu, X. A density-based algorithm for discovering clusters in large spatial databases with noise. in *Proc. Second Int. Conf. Knowl. Discov. Data Min.* 226–231 (AAAI Press, 1996).

13. Miura, K. Bleach correction ImageJ plugin for compensating the photobleaching of time-lapse sequences. Preprint at https://doi.org/10.12688/f1000research.27171.1 (2020)

14. Nandana, V., Al-Husini, N., Vaishnav, A., Dilrangi, K. H. & Schrader, J. M. Caulobacter crescentus RNase E condensation contributes to autoregulation and fitness. *Mol. Biol. Cell* **35,** ar104 (2024).

15. Nandana, V., Rathnayaka-Mudiyanselage, I. W., Muthunayake, N. S., Hatami, A., Mousseau, C. B., Ortiz-Rodríguez, L. A., Vaishnav, J., Collins, M., Gega, A., Mallikaarachchi, K. S., Yassine, H., Ghosh, A., Biteen, J. S., Zhu, Y., Champion, M. M., Childers, W. S. & Schrader, J. M. The BR-body proteome contains a complex network of protein-protein and protein-RNA interactions. *Cell Rep.* **42,** 113229 (2023).
